# Benchmarking four large language models on generating and selecting hypotheses that match published findings in 50 studies of ageing biology

**DOI:** 10.64898/2026.11.17.767323

**Authors:** Yuriy Poteshkin, Evgenii Udodov, Mikhail Batin

**Affiliations:** Open Longevity, Sherman Oaks, California, USA

**Keywords:** large language models, scientific hypothesis generation, ageing biology, benchmark, forecasting experimental confirmation, predictive value, model agreement, hypothesis selection

## Abstract

**Objective:** We asked how often hypotheses generated by large language models for 50 research tasks matched the withheld finding of the published study, and whether model forecasts could identify the matching hypotheses.

**Methods:** We built tasks from 50 peer-reviewed studies in ageing biology published after the models’ stated training cutoffs; each task gave the background and a research question and withheld the study’s finding (the reference finding). GPT-6 Astra, Grok 4.5, Claude Opus 5.5, and Gemini 4 Argon each generated five hypotheses per task by three methods: a single request, a multistep pipeline, and the Omega Point method without literature search (2997 hypotheses). A hypothesis matched the reference finding if at least two of three judge models from other developers scored it at least 3 (primary threshold) or at least 4 (strict threshold) on a 0 to 5 scale; refusals counted as non-matches. A selection rule, chosen from 112 variants evaluated on the Grok 4.5 hypotheses, kept only the hypotheses for which all four predictor models forecast experimental confirmation; the predictors never saw the published finding. The controls were recorded random answers and each generator’s first-listed hypothesis. Confidence intervals (95% CIs) were calculated by cluster bootstrap over tasks.

**Results:** Of the hypotheses generated by a single request, 13.2% to 18.0% matched the reference finding, compared with 11.2% to 18.8% for the multistep pipeline and 2.8% to 7.6% for the Omega Point method. With a single request, the hypothesis listed first matched in 37.5% of cases (75/200), and the hypotheses in positions 2 to 5 matched in 10.1% (81/800). Among the hypotheses selected unanimously, the match rate was 73.7% (14/19) for Grok 4.5, 47.4% (9/19) for Gemini 4 Argon, and 63.6% (14/22) for Claude Opus 5.5, compared with 12.1% to 17.9% for the random control and 37.5% to 40.0% for the first-listed hypotheses. The selected lists captured 27.3% to 35.0% of all matching hypotheses. Pooled across the three sets, the match rate among selected hypotheses exceeded that among first-listed hypotheses by 23.2 percentage points (pp; 95% CI [13.0, 32.6]). When Gemini 4 Argon re-extracted the reference findings, the match rate among selected hypotheses was 59.6% (34/57), compared with 61.4% (35/57) against the original reference findings for the same tasks. In a manual audit with the generator, judge scores, and selection status hidden, our classifications agreed with the judges’ at the primary threshold in 81/100 cases (Cohen’s kappa 0.62). Agreement was 89.7% after weighting the sample strata by their size in the corpus. We classified 19 of 30 sampled selected hypotheses as matches, and the judges classified 20 of 30.

**Conclusions:** In the evaluated sets, unanimous selection by the four predictors increased the match rate among selected hypotheses; the resulting match rate depended on the matching threshold. Generators listed a matching hypothesis first more often than in later positions, and 50 of the 60 selected hypotheses were listed first. The advantage over the first-listed hypotheses and over the best single predictor at equal list length was statistically significant for the Grok and Opus hypotheses but not for the Gemini hypotheses, and the pairwise differences of the advantage between generators were not statistically significant (*p* ≥ 0.16 after Holm correction). We selected the rule using the Grok hypotheses and evaluated it on the Gemini and Opus hypotheses for the same 50 tasks; it has not been tested on new tasks.

## 1. Introduction

### 1.1. Generating and selecting hypotheses for testing

Language models propose more hypotheses than a laboratory can test. We therefore measured not only the overall match rate but also whether models could identify which of the generated hypotheses matched the published finding.

We addressed two questions. First, how often do models, given the background and the research question, propose hypotheses that match the findings of 50 published experimental studies? Second, do the hypotheses that models forecast to be confirmed, without seeing the published finding, actually match that finding? Both “yes” and “no” forecasts were also evaluated on a separate set of confirmed and refuted statements.

### 1.2. Existing approaches to evaluating hypotheses

Published benchmarks evaluate hypotheses in three ways: judges assess the text of the hypothesis, the hypothesis is compared with an already published finding, or the proposed study is carried out in the laboratory.

In LiveIdeaBench, models receive a keyword and propose ideas, which other models rate for originality, feasibility, and other properties 1. This approach can evaluate ideas that have not yet been tested experimentally, but the score reflects the judges’ opinion rather than experimental confirmation.

In MOOSE-Chem, models receive the background and the literature that preceded a selected study and are tested on whether they propose a hypothesis matching its published finding 2. In ResearchBench, tasks derived from publications cover identifying the sources of ideas, formulating hypotheses, and ranking them 3. The published hypothesis, hidden from the model, serves as the reference finding. A low match rate does not show whether an alternative hypothesis would be confirmed if it were tested experimentally.

In BrainBench, a model chooses the original abstract from a pair in which the second abstract reports an altered finding, such as a reversed direction of effect, while the methods and background are unchanged. The model’s choice is the abstract with the lower perplexity. On 200 expert-written tasks, mean accuracy was 81.4% for 15 models and 63.4% for 171 neuroscientists (t(14) = 25.8; *p* < 0.001) 4. Our balanced set of confirmed and refuted statements (the 50/50 set) also pairs published findings with altered versions, but it uses a different presentation format and response procedure (Section 2.6), so accuracy on the two sets is not directly comparable.

In a study involving more than 100 natural language processing researchers, blinded reviewers judged ideas from a language-model agent more novel than those of human experts and slightly less feasible 5. In a follow-up study 6, 43 experts carried out projects based on such ideas, spending an average of 103 hours on each, and reviewers then evaluated the results. Co-Scientist adds laboratory testing of selected hypotheses to hypothesis generation, critique, and refinement 7. Automated research systems generate ideas, run computational experiments, and write up the results 8, or revise ideas iteratively to make them more novel relative to prior work 9. In forecasting future events, the aggregated forecast of several language models was statistically indistinguishable from that of a human crowd 10, and a language-model system approached the accuracy of the aggregated human crowd forecast 11. Agreement among multiple sampled answers to the same task helps identify the correct answer and improves reasoning accuracy 12, and large models can, for the most part, estimate the probability that their own answers are correct 13.

We evaluated generation by agreement with the hidden reference finding and evaluated forecasts for the generated hypotheses by the same criterion. Selection was considered effective if the match rate among selected hypotheses exceeded both the baseline match rate of the full set and the match rate of the random control. The set of confirmed and refuted statements also allowed “no” forecasts to be checked against known experimental outcomes.

### 1.3. Origin of the study

The Omega Point method was developed at Open Longevity to generate scientific hypotheses through a sequence of prompts with assigned creative roles. This study was conducted by the Eternal Search team, a separate Open Longevity project that was not involved in developing Omega Point. The work began by comparing an abridged Omega Point (without literature search) with single-request generation and a multistep pipeline; it was later extended to forecasting the experimental confirmation of existing hypotheses and to selecting hypotheses for priority testing.

### 1.4. Contributions of this work

We built hypothesis-generation tasks with a hidden experimental finding from 50 articles in ageing biology and checked their publication and preprint dates against the models’ stated training cutoffs. To evaluate negative forecasts, we compiled a set of confirmed and refuted statements, so that “no” answers could be checked against known experimental outcomes. The proposed selection rule keeps a hypothesis only if all four predictor models answer “yes”. The rule was chosen on the hypotheses of one generator and then applied to those of two others.

## 2. Materials and methods

This retrospective benchmark uses tasks derived from published studies (Figure 1). Generator models proposed hypotheses from the background and a question whose answer was hidden, judge models assessed whether each hypothesis matched the reference finding, and predictor models, which did not see the reference finding, forecast whether each hypothesis would be confirmed. The list selected from these forecasts was compared with the random control and with the hypotheses each generator listed first. Reporting was guided by the TRIPOD-LLM guideline for studies using large language models 14. The completed checklist maps each reporting item to a section of the manuscript.

**Figure 1.**
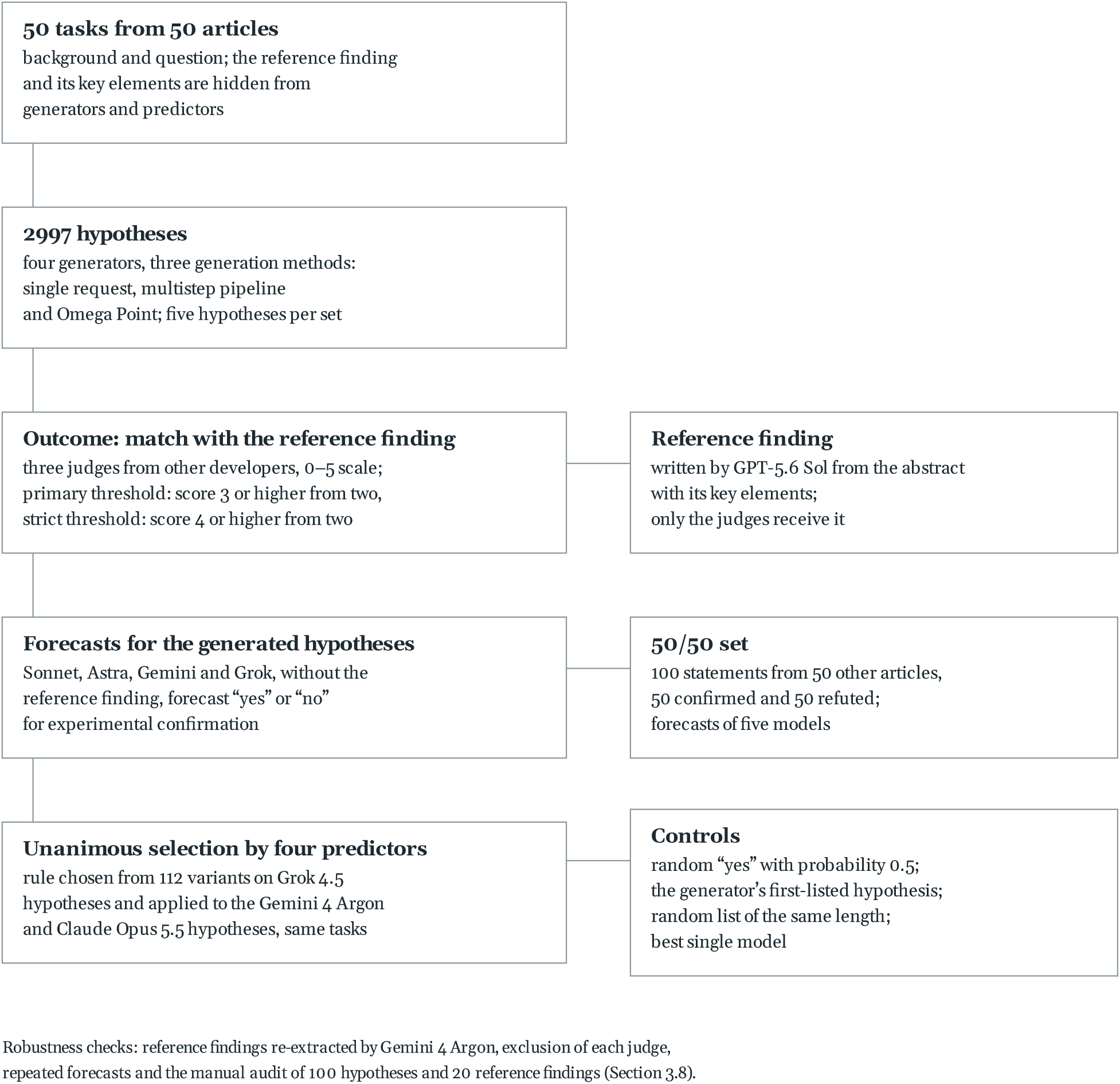
Study procedure. Only the judges received the reference finding. The selection rule was chosen on the Grok 4.5 hypotheses and applied to the Gemini 4 Argon and Claude Opus 5.5 hypotheses for the same tasks; the manual audit covers a sample of 100 hypotheses and 20 reference findings.

### 2.1. Article selection and task construction

We searched PubMed on 13 October 2026 (export timestamp 01:31:11 UTC). The ageing-biology query was restricted to 15 journals, the publication type Journal Article, and PubMed entry dates from 1 August to 12 October 2026; the full query is given in Appendix I. Of the 284 records, a preliminary filter excluded 42 because an earlier preprint was found and 30 because the abstract was shorter than 600 characters or the publication date fell before the start of the window, leaving 212 candidates. The order in which candidates were reviewed was fixed deterministically by a hash of the PubMed identifier (Appendix L.7).

To assemble the final set, we checked three conditions: suitability of the task, judged by GPT-5.6 Sol in the same request that constructed the task; concealment of the published finding in the background and the research question; and absence of an earlier preprint (Figure 2). The final 50 tasks are not a random sample of the 212 candidates. The search, exclusions, and candidate order are recorded in the selection log, and the completeness of the log is discussed in Appendix I.

**Figure 2.**
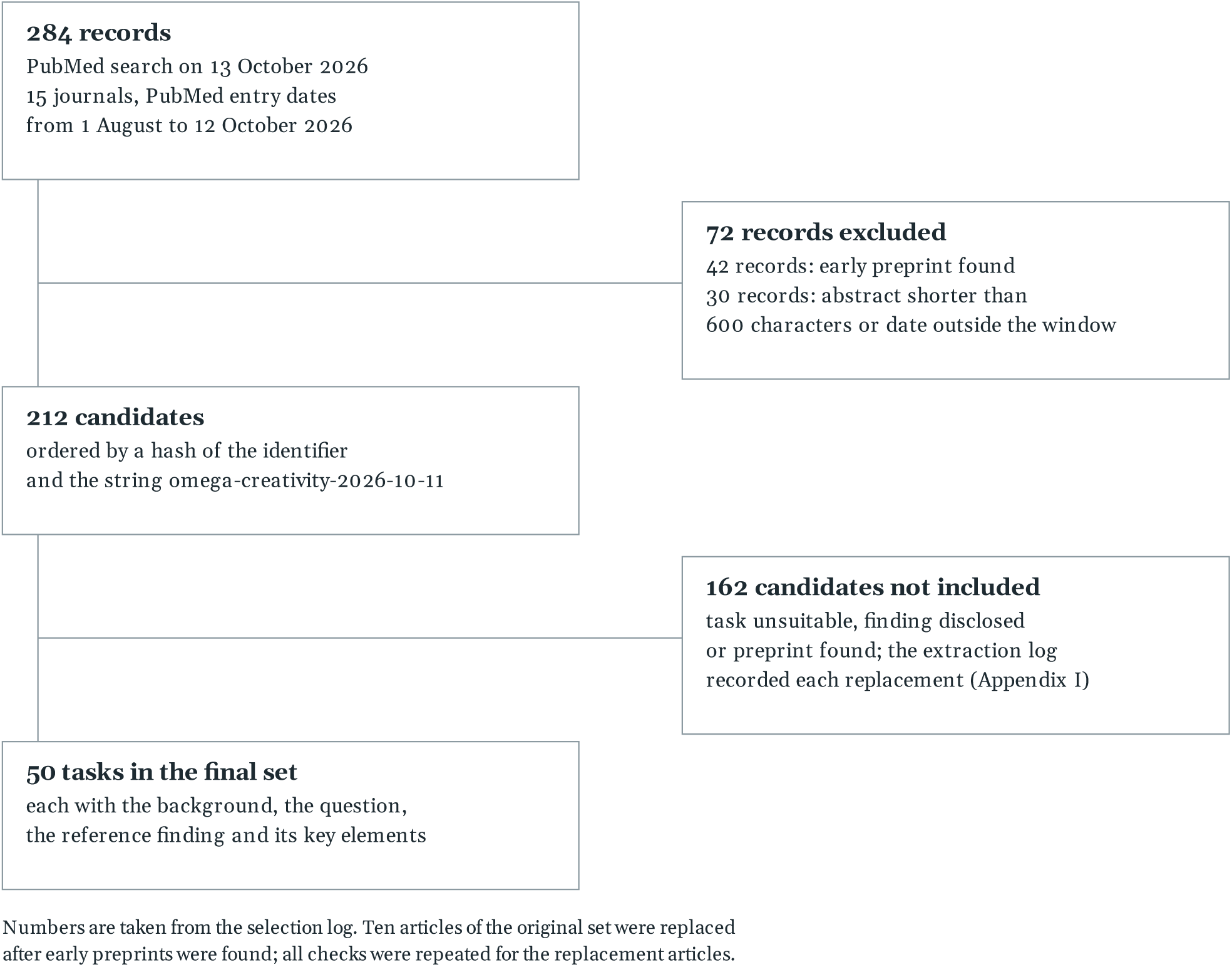
Article selection for the benchmark tasks.

The preliminary filter excluded 72 of the 284 records, and 50 of the 212 candidates entered the final set. Intermediate extraction logs were overwritten when articles were replaced, so the reasons for excluding the other 162 candidates cannot be reconstructed from those logs (Appendix I).

The final set comprised 50 peer-reviewed articles first published between 1 August and 9 October 2026 (20 in August, 20 in September, and 10 in October). They appeared in Aging Cell (22 articles), Science Advances (10), Nature Aging (8), Nature Communications (6), and PNAS, Science, Neuron, and Immunity (one each).

Eligibility under the date cutoff was based on the earliest public version of each study, including any preprint. Earlier versions were searched in six ways: preprint links in PubMed, records of preprint postings in OpenAlex, has-preprint relations in Crossref, author searches in Europe PMC, posted-content records in Crossref, and a separate preprint search in OpenAlex. Candidate versions were compared with the article by title and abstract, allowing for changes of title. Ten articles in the initial set had preprints posted before the common cutoff date; they were replaced, and date verification, hypothesis generation, and judging were repeated for the replacements. The final set was re-verified on 19 October 2026; the search for earlier versions covers the listed sources up to that date.

From the full PubMed abstract, GPT-5.6 Sol extracted the background, the research question, the reference statement, and a list of its key elements. The reference statement (hereafter the reference finding) is the experimental result reported in the peer-reviewed article. GPT-5.6 Terra checked whether the background or the research question disclosed the reference finding. Generators received only the background and the research question. To check whether the reference findings depended on the extracting model, Gemini 4 Argon at high reasoning effort extracted them again from the same abstracts with the same prompt, and Grok 4.7 compared the two versions of each finding (Section 3.8).

### 2.2. Models

Hypotheses were generated by GPT-6 Astra (hereafter Astra), Grok 4.5 (Grok), Claude Opus 5.5 (Opus), and Gemini 4 Argon (Gemini), all at high reasoning effort. Each model call was limited to 32,000 tokens, shared between reasoning and output, for all models and generation methods. The predictor models were Claude Sonnet 5.5 (Sonnet), Astra, Gemini, and Grok; Opus, Claude Haiku 5.5, and Claude Opus 4.8 served as predictors in separate runs. The settings of each model in each role are given in Appendix L.1.

The stated training cutoffs were January 2026 for Grok, March 2026 for Gemini, 30 April 2026 for Astra, and June 2026 for Opus and Sonnet. The developers call these dates knowledge cutoffs or pretraining data cutoffs; we used them as stated when selecting tasks. The date restriction did not apply to the judges, which received the reference finding explicitly.

All models ran with their default sampling settings (temperature and truncation parameters). No random seed was fixed; the saved model responses provide the basis for reproducing the analyses. A request was repeated if the response could not be parsed or the model refused to answer; network errors did not count as attempts. Model identifiers, settings, prompts, and responses are preserved in the logs.

### 2.3. Generation methods

#### Single-request generation (labelled N in the data)

The model received the background, the research question, and a prompt to propose five hypotheses in one response. The prompt asked for five hypotheses without ranking them; the position of a hypothesis in the response, from first to fifth, is the order in which it was generated. This method shows which hypotheses a model proposes directly from the task statement and is the point of comparison for the other two methods.

#### The multistep pipeline (A)

The pipeline is the control for the Omega Point method: it keeps the five stages of Omega Point and the transfer of results between them, and gives neutral stage instructions. Each stage was a separate request that contained the task and the results of all earlier stages; the model analysed the task, summarised established knowledge and existing approaches, formulated open questions, proposed hypotheses, and improved the three strongest of them. The pipeline omits the creative roles of Omega Point. Additional requests, each asking for new hypotheses different from those already proposed, brought its generated-token count close to that of Omega Point on the same task. In a final request, the model chose from all its candidates the five hypotheses it considered most likely to be confirmed by experiment and most valuable as answers to the research question, best first.

#### The Omega Point method (O)

Omega Point was designed to generate hypotheses with distinct mechanisms, including hypotheses that would force a revision of the accepted model of a field. We used an abridged five-stage version without literature search: analysing failures of the system described in the task; searching for contradictions and links to other fields; formulating questions about gaps in knowledge; generating hypotheses from the perspectives of assigned creative roles; and developing the chosen hypotheses, followed by final selection. As in the multistep pipeline, the stages were separate requests carrying the results of the earlier stages. At the third stage the request set one of five viewpoints on system failure, fixed for each task. At the fourth stage the system instruction required three creative roles: a hypothesis that would force a subfield to revise its core model; a transfer of a law or equation from physics, engineering, ecology, or materials science; and at least two hypotheses with mechanisms from fields distant from the task. In 10 of the 50 tasks, chosen by a hash of the task identifier, a fourth role was added: a hypothesis that the effect under study is a measurement artefact, an epiphenomenon, or several phenomena lumped together. At the fifth stage the request asked for one variant of each of the three strongest hypotheses, at least one of them obtained by inverting an assumption and at least one by transferring a mechanism from another field. The model chose five hypotheses with the same final request as in the multistep pipeline.

Across the four generators, the median number of candidates from which the model chose five hypotheses was 21.5 to 41 for the multistep pipeline and 7 to 11 for the Omega Point method (counts per set). The generation methods page describes the stages, roles, viewpoints of the third stage, the rule for additional requests, the number of requests per set, and how the stages of the abridged version correspond to the steps of the full Omega Point pipeline; the full prompts are on the prompts page.

Generated tokens are the recorded output and hidden reasoning tokens, excluding input prompt tokens; actual token usage is reported in Section 3.1.

During Opus generation, Claude’s safety classifier was triggered. A refusal counted as a non-match: a set that the model failed to generate in all attempts counted as five non-matching hypotheses, and hypotheses missing from an incomplete response were treated in the same way. The denominator was 250 hypotheses for each model–method combination; 2997 hypotheses were recorded in total.

Originality of research ideas was compared on a separate set of 100 scientific keywords in an adaptation of LiveIdeaBench. Each model proposed two ideas per keyword, and the judges rated originality (on a 1 to 10 scale), feasibility, clarity, and how distinct the two ideas were.

### 2.4. Judges and outcome definition

The judges were GPT-5.6 Sol, Grok 4.7, Claude Opus 5.5, and Gemini 4 Argon, run at medium reasoning effort (reasoning = medium). The outcome for each hypothesis was decided by the three judges from developers other than the generator’s, following the panel-of-diverse-models approach 15. This rule, adopted because judge models can favour their own outputs 16, was fixed on 13 October 2026, before the three-judge panel scored the saved hypotheses. In one request, each judge received the background, the research question, the reference finding with its key elements, and the full text of the five hypotheses of a set, shuffled into a pseudorandom order fixed for each task and generation method and labelled P1 to P5. The judge scored each hypothesis for agreement with the reference finding:

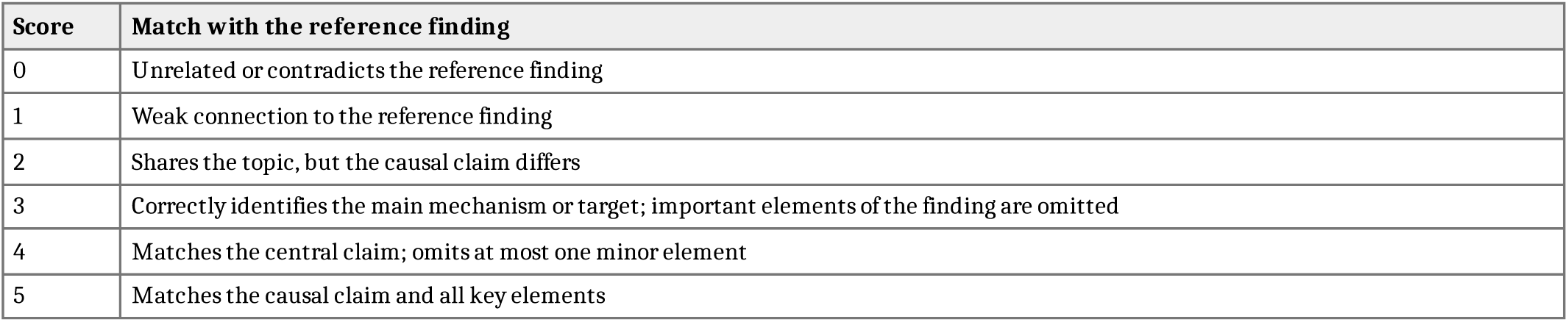

A hypothesis matched the reference finding if at least two of the three judges scored it at least 3 (the primary threshold). The primary threshold requires agreement on the mechanism or target and allows important elements of the finding to be missing. The levels of the scale measure how much of the finding a hypothesis reproduces, and they do not separate a key element that the hypothesis replaces from one it omits, so a hypothesis that names the mechanism of the article and substitutes one participant of that mechanism can reach 3. The strict threshold required a score of 4 or 5 from at least two judges. The majority decision was made separately for each threshold, and scores were not averaged. A score below the threshold indicates divergence from the reference finding; it does not mean that the hypothesis was refuted experimentally. The reference findings, the scale, and the scores of all four judges are published, so another judge can re-evaluate any hypothesis.

Inter-judge agreement was calculated for the binary vote “score of 3 or higher”, which decides the outcome at the primary threshold; all four judges scored each of the 2997 hypotheses. Cohen’s kappa across the six pairs of judges was 0.70 to 0.77, and Fleiss’ kappa for the three judges deciding the outcome was 0.72 (95% CI [0.67, 0.77]). For 283 hypotheses (9.4%), a single changed vote would reverse the majority decision. For the binary vote “score of 4 or 5”, pairwise Cohen’s kappa was 0.47 to 0.72, and Fleiss’ kappa was 0.57 (95% CI [0.41, 0.67]). Vote distributions and weighted kappa values are reported in Appendices J and L.2.

### 2.5. Forecasting with the detailed and short prompts

The forecasting benchmark can evaluate hypotheses from any source, independently of how they were generated. The two prompt variants were compared on the 250 hypotheses that Astra generated by a single request for the 50 tasks. In one request, the predictors Sonnet, Astra, Gemini, and Grok received the task background, the research question, and all five hypotheses in shuffled order with anonymised identifiers; the article title, the reference finding, the judge scores, and the answers of other models were withheld. In this comparison, Astra was both the generator and a predictor. Opus, Claude Haiku 5.5, and Claude Opus 4.8 also received the detailed prompt for the same 250 hypotheses.

The detailed prompt listed scientific criteria: the specificity of the causal claim, consistency with established biological data, testability, fit to the specified biological system, and the number of additional assumptions. For each hypothesis it required two separate answers: a subjective probability of experimental confirmation from 0 to 1 (or a statement that the data were insufficient) and a recommendation on whether to test the hypothesis experimentally (“yes” or “no”). The recommendation was taken directly from the response, not derived from the probability. The model also justified its decision, listed assumptions and contradictions, and proposed the intervention, the measured readout, and the control conditions of the experiment.

The short prompt was a single question (“For each hypothesis below, will it be confirmed experimentally?”) and required only a “yes” or “no” forecast. Responses to both prompt variants were returned in JSON format; the prompt texts are given in Appendix B. The detailed prompt thus asked for a recommendation to run an experiment, and the short prompt for a forecast of its outcome.

A response to a task was valid if it contained decisions on all five hypotheses in the required format. If either variant lacked a valid response for a task, all hypotheses of that task were excluded from both variants; technical failures were recorded separately from “no” answers. The paired comparison included 245 hypotheses from 49 tasks for Sonnet, 250 hypotheses from 50 tasks each for Astra and Gemini, and 225 hypotheses from 45 tasks for Grok.

### 2.6. Set of 50 confirmed and 50 refuted statements

A set with known positive and negative outcomes (hereafter the 50/50 set) was assembled from 50 other articles in ageing biology and related areas of experimental biology first published between 1 July and 5 October 2026. Articles already in the generation set were excluded by PubMed identifier. Earlier versions of each article were searched by the same six methods against a common cutoff date of 30 June 2026, and the search records were kept. Another 15 candidate articles were excluded before the statements were written because their preprint check could not be completed.

From the full text of the results sections and figure legends of each article, GPT-6.1 Sol at high reasoning effort wrote and verified two statements about the effect of an experimental intervention. The confirmed statement agreed with the measured outcome, and the refuted statement directly contradicted it. In 49 refuted statements, the predicted direction of change in the readout was opposite to the measured one. In the remaining statement, the intervention was predicted to preserve muscle stem cells after muscle injury, whereas the experiment showed that their pool was depleted. For each statement, we recorded the outcome used to assign the label, a verbatim quote from the article, and the figure or paragraph it came from.

Forecasts were obtained from Sonnet, Astra, Gemini, Grok, and Opus. Each request contained one statement and the question “Will this hypothesis be confirmed experimentally?”, with no request for an explanation or a numerical probability. The request omitted the answers to other statements, the article title and text, bibliographic identifiers, experimental outcomes, and labels, and the models were not told the ratio of the two groups. The two statements from one article could receive the same answer: unlike in BrainBench 4, the 50/50 ratio describes the composition of the set and imposes no forced choice within a pair. Each of the five models gave 100 valid forecasts.

### 2.7. Random control and consensus rules

For each hypothesis, a program drew a single “yes” or “no” answer, each with probability 0.5 (the Node.js functions crypto.randomInt and crypto.randomBytes; Appendix L.3). The cryptographic generator cannot be seeded, so the random answers were saved together with the hypothesis identifiers and generation timestamps, and the same sequence was used in all calculations. Whenever a model forecast was technically invalid, the corresponding random answer was excluded from the paired comparison. Each run was compared with its own random control. For the 250 Astra hypotheses, a second random sequence was drawn on 4 November 2026 from a seed fixed before the calculation (Appendix O.12).

The random control answers “yes” for about half of the hypotheses, whereas four-model unanimity keeps 19 to 22 of 240 to 250. We therefore also calculated the distribution of the number of matches in a random list of the same length as the predictor panel’s list, drawn either from the full set (hypergeometric distribution) or by random subsampling of each individual model’s positive forecasts (Appendices G and L.3).

The second control was the generator’s first-listed hypothesis in each task; it uses no forecasts and gives one hypothesis per task, 48 to 50 per set.

Joint selection was defined by the number of models answering “yes”: at least one, a majority, or all. The chosen rule determines list length, positive predictive value (PPV), and sensitivity jointly, so for unanimous selection we always report sensitivity alongside PPV.

### 2.8. Unanimous selection by four predictors

The selection rule was chosen on the saved forecasts for the hypotheses that Grok generated by a single request, the only set that, as of 23 October, had short-prompt forecasts from all five predictors. Each of four models (Sonnet, Astra, Gemini, and Grok) gave a forecast for every hypothesis (Appendix M.4), and the list included the hypotheses for which all four predictors answered “yes”. The 23 October protocol states the rule in two steps: a “yes” from Astra, then four-model unanimity. Because Astra is on the panel, both formulations give the same list. The panel holds one predictor per developer: Claude Sonnet 5.5 from Anthropic, GPT-6 Astra from OpenAI, Gemini 4 Argon from Google, and Grok 4.5 from xAI. No developer therefore casts two votes, and the same panel applies to a generator from any of the four developers; Claude Opus 5.5 stayed outside it. On the panel, Gemini ran at high reasoning effort, as in generation; as a judge, the same model ran at medium effort (Section 2.9).

On the Grok hypotheses, we evaluated 80 voting rules: every threshold for the number of “yes” answers, from one to unanimity, for each of the 31 non-empty combinations of the five models (Sonnet, Astra, Gemini, Grok, and Opus). We also examined 32 two-stage schemes that paired Astra or Sonnet at the first stage with each of 16 panel compositions at the second. The 112 procedural variants correspond to 90 distinct logical rules; the full enumeration is described in Appendix L.4, and the results are saved in the extended experiment report and the report on the 32 two-stage schemes. Each candidate list was re-evaluated at both matching thresholds and under both methods of parsing model responses. These evaluations are not independent, because the rules draw on the same forecasts.

For task R42725837, Grok’s forecasts were unavailable, and for task R42749700, Gemini’s response timed out. The ten hypotheses from these tasks were excluded from all comparisons because their forecasts were missing after technical failures, leaving 48 tasks and 240 hypotheses for the analysis of the Grok hypotheses. For Gemini and Opus, 50 tasks and 250 hypotheses each were analysed.

The rule chosen on the Grok hypotheses was then applied to the hypotheses generated by Gemini and Opus, with the same panel and the same unanimity requirement. New forecasts were obtained for these hypotheses; the tasks and judge scores stayed the same. The dataset composition and the selection rule were both locked on 23 October 2026, before the new forecasts were obtained (Appendix L.4).

### 2.9. Model isolation and role overlap

Reference findings were withheld from all requests to generators and predictors, and publication dates were checked to reduce the chance that the findings were in the training data. In all forecasting runs except the forecast on the full corpus of 2997 hypotheses (Appendix E), the saved responses, call logs, and request settings confirm that internet access and external tools were disabled.

Network access was detected in 51 Astra responses generated with the Omega Point method (17 hypothesis sets and 34 idea sets); these responses and the 424 judge verdicts on them were removed from the dataset and regenerated.

For the original generation and judging requests, the preserved logs do not confirm per request that tools and internet access were disabled (Section 5, item 6); a search of the 2916 saved raw responses of the four generators found no URLs, DOIs, PubMed identifiers, or mentions of the year 2026 (Appendix L.5). For the initial Opus generations (150 sets, 747 hypotheses, 788 requests, 13 to 18 October), the logs cannot confirm that tools were disabled: before a fix on 21 October, the tool-disabling parameter was dropped when requests were streamed, and the request logs for those dates were not kept. When the earlier configuration was reproduced, every attempt by the model to call a tool was rejected, and the initial generation responses contained no web links, DOIs, PubMed IDs, or mentions of the year 2026 (Appendix L.5). Opus generation was repeated on 27 and 28 October with the same tasks and prompts and with tools confirmed as disabled in every request, and the resulting hypotheses were scored by the same judges. This repeat generation serves as a sensitivity analysis; the initial set remains the primary dataset.

In the forecast on the full corpus of 2997 hypotheses, external predictors had not generated or judged the hypotheses for which they made forecasts, while each generator separately forecast its own hypotheses (Appendix E). In the other runs, Gemini at high reasoning effort forecast confirmation for hypotheses that Gemini at medium reasoning effort had previously scored as a judge, and Opus was both a judge and a predictor in the extended experiment on the Grok hypotheses. Judge scores and forecasts were collected in separate requests.

### 2.10. Statistical analysis

For each hypothesis, the binary outcome was compared with the binary forecast. The outcome was a match with the reference finding in the generation set and experimental confirmation in the 50/50 set. We calculated PPV, negative predictive value (NPV), sensitivity, specificity, balanced accuracy, and the area under the ROC curve (AUC); the definitions are given in Appendix L.8. In the generation set, a negative forecast for a hypothesis that matches the reference finding counted as a false negative. For each generation method, we calculated the overall match rate, the proportion of all 250 hypotheses that matched the reference finding. Among the hypotheses selected by positive forecasts, the match rate is the PPV.

Confidence intervals were calculated by cluster bootstrap over tasks (10,000 resamples; 2.5th and 97.5th percentiles). The resampling unit was the task with all its hypotheses, scores, and forecasts in the generation set, and the article with both its statements in the 50/50 set. In the analysis of unanimous selection, the full set, the final list, and the random control were recalculated together in each resample, with the same resamples used at both thresholds. When a denominator in a resample was zero, the value was treated as missing. Random number generator seeds are reported in Appendix L.6. The ranges reported for random lists of equal length describe random sampling of a list from a fixed dataset; they are not confidence intervals.

Generation methods were compared by permuting generation-method labels across whole response pairs within each task, with the difference in overall match rate as the test statistic. Exact enumeration was used when swapping labels changed the difference in at most 16 pairs; otherwise, 19,999 random permutations were used, with one added to both the numerator and the denominator 17. The procedure assumes that labels are exchangeable under the null hypothesis. Holm correction 18 was applied across the 12 within-model comparisons (three pairs of generation methods for each of the four models). The analysis was exploratory. Confidence intervals for individual differences were not adjusted, so statistical significance was judged from the adjusted *p*. With 19,999 permutations and adjustment for 12 comparisons, the smallest attainable value is 0.0006, so no threshold stricter than 0.001 is used. For the originality of research ideas, the Holm adjustment across the full family of 150 comparisons from our original analysis of the LiveIdeaBench adaptation was kept (Appendix L.6).

The intervals for the comparison of the two prompt variants and for the 50/50 set were calculated from 10,000 resamples of tasks or articles and are reported as descriptive statistics, without adjustment for the evaluation of all consensus rules.

The degree of cueing in each of the 50 research questions was rated on a 0 to 2 scale: 0 if only the phenomenon under study was named; 1 if a class of explanations, a source of perturbation, or a general process was indicated; and 2 if a substantial part of the reference finding was given, such as an intervention with its direction of effect or the key molecular participants.

GPT-6 Astra at high reasoning effort assigned the scores from the task texts and the reference findings after the initial hypothesis corpus had been collected; at that time, the model had access to the aggregate results and the examples involving AMP-activated protein kinase (AMPK) and motor neuron death. The scale, scores, and analysis plan were locked before they were linked to task-level generation outcomes. Later, Gemini 4 Argon and Grok 4.7 re-coded the 50 questions on the same scale from the task texts and reference findings, without access to any outcome.

The primary analysis was the Spearman correlation, over the 50 tasks and at both thresholds, between the cueing score and the overall match rate of single-request hypotheses averaged across the four generators. The *p* value was calculated from 99,999 permutations of scores across tasks, with Holm adjustment for the two tests. Further checks were exploratory (Appendix L.6).

Appendix L.6 describes the comparisons with the best single model at equal list length, ranking by the numerical probability of confirmation, selection without the predictor from the generator’s developer, within-pair decisions on the 50/50 set, lexical verification of key elements, the comparison of originality between selected and excluded hypotheses, and forecast reruns. New model responses were obtained on 30 October for the last two checks; all other checks used saved responses. Calculations were done in Python with NumPy 2.4.6.

We also performed the following analyses of the saved data (Appendix O):

1. the match rate by the position of a hypothesis in the generator’s response, with an exact test against random allocation of matches to positions within each generator and task;
2. grouping of hypotheses within a task by the cosine similarity of their statement embeddings;
3. the first-listed hypothesis as a control rule;
4. conditional logistic regression of a match on selection, stratified by generator and task, with first position, the share of named key elements, hypothesis length, and the numbers of named mechanisms and entities as covariates;
5. a leave-one-task-out estimate for the variant chosen among the 112;
6. random subsets of the best single model’s positive forecasts;
7. the generator ranking under each three-judge panel and under all four judges;
8. judge self-preference as a difference in differences of scores;
9. dependence between predictor errors compared with independence given the outcome;
10. the Brier score and reliability of the probability of confirmation;
11. results on the 50/50 set by publication date.

New model responses were obtained for four checks: the second extraction of the reference findings; the re-scoring of the selection sets against it by the same judges; forecasts by Gemini 3.8 Flash in place of Gemini 4 Argon under a protocol fixed before the first request; and the blind re-coding of question cueing.

One of us scored a frozen stratified sample of 100 of the 1000 single-request hypotheses on the judges’ 0 to 5 scale, with generator, judge scores, selection status, and stratum hidden. The sample comprised 30 selected hypotheses, 40 unselected hypotheses with split judge votes, and 30 with unanimous judge votes; each stratum’s sampling weight was its population size divided by its sample size. The same rater checked a separately frozen sample of 20 reference findings against 16 full texts and four abstracts. Binary agreement, Cohen’s kappa, and task-cluster bootstrap intervals were calculated at both thresholds (Appendix P).

### 2.11. Protocol deviations

The rule for handling Claude refusals was adopted on 15 October, while the initial generation set was being collected (13 to 18 October), and was then applied unchanged to all runs. In the initial generation set, two final Omega Point responses contained three and four hypotheses instead of five. In the first of them, a fourth-step response truncated by the safety classifier was saved together with the model’s retry, and the parsing script failed to extract the hypotheses from that step; the bug was fixed on 14 October. The missing hypotheses were neither filled in nor scored. The random control for the 250 Astra hypotheses was generated after the model results had been inspected, whereas for the 50/50 set the random sequence was locked before the forecasts were obtained; for the Astra hypotheses, a second sequence was drawn from a seed fixed before the calculation (Section 2.7). The protocol of the 50/50 experiment named balanced accuracy as the primary metric (Appendix H); the analysis of PPV and NPV, the within-pair decisions, and the comparison with BrainBench were added after the results were obtained. The cueing analysis was conducted after the initial hypothesis corpus had been collected. The analyses in Appendix O were added after the results were known. A fifth generator, Claude Fable 5.1, was part of the study plan. Its generation was stopped at 120 of 150 hypothesis sets because of its cost, so the analysis covers four generators; Fable is deferred to a later round. By then the judges had scored 50 Fable sets: 10.8% (27/250) of these hypotheses matched the reference finding, compared with 10.0% to 15.2% for the four generators on the same tasks and generation methods (judge scores). These scores were not extended to the other sets and are not part of the analysis.

### Box 1. Example of task R42744804

This task was built from the article by Huang et al. on motor neuron death in transgenic SOD1^G93A^ mice, a model of amyotrophic lateral sclerosis 19. The background described motor neuron loss, muscle denervation, and premature death of the animals, and mentioned that interactions between neurons and surrounding cells might be involved. The research question was: “What disease-associated mechanism drives motor neuron death and accelerates mortality in SOD1^G93A^ mice?”

According to the reference finding, microglia recognise phosphatidylserine exposed on still-viable neurons through the Axl and MerTK receptors and engulf them; genetic inactivation of these receptors preserved motor neurons and neuromuscular junctions and extended survival. Neither microglia, nor these receptors, nor the engulfment of viable cells was mentioned in the background or the research question.

With a single request, Gemini proposed five hypotheses. The second described engulfment of stressed but still-viable motor neurons by microglia and macrophages, involving phosphatidylserine, the MerTK receptor, and complement components, without naming Axl. GPT-5.6 Sol, Claude Opus 5.5, and Grok 4.7 scored it 3, 4, and 4, so the hypothesis matched the reference finding at both thresholds. Every judge scored each of the other four hypotheses below 3.

All four predictors forecast “no” for the matching hypothesis, so it was not selected. Astra forecast “yes” only for the hypothesis attributing neuron death to cytotoxic T lymphocytes and natural killer cells (judge scores 1, 1, and 2); that hypothesis did not receive a unanimous “yes”, so the final list for the task was empty. We chose this example because the answer requires a specific mechanism of neuronal death. Decisions on all five hypotheses are given in Appendix A.

### Box 2. Example of task R42806003

This task was built from the article by Hua et al. on placental senescence in advanced maternal age 20. According to the reference finding, macrophages express CD38 under the control of IRF5, deplete NAD^+^ in decidual stromal cells, and thereby promote placental senescence; CD38 inhibition or NAD^+^ precursors reduce senescence and rescue intrauterine growth restriction.

Grok’s first-listed hypothesis proposed paracrine signalling from senescent decidual stromal cells to trophoblasts (judge scores 2, 2, and 2), and all four predictors forecast “no” for it. The third hypothesis attributed placental senescence to NAD^+^ depletion by CD38 on decidual and circulating immune cells and predicted rescue by CD38 inhibition or NAD^+^ precursors; GPT-5.6 Sol, Claude Opus 5.5, and Gemini 4 Argon scored it 3, 4, and 4. All four predictors forecast “yes” for this hypothesis alone, and it formed the final list for the task.

## 3. Results

### 3.1. Matches between generated hypotheses and the reference finding

With a single request, 13.2% to 18.0% of the hypotheses from the four models matched the reference finding, compared with 11.2% to 18.8% with the multistep pipeline and 2.8% to 7.6% with the Omega Point method (Table 1).

**Table 1.** Match rates by generator and generation method.

| Generator | Single request, % (n/250) [95% CI] | Multistep pipeline, % (n/250) [95% CI] | Omega Point, % (n/250) [95% CI] |
|---|---|---|---|
| GPT-6 Astra | 14.4% (36/250) [9.6, 19.6] | 16.0% (40/250) [10.4, 22.0] | 3.2% (8/250) [0.8, 6.0] |
| Grok 4.5 | 16.8% (42/250) [11.6, 22.4] | 18.8% (47/250) [12.0, 26.4] | 2.8% (7/250) [0.8, 5.2] |
| Claude Opus 5.5 | 18.0% (45/250) [13.2, 23.2] | 11.2% (28/250) [5.2, 18.4] | 4.0% (10/250) [1.6, 6.8] |
| Gemini 4 Argon | 13.2% (33/250) [9.6, 17.2] | 17.2% (43/250) [12.4, 22.4] | 7.6% (19/250) [4.0, 11.6] |
| Claude Opus 5.5, repeat generation | 18.0% (45/250) [12.8, 23.6] | 14.4% (36/250) [9.2, 20.0] | 1.2% (3/250) [0.0, 2.8] |

The primary threshold requires a score of at least 3 from at least two of the three judges from other developers. Hypotheses missing because of Claude refusals count as non-matches. The Opus repeat-generation row is a sensitivity analysis (Section 2.9). Confidence intervals (CIs) are calculated from 10,000 resamples of the 50 tasks. The ranking of generators depends on the composition of the judge panel (Section 3.8).

After Holm correction, the multistep pipeline did not significantly outperform a single request for any model, and the estimated differences were positive for some models and negative for others (Table 2). The overall match rate with the Omega Point method was lower than with a single request for all four models (adjusted *p* < 0.05).

**Table 2.** Differences in overall match rate relative to single-request generation.

| **Model** | **Method** | **Difference, pp [95% CI]** | **Tasks with a higher match rate under the method** | **Tasks with a lower match rate under the method** | **Tasks with equal match rates** | **Holm-adjusted *p*** |
|---|---|---|---|---|---|---|
| GPT-6 Astra | Multistep pipeline | 1.6 [−3.2, 6.4] | 12 | 10 | 28 | 1.000 |
| GPT-6 Astra | Omega Point | −11.2 [−17.2, −6.0] | 3 | 20 | 27 | < 0.01 |
| Grok 4.5 | Multistep pipeline | 2.0 [−2.8, 7.2] | 14 | 12 | 24 | 1.000 |
| Grok 4.5 | Omega Point | −14.0 [−19.6, −8.8] | 3 | 26 | 21 | < 0.001 |
| Claude Opus 5.5 | Multistep pipeline | −6.8 [−12.0, −1.2] | 6 | 22 | 22 | 0.156 |
| Claude Opus 5.5 | Omega Point | −14.0 [−19.6, −8.8] | 2 | 25 | 23 | < 0.001 |
| Gemini 4 Argon | Multistep pipeline | 4.0 [0.4, 7.6] | 12 | 4 | 34 | 0.190 |
| Gemini 4 Argon | Omega Point | −5.6 [−9.2, −2.0] | 4 | 16 | 30 | < 0.05 |
| Claude Opus 5.5, repeat generation | Multistep pipeline | −3.6 [−8.4, 1.2] | 5 | 12 | 33 | 0.194 |
| Claude Opus 5.5, repeat generation | Omega Point | −16.8 [−22.0, −12.0] | 0 | 27 | 23 | < 0.001 |

The difference is in percentage points (pp); a positive difference means a higher overall match rate with the method. The three task-count columns in each row sum to 50. *P* values are Holm-adjusted for the 12 within-model comparisons (including the four comparisons of the Omega Point method with the multistep pipeline given below the table) and, for the Opus repeat generation, for its three within-model comparisons; CIs are unadjusted.

Compared with its control, the multistep pipeline, the Omega Point method gave a lower overall match rate for every generator: by 12.8 pp (95% CI [−18.8, −7.2]) for Astra, 16.0 pp (95% CI [−23.6, −9.2]) for Grok, 7.2 pp (95% CI [−14.0, −1.2]) for Opus, and 9.6 pp (95% CI [−14.4, −4.8]) for Gemini (Holm-adjusted *p* < 0.001, < 0.001, 0.190, and < 0.01).

At the strict threshold, 1.6% to 3.6% of single-request hypotheses matched the reference finding: the models named the mechanism studied in the article more often than they reproduced its central causal claim with its key elements (Appendix C).

Match rates varied with the position of a hypothesis in the response. Across the four generators, the first-listed hypothesis matched the reference finding in 37.5% of cases (75/200; 95% CI [26.5, 48.0]), and the hypotheses in positions 2 to 5 in 17.0%, 8.5%, 9.5%, and 5.5%. The difference between the first position and positions 2 to 5 was 27.4 pp ([17.6, 37.0]). In 75 of the 112 generator-task pairs with at least one match, the first-listed hypothesis matched, compared with 31.2 expected if matches were allocated to positions at random within each pair (*p* < 0.001). At the strict threshold, 20 of the 25 matches were first-listed. The judges saw the hypotheses in shuffled order, and the proportion of votes of 3 or higher was 13.7% to 18.5% across the five displayed positions (Appendix O.1).

Hypotheses from different generators often converged on the same claim. Within each task, hypotheses were grouped by the cosine similarity of their statement embeddings (connected components at a threshold of 0.75): 54.0% of the single-request hypotheses (540/1000) fell into groups containing hypotheses from all four generators, and 33.0% (330/1000) had no counterpart from another generator. The match rate was 22.2% (120/540; 95% CI [15.9, 29.0]) in the groups shared by all four generators and 7.6% (25/330; [4.8, 10.7]) among hypotheses without a counterpart. In 10 of the 50 tasks, no generator proposed a matching hypothesis with a single request, and in 18 tasks all four did (Appendix O.2).

The repeat Opus generation, with tools confirmed as disabled, gave the same overall match rate for a single request, 18.0% (45/250; paired difference 0.0 pp [−4.4, 4.4]). With the multistep pipeline, the overall match rate was 14.4%, compared with 11.2% in the initial generation (difference 3.2 pp [−2.4, 8.8]; *p* = 0.688 after Holm correction for three comparisons). With the Omega Point method, it was 1.2%, compared with 4.0% (−2.8 pp [−5.2, −0.4]; *p* = 0.188; with eight discordant tasks, the permutation test has low power). In the repeat generation, the overall match rate with the Omega Point method was also lower than with a single request (−16.8 pp [−22.0, −12.0]; *p* < 0.001). The difference persisted after excluding every task set in which at least one step contained a Claude refusal or a truncated response: −16.5 pp in the initial generation and −17.8 pp in the repeat generation (*p* < 0.001). Without the initial Opus generations, the overall match rate with the Omega Point method was lower than with a single request for each of the other three models (*p* < 0.05 after Holm correction for nine comparisons). Claude refusals and recalculations are reported in Appendix M.1.

Of the 50 research questions, 8 received a cueing score of 0, 30 a score of 1, and 12 a score of 2. With a single request, 25.8% of the hypotheses (62/240; 95% CI [14.7, 36.8]) matched the reference finding in the group scored 2, compared with 12.4% (94/760) in the other two groups; the difference was 13.5 pp (unadjusted 95% CI [2.1, 25.0]). Spearman’s correlation between the cueing score and the overall match rate was ρ = 0.17 at the primary threshold (95% CI [−0.15, 0.46]) and ρ = 0.17 at the strict threshold (95% CI [−0.16, 0.48]); *p* = 0.434 in both cases after Holm correction. On the 38 tasks scored 0 or 1, the overall match rate of single-request hypotheses was lower than on all 50 tasks for each of the four models (Appendix M.2). In the blind re-coding, Gemini 4 Argon and Grok 4.7 scored 5 and 6 questions as 2 (agreement with the original scores, Cohen’s kappa 0.59 and 0.47). With their scores, the match rate was 35.0% (35/100) and 37.5% (45/120) in the group scored 2, compared with 13.4% (121/900) and 12.6% (111/880) in the other groups (Appendix O.14). Both blind re-codings reproduce the direction of the association.

The ratio of generated tokens in the multistep pipeline to those in the Omega Point method was 0.978 for Astra, 1.080 for Grok, 1.059 for Opus, and 1.119 for Gemini. A single request used fewer tokens than either method for all four models (token data).

### 3.2. Originality of research ideas

As scored by the same judges from other developers, ideas from the Omega Point method received higher originality scores than ideas from a single request. After Holm correction for a family of 150 comparisons, the difference remained significant for Astra, Grok, and Gemini (*p* < 0.01); for Opus, the adjusted *p* was 0.205 (Table 3). Originality was evaluated on a separate set of keywords, so this result does not bear on matching the reference finding.

**Table 3.** Originality of ideas from the Omega Point method and from a single request.

| **Model** | **Keywords compared** | **Score difference, Omega Point minus single request [95% CI]** | **Keywords where Omega Point scored higher** | **Keywords where Omega Point scored lower** | **Tied keywords** | **Holm-adjusted *p*** |
|---|---|---|---|---|---|---|
| GPT-6 Astra | 89 | 0.39 [0.24, 0.55] | 53 | 19 | 17 | < 0.01 |
| Grok 4.5 | 100 | 0.69 [0.49, 0.89] | 69 | 21 | 10 | < 0.01 |
| Claude Opus 5.5 | 100 | 0.28 [0.07, 0.46] | 61 | 30 | 9 | 0.205 |
| Gemini 4 Argon | 98 | 0.37 [0.24, 0.49] | 59 | 19 | 20 | < 0.01 |

Scores are on a 1 to 10 scale. The comparison includes keywords with valid scores for both generation methods; the resampling unit is the keyword. Holm correction is applied across 150 comparisons (the composition of the family is given in Appendix L.6).

### 3.3. Comparison of the detailed and short prompts

Of the 250 Astra hypotheses, 36 matched the reference finding. With the detailed prompt, Astra recommended testing 200 hypotheses, of which 16.0% (32/200) matched the reference finding; the sensitivity of the recommendation was 88.9% (32/36). With the short prompt, Astra forecast “yes” for 75 hypotheses, of which 30.7% (23/75) matched, with a sensitivity of 63.9% (23/36). The difference in PPV was 14.7 pp (95% CI [7.8, 22.0]). For all four models, the point estimate of PPV was higher with the short prompt; for Astra and Gemini, the confidence interval for the difference lies above zero, and for Sonnet and Grok it includes zero (Table 4).

**Table 4.** PPV of the two prompt variants on the same hypotheses.

| **Predictor** | **Shared hypotheses** | **Detailed prompt, PPV, % (n/N) [95% CI]** | **Short prompt, PPV, % (n/N) [95% CI]** | **Difference in PPV, pp [95% CI]** |
|---|---|---|---|---|
| Claude Sonnet 5.5 | 245 | 20.6% (26/126) [13.2, 28.9] | 28.6% (14/49) [16.7, 40.9] | 7.9 [−0.1, 16.2] |
| GPT-6 Astra | 250 | 16.0% (32/200) [10.1, 22.6] | 30.7% (23/75) [20.0, 41.8] | 14.7 [7.8, 22.0] |
| Gemini 4 Argon | 250 | 21.1% (32/152) [13.4, 29.6] | 33.3% (21/63) [21.8, 45.6] | 12.3 [4.5, 20.8] |
| Grok 4.5 | 225 | 19.7% (27/137) [12.2, 27.9] | 23.5% (20/85) [14.1, 34.0] | 3.8 [−1.2, 9.2] |

For the detailed prompt, PPV refers to the recommendation to test a hypothesis; for the short prompt, to the forecast of experimental confirmation. The analysis includes hypotheses with a valid response under both variants; n is the number of matching hypotheses and N the number of positive decisions. The difference is calculated from paired resamples of tasks, and the intervals are descriptive. On the full set with the short prompt, Grok forecast “yes” for 94 hypotheses, 25 of which matched (26.6%).

Of the 50 hypotheses that Astra recommended not testing, 92.0% (46/50) did not match the reference finding; of the hypotheses with a “no” forecast under the short prompt, 92.6% (162/175) did not match. For the random control on the same 250 hypotheses, PPV was 14.5% (17/117) and NPV was 85.7% (114/133); for the second random sequence, PPV was 13.0% (17/131) and NPV 84.0% (100/119). The other metrics of the two prompt variants are reported in Appendix D, and those of the second random sequence in Appendix O.12.

The probability of confirmation elicited by the detailed prompt allows hypotheses to be ranked; its AUC on the same hypotheses ranged from 0.62 to 0.74 across the seven predictors. Mean predicted probabilities ranged from 0.24 to 0.51, compared with an observed match rate of 0.14. For Sonnet, the Brier score was close to that of a constant forecast equal to the match rate (skill score 0.01); for the other six models, it was higher (skill score −1.03 to −0.10), and the calibration slope was 0.77 to 1.46 (Appendix O.10). Unanimous selection by the four predictors, applied to their short-prompt answers for the 245 hypotheses answered by all four, kept 22 hypotheses, of which 50.0% matched the reference finding (11/22; 95% CI [30.0, 70.4]), compared with a baseline match rate of 14.7% (36/245). Among the 22 hypotheses with the highest probability of confirmation from each individual model, the expected match rate was 27.3% to 38.6% (for ties at the cut-off, the match rate within the tied group was used). The difference in favour of unanimous selection was 11.4 to 22.7 pp; *p* = 0.30 to 0.66 after Holm correction for seven models (Appendix K.1).

### 3.4. Forecasts for confirmed and refuted statements

On the 50/50 set, NPV (the proportion of refuted statements among those with a “no” forecast) was 73.6% to 79.5% for individual models and 52.1% (25/48) for the random control, and the PPV of the five models was 67.2% to 80.4% (Table 5). Balanced accuracy was 71.0% to 78.0% for the models and 52.0% for the random control (Appendix H). In this set, a negative outcome means direct experimental refutation, whereas in the generation set it means that a hypothesis does not match the reference finding.

**Table 5.** Forecasts of individual models on the 50/50 set.

| **Predictor** | **PPV, % (n/N) [95% CI]** | **NPV, % (n/N) [95% CI]** | **Sensitivity** | **Specificity** |
|---|---|---|---|---|
| Claude Sonnet 5.5 | 72.2% (39/54) [64.0, 81.5] | 76.1% (35/46) [66.7, 86.0] | 78.0% (39/50) | 70.0% (35/50) |
| GPT-6 Astra | 67.2% (41/61) [60.3, 75.5] | 76.9% (30/39) [66.7, 87.9] | 82.0% (41/50) | 60.0% (30/50) |
| Gemini 4 Argon | 80.4% (37/46) [70.6, 90.5] | 75.9% (41/54) [67.2, 85.4] | 74.0% (37/50) | 82.0% (41/50) |
| Grok 4.5 | 76.6% (36/47) [66.0, 87.2] | 73.6% (39/53) [64.0, 83.7] | 72.0% (36/50) | 78.0% (39/50) |
| Claude Opus 5.5 | 68.9% (42/61) [62.0, 76.9] | 79.5% (31/39) [70.0, 90.3] | 84.0% (42/50) | 62.0% (31/50) |
| Random control | 51.9% (27/52) [42.9, 61.1] | 52.1% (25/48) [42.2, 62.2] | 54.0% (27/50) | 50.0% (25/50) |

The set contains 100 statements from 50 articles, 50 of each kind (confirmed and refuted). CIs are calculated by resampling articles; the intervals for sensitivity and specificity are given in the structured report.

All five models forecast “yes” for 26 statements, 88.5% of which (23/26; 95% CI [76.7, 100.0]) were confirmed experimentally; for the random control, the proportion was 51.9% (27/52; 95% CI [42.9, 61.1]). The sensitivity of unanimity was 46.0% (23/50; 95% CI [32.0, 60.0]), so the other 27 confirmed statements were not selected. Results for other model combinations are given in the published report. On the 25 articles first published on or after 1 September 2026, balanced accuracy was lower than on the 25 earlier articles for four of the five models, including the three with training cutoffs from January to April. The mean change for Opus and Sonnet, whose cutoff is June 2026, differed from the mean change for the other three models by 6.3 pp (95% CI [−5.0, 18.0]); leakage of the earlier articles’ results into the training data would have made this difference negative (Appendix O.11).

Within pairs, the models accepted the confirmed statement and rejected the refuted one in 46.0% to 62.0% of the 50 pairs, and the random control did so in 24.0% (12/50); the models made the opposite decision in 1 to 5 pairs, and the random control made it in 10. The deterministic rule “effect in the favourable direction” labelled 61 of the 100 statements and agreed with the experimental outcome in 65.6% of them (40/61); on the same statements, the models answered correctly in 68.9% to 77.0% of cases. Of the 20 pairs in which the rule labelled both statements, the rule separated 5 correctly, the models separated 8 to 13, and the random control separated 5 (Appendices K.4 and M.8).

### 3.5. Unanimous selection across three generators on the same tasks

All four predictors forecast “yes” for 19 of the 240 Grok hypotheses, 14 of which matched. The PPV of the final list was 73.7% (14/19; 95% CI [52.6, 93.3]), compared with 17.9% (22/123) for the random control, and sensitivity was 35.0% (14/40; 95% CI [25.0, 46.2]; Table 6).

**Table 6.**
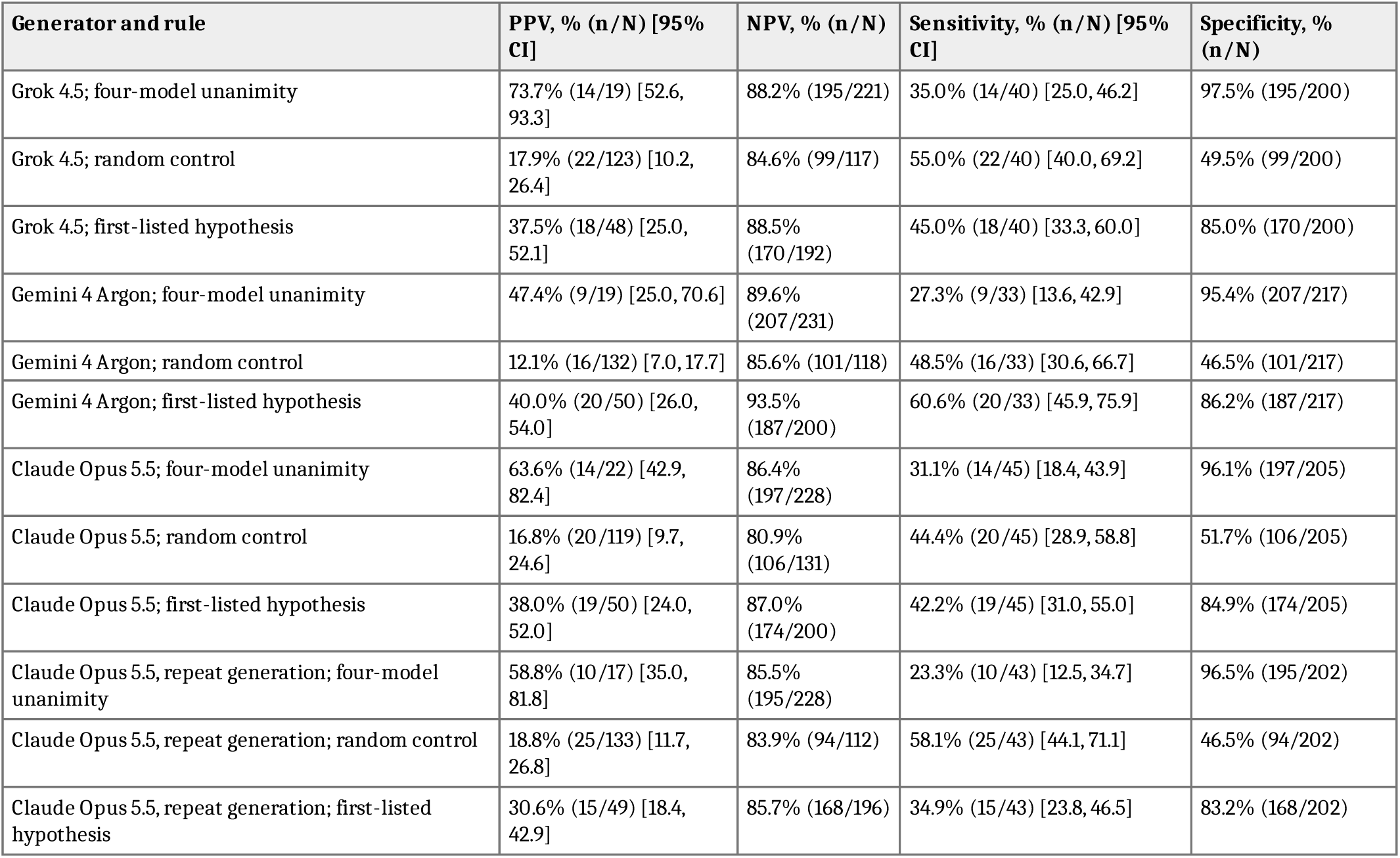
Unanimous selection, the random control, and selection of the first-listed hypothesis on the same hypothesis sets.

For Gemini, 19 hypotheses received a unanimous “yes”, 9 of which matched: PPV was 47.4% (95% CI [25.0, 70.6]) and sensitivity 27.3% (9/33; 95% CI [13.6, 42.9]). For Opus, 22 hypotheses received it, 14 of which matched: PPV was 63.6% (95% CI [42.9, 82.4]) and sensitivity 31.1% (14/45; 95% CI [18.4, 43.9]). Of the 245 hypotheses from the repeat Opus generation, 43 matched the reference finding; the final list contained 17 hypotheses, 10 of which matched. PPV was 58.8% (95% CI [35.0, 81.8]), compared with 18.8% for the random control (difference 40.0 pp [16.5, 62.6]), and sensitivity was 23.3% (95% CI [12.5, 34.7]).

Sets: Grok, 240 hypotheses, 40 of them matching; Gemini, 250 and 33; Opus, 250 and 45; the repeat Opus generation, 245 and 43 (the five hypotheses of one task were excluded because of a technical failure by Grok). CIs were calculated from paired resamples of tasks; full intervals are given in Appendix F.

In all sets, the match rate among selected hypotheses exceeded that of the random control. The intervals are wide, however, and the point estimates differ, so the 73.7% obtained on the Grok hypotheses cannot be generalised to other generators. On the 48 tasks shared by all three generators, the 95% CIs for all pairwise differences between generators included zero, both for the match rate among selected hypotheses and for sensitivity (Appendix M.3).

Unanimity selected mainly first-listed hypotheses: 50 of the 60 selected hypotheses in the three sets were listed first, and 6 of the 61 matching hypotheses in positions 2 to 5 were selected (Box 2). The match rate among unanimously selected hypotheses exceeded that among first-listed hypotheses by 36.2 pp (95% CI [19.2, 54.5]) for Grok, 25.6 pp ([9.7, 43.0]) for Opus, and 7.4 pp ([−12.7, 28.0]) for Gemini. Pooled across the three sets, the difference was 23.2 pp ([13.0, 32.6]), with sensitivity lower by 16.9 pp ([7.8, 27.5]). For the repeat Opus generation, the difference was 28.2 pp ([9.2, 48.6]). Compared directly on the tasks that two generators share, the gains differed by 28.4 pp (unadjusted 95% CI [1.4, 56.1]) between Grok and Gemini, by 18.3 pp ([−7.8, 44.6]) between Opus and Gemini, and by 8.7 pp ([−18.5, 34.6]) between Grok and Opus (*p* ≥ 0.16 after Holm correction across the three pairs; Appendix O.15). Within the tasks of each generator, the odds ratio of a match for selected hypotheses was 9.0 (95% CI [4.7, 21.2]); with first position as a second covariate, it was 3.3 ([1.5, 8.1]), and the odds ratio for first position was 4.1 ([2.1, 8.5]) (Appendix O.3).

Among the 112 variants evaluated on the Grok hypotheses, the chosen rule ranked second by PPV among those with lists of at least 10 hypotheses, after unanimity of all five models including Opus (75.0%, 12/16). Leave-one-task-out validation estimates the performance of the procedure that takes whichever variant a criterion ranks first: the highest-PPV variant among such lists was chosen on 47 tasks and applied to the held-out task, for each of the 48 tasks in turn, and 66.7% (12/18) of the hypotheses it selected matched the reference finding. The criterion picked unanimity of all five models in 46 of the 48 folds, the chosen rule in one, and unanimity of Sonnet, Astra, Grok, and Opus in the remaining one, so this value belongs to the procedure of choosing. The four-model rule was fixed on 23 October, and its own check is its application to the Gemini and Opus hypotheses, whose forecasts were obtained afterwards (Section 2.8, Appendix O.4).

### 3.6. Comparison with a random list and with the best single model

A random list of the same length drawn from the full set would contain on average 3.17 matching hypotheses out of 19 for Grok (central 95% range 0 to 6), 2.51 out of 19 for Gemini (0 to 6), and 3.96 out of 22 for Opus (1 to 8). The predictor panel selected 14, 9, and 14 matching hypotheses, above the upper end of the range in all three sets (Appendix M.6) and 4.4, 3.6, and 3.5 times the average for a random list of the same length.

The contribution of the panel relative to a single model was evaluated at equal list length. A random subset of the best single model’s positive forecasts, of the same length as the panel’s list, contained on average:

1. 9.0 matching hypotheses out of 19 for Grok (from Astra’s 53 positive forecasts; probability of at least 14 matches *p* = 0.004);
2. 9.4 out of 22 for Opus (Astra’s 49 positive forecasts; *p* = 0.009);
3. 5.9 out of 17 for the repeat Opus generation (Gemini’s 61 positive forecasts; *p* = 0.015);
4. 6.5 out of 19 for Gemini (Gemini’s 53 positive forecasts; *p* = 0.11).

On the Gemini hypotheses, the panel’s advantage did not reach statistical significance (Appendices G, K.2, and O.5).

Ranking all hypotheses of the three sets by the number of “yes” forecasts placed 37 matching hypotheses among the top 60, compared with 23.1 when the hypotheses were taken in the generators’ listing order and 9.6 for a random list. Among the top 148 hypotheses, the corresponding numbers were 59.6 for ranking by “yes” forecasts, 57.0 for the generators’ order, and 23.6 for a random list (Figure 3; Appendix O.5).

**Figure 3.**
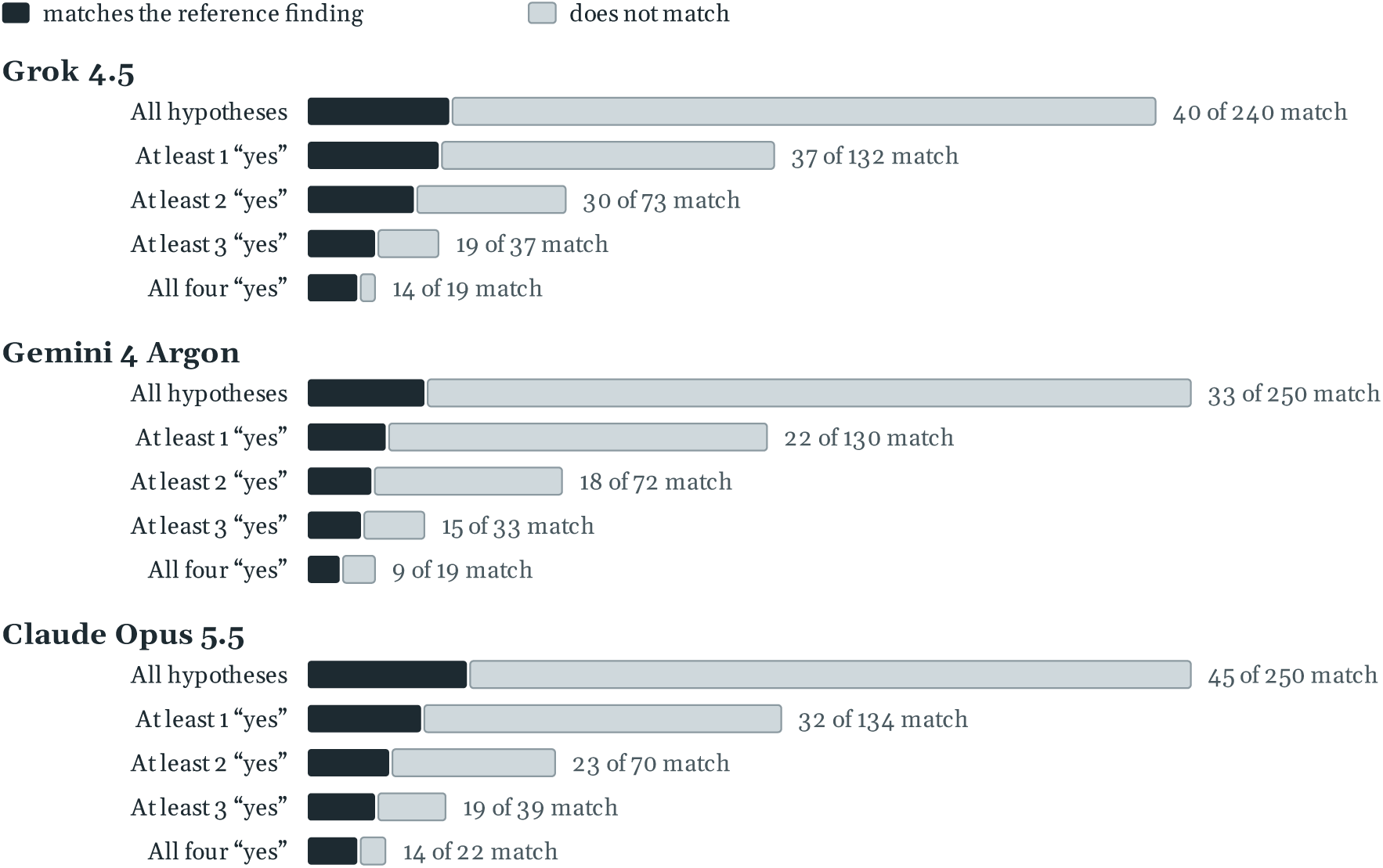
Hypotheses retained at each minimum number of “yes” forecasts.

The dark part of each bar shows hypotheses that match the reference finding. The last row of each set is the list selected by four-model unanimity.

### 3.7. The strict matching threshold

Under the strict threshold, the match rate among selected hypotheses was 26.3% (5/19) for Grok, 31.6% (6/19) for Gemini, and 9.1% (2/22) for Opus; its ratio to the baseline match rate at the same threshold is given in Table 7. Each selected list contains at most six matches under the strict threshold, the intervals are wide, and changing the threshold affects both the match rates and the ranking of generators by their point estimates (Appendices K.5 and M.5). Of the 37 matching hypotheses in the three selected lists, 13 also match at the strict threshold, and 16 received no score above 3 from any judge, agreeing with the mechanism or target of the reference finding while at least one of its key elements is missing or replaced (Appendix O.17).

**Table 7.** Ratio of the match rate among selected hypotheses to the baseline match rate under the strict threshold.

| **Generator** | **Ratio to the baseline rate** | **95% CI** | **Baseline match rate** | **Sensitivity** |
|---|---|---|---|---|
| Grok 4.5 | 9.02 | [4.21, 16.00] | 2.9% (7/240) | 71.4% (5/7) |
| Gemini 4 Argon | 8.77 | [5.18, 14.71] | 3.6% (9/250) | 66.7% (6/9) |
| Claude Opus 5.5 | 4.55 | [0.00, 11.03] | 2.0% (5/250) | 40.0% (2/5) |

The baseline match rate is the proportion of all hypotheses in the set that match under the strict threshold.

### 3.8. Robustness checks

On the 38 tasks with a cueing score of 0 or 1, the match rate among selected hypotheses was 63.6% (7/11) for Grok, 35.7% (5/14) for Gemini, 62.5% (10/16) for Opus, and 54.5% (6/11) for the repeat Opus generation. Its ratio to the baseline match rate on the same tasks was 5.0, 3.2, 4.2, and 4.2, with lower 95% CI bounds of 3.0, 1.1, 2.6, and 2.1.

When each judge was excluded in turn, the match rate among selected hypotheses was 68.4% to 73.7% for Grok, 47.4% for Gemini, 54.5% to 59.1% for Opus, and 52.9% to 58.8% for the repeat Opus generation; the lower 95% CI bounds of the ratio to the baseline rate were at least 2.0. Each of the four judges scored all 2997 hypotheses. Relative to the other judges, GPT-5.6 Sol scored Astra’s hypotheses 0.13 points higher (95% CI [0.07, 0.18]) and Gemini 4 Argon scored Gemini’s hypotheses 0.15 points higher ([0.10, 0.20]) on the 0 to 5 scale. For the vote of 3 or higher, a difference was found only for Grok 4.7 on Grok’s hypotheses (2.2 pp [0.6, 4.0]). Under each of the four fixed three-judge panels and under a majority of all four judges, Astra and Gemini ranked third and fourth by the number of single-request matches, whereas Opus and Grok swapped first and second place (Appendix O.6).

Without the predictor from the generator’s developer, unanimity of the remaining three models gave match rates of 68.2% (15/22) for Grok, 45.0% (9/20) for Gemini, 53.6% (15/28) for Opus, and 46.4% (13/28) for the repeat Opus generation. Without one of the other panel members, the rates were on average 64.0%, 47.0%, 59.9%, and 49.5%. The predictors showed no consistent preference for hypotheses from their own developer (Appendix K.3). Without Astra, unanimity of the other three models gave 60.9% (14/23), 43.5% (10/23), and 57.1% (16/28) for the Grok, Gemini, and Opus hypotheses; without Gemini, 69.6% (16/23), 45.0% (9/20), and 60.0% (15/25). Replacing Gemini 4 Argon with Gemini 3.8 Flash, which did not serve as a judge, gave 73.3% (11/15), 50.0% (8/16), and 57.9% (11/19) (Appendix O.7).

The errors of the four predictors were correlated: across the three sets, unanimous “yes” forecasts for non-matching hypotheses occurred 23 times, against 0.84 expected if the models erred independently given the outcome (ratio 27.5, 95% CI [18.3, 40.7]). The mean match rate among hypotheses selected unanimously by panels of one, two, three, or four models was 34.7%, 47.7%, 56.9%, and 61.7%, and the mean sensitivity was 53.4%, 40.7%, 34.7%, and 31.4% (Appendix O.8).

In the second extraction, Gemini 4 Argon accepted 47 of the 50 tasks; for all 47, Grok 4.7 judged the core finding, the mechanism or target, and the direction of the effect to be the same as in the version by GPT-5.6 Sol. The judges re-scored the single-request hypotheses of the three selection sets against the new reference findings: for the 695 hypotheses of these tasks, the outcome was the same under the new and original reference findings in 95.7% of cases (665/695). The match rate among unanimously selected hypotheses was 59.6% (34/57), compared with 61.4% (35/57) under the original reference findings. The advantage over first-listed hypotheses was 24.4 pp (95% CI [13.2, 36.0]), compared with 22.6 pp ([11.8, 33.6]) (Appendix O.9).

When the same requests were rerun, the decisions of individual models agreed with the initial ones in 88.3% to 96.2% of cases (Cohen’s kappa 0.71 to 0.89); of the 58 hypotheses selected in the initial run, 48 were selected again. The match rate among selected hypotheses in the rerun was 68.4% (13/19) for Grok, 50.0% (9/18) for Gemini, and 66.7% (14/21) for Opus, compared with 72.2% (13/18), 47.4% (9/19), and 61.9% (13/21) in the initial run on the same hypotheses (Appendix K.7).

For the 59 selected hypotheses compared, the mean originality score was lower than that of the excluded hypotheses from the same tasks (difference −0.21; 95% CI [−0.47, 0.04]; *p* = 0.118 after Holm correction across three measures). Feasibility of the selected hypotheses was higher by 0.45 points ([0.17, 0.73]; *p* < 0.01), and clarity was higher by 0.38 points ([0.24, 0.53]; *p* < 0.001). The odds of matching the reference finding were 4.72 times higher for selected than for excluded hypotheses (95% CI [2.36, 9.42]), and 3.85 times higher after adjustment for the mean clarity score ([1.81, 8.20]; *p* < 0.001) (Appendix K.6).

By a lexical criterion (at least two-thirds of an element’s content words present in the hypothesis text), at least one key element of the reference finding was named in 46.2% (72/156) of matching and 16.4% (138/844) of non-matching hypotheses among the 1000 obtained with a single request (odds ratio 4.39 [2.87, 6.81]). The AUC of the share of named elements for predicting the judges’ decision was 0.65 ([0.60, 0.70]) (Appendix K.8). Within the tasks of each generator, the odds ratio of a match for unanimously selected hypotheses was 9.0 (95% CI [4.7, 21.2]); it was 7.6 ([4.0, 17.9]) after adjustment for the share of named key elements, and 5.5 ([2.8, 13.1]) after further adjustment for hypothesis length and the numbers of named mechanisms and entities (Appendix O.13).

In the manual audit, with generator, judge scores, and selection outcome hidden, our classifications agreed with the judges’ at the primary threshold in 81.0% of cases (81/100; 95% CI [73.4, 88.2]; Cohen’s kappa 0.62 [0.46, 0.76]). With the sample strata weighted by their size in the corpus, agreement was 89.7% [77.1, 97.0] (kappa 0.64 [0.37, 0.87]). At the strict threshold, agreement was 92.0% (92/100; kappa 0.62). Of the 30 sampled selected hypotheses, we classified 63.3% as matches (19/30; 95% CI [41.7, 81.8]) and the judges classified 66.7% (20/30); the paired difference was −3.3 pp [−19.0, 10.7]. Of the 16 reference findings checked against the full articles, 4 received a score of 3, 11 a score of 2, and one a score of 1 on the accuracy scale described in Appendix P; the four reference findings checked against abstracts only all received a score of 3.

We checked four reference findings against abstracts only: nos. 1 (R42601407, Nat Aging), 4 (R42821663, Science), 11 (R42759510, Neuron), and 20 (R42822442, Immunity), whose full texts are not in PMC and are available from the publisher only by subscription. No discrepancies with the abstracts were found, so all four received a score of 3; this check, however, shows only that each reference finding does not distort the text from which it was compiled.

One reference finding overstated the causal support for the proposed mechanism. After excluding task R42844664, the pooled match rate among selected hypotheses was 62.1% (36/58), compared with 61.7% (37/60) in the full dataset (Appendix P).

Excluding all 12 tasks whose reference findings the audit scored below 3, the match rate among selected hypotheses was 63.0% (29/46) and the advantage over the first-listed hypotheses 24.7 pp (95% CI [13.0, 35.7]), compared with 61.7% (37/60) and 23.2 pp on all 50 tasks (Appendix O.16).

## 4. Discussion

### 4.1. Hypothesis generation and selection accuracy

Most single-request hypotheses diverged from the published finding of the selected article, and the outcome of testing them experimentally is unknown. The overall match rate therefore does not by itself show which hypotheses to test first.

Selection based on model forecasts produced short lists with a higher match rate among selected hypotheses, but these lists captured only 27.3% to 35.0% of all matching hypotheses. Per 100 single-request hypotheses, unanimity kept 7.6 to 8.8, of which 3.6 to 5.8 matched the reference finding; the 20 first-listed hypotheses per 100 included 7.5 to 8.0 matching ones. Under the strict threshold, the match rate among selected hypotheses was 9.1% to 31.6%.

The generators placed a matching hypothesis first more often than in later positions, and the first-listed hypotheses of different generators more often converged on the same claim (Section 3.1). Unanimity of the predictors selected mainly these hypotheses and raised the match rate among them by 7.4 to 36.2 pp, at a lower sensitivity (Section 3.5). The position of a hypothesis in the response may also explain the differences between generation methods in Table 2: with a single request, the generator probably lists first the hypothesis it considers most likely, whereas the multistep pipeline and the Omega Point method select five hypotheses from a larger pool of candidates.

### 4.2. Experimental confirmation and matching the reference finding

The predictors were asked whether a hypothesis would be confirmed experimentally, but their answers were evaluated by whether the hypothesis matched one published finding. An article may investigate one mechanism while a proposed hypothesis describes another that has not yet been tested. Divergence from the article therefore does not by itself refute a hypothesis.

In the 50/50 set, the outcomes of both groups are known, and five-model unanimity gave a PPV of 88.5% at a sensitivity of 46.0%. However, the confirmed and refuted statements were compiled from completed studies, and the ratio of the groups was set artificially; evaluating forecasts of future experiments requires prospective testing.

In BrainBench, removing context degraded performance, and its authors attributed successful choices to the integration of information on background and methods 4. In our set, a model received only the statement, without methods or background. The “effect in the favourable direction” rule leaves molecular readouts without an established sign unlabelled. Telling outcomes apart does not reduce to the sign of the expected effect: of the 20 pairs in which the rule labelled both statements, it separated 5 correctly, as did the random control, whereas the models separated 8 to 13.

### 4.3. Positive predictive value and outcome prevalence

PPV depends on the prevalence of positive outcomes in the sample:

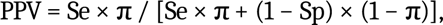

where Se is sensitivity, Sp specificity, and π the prevalence of positive outcomes. Values obtained on a balanced set therefore do not transfer directly to sets of generated hypotheses, in which matches with the reference finding are rare. With the sensitivity of 46.0% and specificity of 94.0% of five-model unanimity on the 50/50 set and π of 13.2% to 18.0%, the formula gives a PPV of 53.8% to 62.7%. The calculation shows only the direction of the adjustment: sensitivity and specificity were measured on statements with a reversed direction of effect, which are worded differently from generated hypotheses.

For the same reason, a high NPV is expected in the generation set. In the Astra set, 36 of 250 hypotheses matched the reference finding, so a model forecasting “no” for every hypothesis would be right in 85.6% of cases (214/250). An NPV of about 92% should therefore be read together with sensitivity and specificity. Selection with a high NPV can exclude suitable hypotheses: Astra forecast “no” for 15 of the 40 matching Grok hypotheses.

### 4.4. Detailed and short prompts

With the short prompt, PPV was higher for all four predictors, although for Sonnet and Grok the confidence interval for the difference included zero. The detailed prompt required estimating the probability of confirmation, justifying the decision, designing an experiment, and recommending whether to test, with the recommendation and the probability recorded as separate response fields. The short prompt required only a binary forecast. Because the content of the prompt and the nature of the required decision changed together, our data cannot separate the contribution of each.

### 4.5. Model agreement and independence of evidence

Unanimity shortened the list and increased the match rate among selected hypotheses relative to the full set. At equal list length, the predictor panel significantly outperformed the best single model for the Grok hypotheses and for both the initial and repeat Opus generations; the advantage was not statistically significant for the Gemini hypotheses; in a direct comparison of the three generators, the pairwise differences of the gain over the first-listed hypotheses were not statistically significant either (Appendix O.15). The models draw on shared sources of scientific literature, some come from the same developer, and in some runs one model served as both a judge and a predictor. Panickssery, Bowman, and Feng showed that judge models can favour their own outputs 16. The primary analysis excluded the judge from the generator’s developer, but other dependencies remain. Agreement among the models reflects correlated assessments: unanimous “yes” forecasts for non-matching hypotheses were 27.5 times more frequent (95% CI [18.3, 40.7]) than expected under independent errors (Section 3.8). Moreover, changing the generator changed the composition of the three judges, so differences between generators are not fully separated from differences between judge panels; the top two positions in the ranking of generators depended on the panel (Section 3.8). Judge models are sensitive to the order of candidate responses 21, 22; in our data, the hypotheses were shown in shuffled order, and the proportion of votes of 3 or higher was 13.7% to 18.5% across the displayed positions (Section 3.1).

The selected hypotheses received higher clarity and feasibility scores. If a clearly worded hypothesis raises both the judges’ score and the probability of a “yes” forecast, part of the enrichment is explained by clarity. After adjustment for the mean clarity score, the odds ratio of a match for selected hypotheses fell from 4.72 to 3.85 (95% CI [1.81, 8.20]).

### 4.6. Future work

We archived the 60 selected hypotheses from the three generators with their identifiers and the timestamps of the original forecasts and, as a baseline, every generator’s first-listed single-request hypothesis for the 50 tasks. The tracking and the test of the selection rule on new tasks are registered on OSF (osf.io/vj5xb, 5 November 2026) with the SHA-256 hashes of the registered files. We plan to compare the archived hypotheses with new publications and to test the selection rule on new research tasks. Such a test pre-specifies the background, the list of models, the selection rule, the criterion for the experimental outcome, and the observation window, as well as the requirements for matching the intervention, biological system, measured readout, and experimental conditions to the original hypothesis, and the procedure for independent evaluation. Each hypothesis will be assigned one of three outcomes: tested and supported by the data; tested and contradicted by the data; or outcome unknown, including when no publication testing it has appeared.

Unanimous selection by four models runs on the Check your hypotheses page of Eternal Search, where readers can apply it to their own hypotheses. Hypotheses from any source are accepted: each is rewritten in a single form in English without the author’s name, four models answer whether an experiment will confirm it, and the hypotheses that receive four “yes” answers are compared with the published literature. The forecasts on this page come from GPT-6 Astra, Claude Opus 5.5, Gemini 4 Argon, and Grok 4.7; this panel differs from the evaluated one in two members (Claude Opus 5.5 instead of Claude Sonnet 5.5 and Grok 4.7 instead of Grok 4.5), so the accuracy reported here refers to the evaluated panel.

## 5. Limitations

1. The reference findings were compiled by GPT-5.6 Sol from the abstracts of 50 peer-reviewed articles. In our audit of 16 reference findings against the full articles, the central finding was correct in 15 and all key elements were accurate in 4; this audit, the selection result recomputed without the tasks whose reference findings scored below 3, and the second extraction by Gemini 4 Argon are reported in Section 3.8.
2. The primary outcome was decided by the judge models; the manual audit covers a stratified sample of 100 single-request hypotheses (Appendix P). Judge agreement and the number of majority decisions that a single changed vote would reverse are reported in Section 2.4. Among the 60 hypotheses selected by the predictor panel across the three generators, there were 6 such decisions: 3 among the 37 matching and 3 among the 23 non-matching hypotheses.
3. The prompts and rules were developed on the same corpus (Appendix E and Section 3.6). The final rule was chosen after evaluating 80 voting rules and 32 two-stage schemes on the Grok hypotheses (90 distinct logical rules; Section 2.8). The confidence intervals do not account for this selection, so the performance of the chosen rule may be overestimated; Section 3.5 gives the leave-one-task-out estimate for the procedure that picks a variant by a criterion. The check of the chosen rule on the Gemini and Opus hypotheses used the same tasks and judge scores. The comparisons of generation methods were also exploratory.
4. In some runs, the same base model or models from the same developer served as both predictor and judge, which limits how far the panel can be treated as a set of independent sources (Section 3.8).
5. The temporal control relies on the models’ stated training cutoffs and on searches for earlier versions of the articles.
6. For the original generation and judging requests, including the initial Opus generations (747 hypotheses, 788 requests), and for the Opus and Sonnet responses in the forecast on the full corpus of 2997 hypotheses, the preserved logs cannot confirm that tools and internet access were disabled. A search of the 2916 saved raw responses of the four generators found no URLs, DOIs, PubMed identifiers, or mentions of the year 2026 (Appendix L.5). The reproduction of the original configuration and the repeat generation are described in Sections 2.9 and 3.1.
7. After unanimous selection by the four predictors, 19 to 22 hypotheses remain for each generator, and the PPV intervals are wide. Excluding tasks with technical failures limits the generalisability of the results to the full set of original tasks. On the Gemini hypotheses, the advantage of the panel over the best single model at equal list length and over the first-listed hypotheses did not reach statistical significance. Unanimous selection was compared with probability ranking on the Astra hypotheses, where Astra was both the generator and one of the four predictors (*p* = 0.30 to 0.66 after Holm correction; Section 3.3).
8. The random control for the 250 Astra hypotheses was added after the model results had been inspected; the second random sequence was drawn from a seed fixed before the calculation.
9. The 50/50 set is artificially balanced and was compiled by GPT-6.1 Sol from known findings. Of the 50 refuted statements, 49 reverse the measured direction of effect, so the set tests the models’ sensitivity to the direction of an effect. GPT-6.1 Sol and Astra were developed by OpenAI. The experimental format differs from that of BrainBench 4. These factors limit how far the findings generalise to laboratory practice.
10. The number of generated tokens was matched only approximately across generation methods: the ratio of token usage of the multistep pipeline to that of the Omega Point method was 0.978 to 1.119 (Section 3.1), and the multistep pipeline chose its five hypotheses from three to four times as many candidates (medians 21.5 to 41 against 7 to 11; Section 2.3).
11. The wording of the questions could cue the direction of the effect: the question for task R42810392 specifies lifespan extension upon direct activation of the target, although the target itself, AMPK, is hidden. The corpus included only articles with an experimentally supported finding, which may favour an affirmative restatement of the question. The analysis of cueing rests on annotations by AI models (the original annotator and two blind re-coders) and does not establish a causal effect (Sections 3.1 and 3.8).
12. The confidence intervals are calculated with the prompts, selection rule, model responses, and annotations held fixed and do not account for how these were chosen.
13. Of the 60 selected hypotheses, 50 were listed first by the generators (Sections 3.1 and 3.5), and the gain from selection is measured against the first-listed hypotheses.

## 6. Conclusion

1. Across the 50 tasks in ageing biology, 13.2% to 18.0% of single-request hypotheses matched the reference finding, and the multistep pipeline showed no statistically significant improvement over a single request for any model (Holm-adjusted *p* ≥ 0.156).
2. The abridged Omega Point method without literature search yielded fewer matches than a single request for all four models (*p* < 0.05 after Holm correction; for Opus, also after excluding task sets in which Claude refused to respond) and higher originality scores on a separate set of ideas, indicating that the two measures capture different properties of generation.
3. Unanimous selection by the four predictors raised the match rate among selected hypotheses to 47.4% to 73.7%, compared with 12.1% to 17.9% for the random control, and to 9.1% to 31.6% under the strict threshold, shortening the list for priority testing at a sensitivity of 27.3% to 35.0%.
4. The generators listed 50 of the 60 selected hypotheses first, and unanimity raised the match rate among selected hypotheses above that of the first-listed hypotheses by 23.2 pp across the three sets combined (95% CI [13.0, 32.6]); the first-listed hypothesis is therefore a necessary control when a selection rule is evaluated.
5. The advantage over the first-listed hypotheses and over the best single model at equal list length was statistically significant for the Grok and Opus hypotheses, while for the Gemini hypotheses the interval included zero, and the direct pairwise comparison of the gains gave *p* ≥ 0.16 after Holm correction, so these sets show an advantage of selection without showing how its size depends on the generator.
6. With reference findings re-extracted by a model from another developer, the advantage of selection over the first-listed hypotheses held (24.4 pp; 95% CI [13.2, 36.0]), whereas the top two positions in the ranking of generators changed with the judge panel, so the conclusion about selection is more robust than the ranking of generators.
7. The outcome of experimental testing for hypotheses that diverge from the reference finding remains unknown; we plan to track these hypotheses in future publications and to test the selection rule on new tasks.

## Data and code availability

The results are published at eternalsearch.net/omega/benchmark. The tasks, reference findings, and examples of judge evaluations are available on the tasks page. All selected hypotheses with their full text, reference findings, and the scores of the three judges are given in Table S. The manual-audit results and item-level scores, protocols, saved prompts, model responses, and final selection lists are published on the website; Appendix N lists the directories of the forecasting runs and of the final selection lists. The text of this preprint, its supplementary materials, the data files and the code archives are released under the Creative Commons Attribution-NonCommercial 4.0 International licence (CC BY-NC 4.0); material from cited publications remains under its original terms.

Attached to the manuscript are the hypothesis outcomes, the scores of the four judges, the calculation results, the supplementary calculation report, and the data description. The tables in the manuscript were generated by code from these files, without calling the models, and Table 7 repeats the numbers of Appendix K.5; checksums of the source files are published with the results. The analysis scripts and their input data are packaged in an archive. The analyses in Appendix O were calculated from the published files and the saved model responses. Their scripts, the input files they read, and the result files they write are packaged in a second archive, and each script is also published as a separate text document.

## Declarations

### Author contributions

Yuriy Poteshkin: conceptualization, methodology, software, formal analysis, investigation, data curation, visualization, writing – original draft, writing – review and editing, project administration, and supervision (all lead). Evgenii Udodov: resources (lead) and supervision (supporting). Mikhail Batin: conceptualization, supervision, and writing – review and editing (all supporting).

## Use of language models

Language models were used to prepare the analysis scripts and to edit the text; the calculations were performed on the recorded data, and the authors verified the text and conclusions.

## Acknowledgements

We thank Daniil Pimanov for an independent reanalysis of the published data.

## Human and animal research

This work analyses published studies and responses from language models. No new laboratory experiments were conducted for this benchmark.

## Appendix A. Five hypotheses on motor neuron death

This example shows how a matching hypothesis was lost during selection (Box 1). The Gemini hypotheses for task R42744804 are listed in their original order of generation; before being shown to the predictors, they were shuffled and labelled H1 to H5. Judge scores are given in the order GPT-5.6 Sol, Claude Opus 5.5, and Grok 4.7. The table summarises the main claims; the models received the full text, preserved in the original report.

| Hypothesis in the original Gemini response | Main claim | Judge scores | Astra forecast | Four-model unanimity | Random control |
|---|---|---|---|---|---|
| 1, identifier H1 | Microglia shift astrocytes into a state in which they release toxic substances and cause motor neuron death | 2, 1, 2 | No | No | Yes |
| 2, H2 | Microglia and macrophages engulf neurons that are still alive after recognising phosphatidylserine and complement components; MerTK is named among the receptors | 3, 4, 4 | No | No | Yes |
| 3, H4 | Senescent glial cells release inflammatory substances that cause motor neuron death | 1, 1, 2 | No | No | Yes |
| 4, H3 | Cytotoxic T lymphocytes and natural killer cells recognise and kill motor neurons | 1, 1, 2 | Yes | No | Yes |
| 5, H5 | Release of mitochondrial DNA activates the innate immune response and causes cell death through membrane pore formation and lipid peroxidation | 1, 0, 2 | No | No | Yes |

The second hypothesis matches the reference finding at both thresholds, but all four predictors forecast “no” for it. The fourth hypothesis received a “yes” forecast from Astra and Grok and a “no” from Sonnet and Gemini. As a result, none of the five hypotheses entered the final list. The random control gave five “yes” answers in this example; it plays no part in the panel’s decision.

## Supplementary materials

Appendices B to P are given in the supplementary materials: prompts for the predictors (B), the strict matching threshold (C), full metrics comparing the prompt variants (D), the forecast on the full corpus of 2997 hypotheses (E), full intervals for unanimous selection (F), lists of equal length from individual model forecasts (G), forecast accuracy on the 50/50 set (H), the search query and article selection log (I), judge agreement (J), additional selection and forecast calculations (K), further details of methods and definitions of metrics (L), additional results (M), contents of the published data (N), control rules and robustness checks (O), and the manual audit of hypotheses and reference findings (P).

## References

1. Ruan K, Wang X, Hong J, Wang P, Liu Y, Sun H. Evaluating LLMs’ divergent thinking capabilities for scientific idea generation with minimal context. Nature Communications. 2026;17:3625. doi:10.1038/s41467-026-70245-1.

2. Yang Z, Liu W, Gao B, Xie T, Li Y, Ouyang W, Poria S, Cambria E, Zhou D. MOOSE-Chem: Large Language Models for Rediscovering Unseen Chemistry Scientific Hypotheses. ICLR 2025. arXiv:2410.07076.

3. Liu Y, Yang Z, Xie T, Ni J, Gao B, Li Y, Tang S, Ouyang W, Cambria E, Zhou D. ResearchBench: Benchmarking LLMs in Scientific Discovery via Inspiration-Based Task Decomposition. Findings of the Association for Computational Linguistics: ACL 2026. 2026:13187–13207. doi:10.18653/v1/2026.findings-acl.644.

4. Luo X, Rechardt A, Sun G, Nejad KK, Yáñez F, Yilmaz B, et al. Large language models surpass human experts in predicting neuroscience results. Nature Human Behaviour. 2025;9(2):305–315. Published online 27 November 2024. doi:10.1038/s41562-024-02046-9.

5. Si C, Yang D, Hashimoto T. Can LLMs Generate Novel Research Ideas? A Large-Scale Human Study with 100+ NLP Researchers. ICLR 2025. arXiv:2409.04109.

6. Si C, Hashimoto T, Yang D. The Ideation-Execution Gap: Execution Outcomes of LLM-Generated versus Human Research Ideas. ICLR 2026. arXiv:2506.20803.

7. Gottweis J, Weng WH, Daryin A, Tu T, Sirkovic P, Myaskovsky A, et al. Accelerating scientific discovery with Co-Scientist. Nature. 2026;655(8122):487–496. doi:10.1038/s41586-026-10644-y.

8. Lu C, Lu C, Lange RT, Foerster J, Clune J, Ha D. The AI Scientist: Towards Fully Automated Open-Ended Scientific Discovery. 2024. Preprint. arXiv:2408.06292.

9. Wang Q, Downey D, Ji H, Hope T. SciMON: Scientific Inspiration Machines Optimized for Novelty. Proceedings of the 62nd Annual Meeting of the Association for Computational Linguistics (Volume 1: Long Papers). 2024:279–299. doi:10.18653/v1/2024.acl-long.18.

10. Schoenegger P, Tuminauskaite I, Park PS, Bastos RVS, Tetlock PE. Wisdom of the silicon crowd: LLM ensemble prediction capabilities rival human crowd accuracy. Science Advances. 2024;10(45):eadp1528. doi:10.1126/sciadv.adp1528.

11. Halawi D, Zhang F, Yueh-Han C, Steinhardt J. Approaching Human-Level Forecasting with Language Models. Advances in Neural Information Processing Systems. 2024;37:50426–50468. doi:10.52202/079017-1598.

12. Wang X, Wei J, Schuurmans D, Le QV, Chi EH, Narang S, et al. Self-Consistency Improves Chain of Thought Reasoning in Language Models. ICLR 2023. arXiv:2203.11171.

13. Kadavath S, Conerly T, Askell A, Henighan T, Drain D, Perez E, et al. Language Models (Mostly) Know What They Know. 2022. Preprint. arXiv:2207.05221.

14. Gallifant J, Afshar M, Ameen S, Aphinyanaphongs Y, Chen S, Cacciamani G, et al. The TRIPOD-LLM reporting guideline for studies using large language models. Nature Medicine. 2025;31(1):60–69. doi:10.1038/s41591-024-03425-5.

15. Verga P, Hofstätter S, Althammer S, Su Y, Piktus A, Arkhangorodsky A, et al. Replacing Judges with Juries: Evaluating LLM Generations with a Panel of Diverse Models. 2024. Preprint. arXiv:2404.18796.

16. Panickssery A, Bowman SR, Feng S. LLM Evaluators Recognize and Favor Their Own Generations. Advances in Neural Information Processing Systems. 2024;37:68772–68802. doi:10.52202/079017-2197.

17. Phipson B, Smyth GK. Permutation P-values Should Never Be Zero: Calculating Exact P-values When Permutations Are Randomly Drawn. Statistical Applications in Genetics and Molecular Biology. 2010;9(1):Article 39. doi:10.2202/1544-6115.1585.

18. Holm S. A Simple Sequentially Rejective Multiple Test Procedure. Scandinavian Journal of Statistics. 1979;6(2):65–70. JSTOR 4615733.

19. Huang Y, Mavinkurve A, Sabikunnahar B, Stevens B, Lemke G. Microglia deploy TAM receptors to kill motor neurons in a mouse model of amyotrophic lateral sclerosis. Nature Communications. 2026;17:9837. doi:10.1038/s41467-026-76728-5.

20. Hua L, Yang X, Gao D, Yang X, Jin X, Ma L, et al. CD38-activated macrophages drive age-related placental senescence by depleting NAD+ in decidual stromal cells. Nature Communications. 2026;17:10286. doi:10.1038/s41467-026-77217-5.

21. Zheng L, Chiang WL, Sheng Y, Zhuang S, Wu Z, Zhuang Y, et al. Judging LLM-as-a-Judge with MT-Bench and Chatbot Arena. Advances in Neural Information Processing Systems. 2023;36:46595–46623. doi:10.52202/075280-2020.

22. Wang P, Li L, Chen L, Cai Z, Zhu D, Lin B, et al. Large Language Models are not Fair Evaluators. Proceedings of the 62nd Annual Meeting of the Association for Computational Linguistics (Volume 1: Long Papers). 2024:9440–9450. doi:10.18653/v1/2024.acl-long.511.

