## Supplementary Materials for "Benchmarking four large language models on generating and selecting hypotheses that match published findings in 50 studies of ageing biology"

### Supplementary materials to the preprint on the hypothesis benchmark

The main text is the preprint. Appendix lettering continues from the main text, where Appendix A gives an example of five hypotheses. Table S lists all selected hypotheses with their reference findings and the judges' scores.

#### Appendix B. Prompts given to the predictors

The substantive prompts used in the experiments are reproduced below in their original English; technical instructions on the answer format and the absence of tools, appended to some requests, are preserved in the published request records. Bracketed placeholders were replaced with the full text of the task and the hypotheses. The full request records for each model are in the published materials listed under "Data and code availability".

##### B.1. The detailed prompt

System prompt:

```
You advise researchers which causal hypotheses deserve experimental testing in the biology of ageing.
Use only the supplied background, research question, hypothesis text and your prior scientific knowledge. You have no tools or internet access. Experimental outcomes are unknown to you.
Forecast experimental support for each hypothesis's own causal claim in its stated biological system. Return a complete JSON object in this reply.
```

User message:

```
BACKGROUND
[BACKGROUND]

RESEARCH QUESTION
[RESEARCH QUESTION]

CANDIDATE HYPOTHESES
[H1] [FULL HYPOTHESIS 1]

[H2] [FULL HYPOTHESIS 2]

[H3] [FULL HYPOTHESIS 3]

[H4] [FULL HYPOTHESIS 4]

[H5] [FULL HYPOTHESIS 5]

Assess EVERY hypothesis independently. Several different mechanisms may be supported, and none may be
supported. There is no required number of recommendations and no requirement to recommend one hypothesis per
question.

For each hypothesis estimate p_experimental_support between 0 and 1: your probability that a well-controlled
discriminating experiment, competently performed in the species, tissue and scope stated in the hypothesis,
would support its central causal claim. Evaluate that claim itself, including a plausible alternative
mechanism. This is a subjective scientific forecast, not an observed frequency. If the supplied information
cannot justify any numerical estimate, return null and explain the missing information.
```

```
Give recommend_for_testing as a separate boolean: would you advise a researcher to spend experimental
resources testing this hypothesis now, given its scientific support, testability, assumptions and
contradictions? You may recommend multiple hypotheses or none. A recommendation is not a guarantee of success.
A numerical estimate is required for a positive recommendation. The comparison will retain your recommendation
exactly as issued.

Use the existing scientific considerations: a specific causal chain; consistency with established biology; a
feasible discriminating test; fit to the stated question and system; few unsupported assumptions. Novelty is
descriptive and is not evidence for or against a claim. Ignore prose style, length, tone, gene-name density
and the anonymous candidate order. Do not guess the contents of an undisclosed article or predict another
evaluator's score.

Provide a short scientific justification. Quote only literal fragments of the supplied BACKGROUND in
support_from_background, or use an empty array. Keep additional assumptions and contradictions separate. Name
a concrete intervention, measured readout and necessary controls, and specify outcomes supporting and refuting
the claim. These are proposed observations, not results you have seen. Do not announce research, search for
information or describe work you will do later; give the complete forecasts now.

Return exactly this structure, with every supplied candidate once:
{"protocol":"experimental-support-v1","forecasts":
[{"id":"H1","p_experimental_support":0.5,"recommend_for_testing":false,"support_from_background":
[],"assumptions":[],"contradictions":[],"reason":"Short justification","test":
{"intervention":"...","measurement":"...","controls":"...","supports_if":"...","refutes_if":"..."}}]}
```

##### B.2. The short prompt for five hypotheses

System prompt:

```
Return a JSON object.
```

User message:

```
For each hypothesis below, will it be confirmed experimentally?

BACKGROUND
[BACKGROUND]

RESEARCH QUESTION
[RESEARCH QUESTION]

HYPOTHESES
[H1] [FULL HYPOTHESIS 1]

[H2] [FULL HYPOTHESIS 2]

[H3] [FULL HYPOTHESIS 3]

[H4] [FULL HYPOTHESIS 4]

[H5] [FULL HYPOTHESIS 5]

Return a JSON object with a "forecasts" array. For each hypothesis include its "id" and a boolean "confirmed".
```

##### B.3. The short prompt for one statement

System prompt:

```
Return a JSON object.
```

User message:

```
Will this hypothesis be confirmed experimentally?

HYPOTHESIS
[H000000000001] [FULL HYPOTHESIS]

Return a JSON object with a "forecasts" array containing this hypothesis's "id" and a boolean "confirmed".
```

#### Appendix C. Strict threshold for matching the reference finding

The outcome was positive if at least two of the three judges from other developers scored the hypothesis 4 or 5. Each cell gives the percentage, the numerator/denominator, and the 95% confidence interval (CI) from 10,000 resamples of tasks. Match rates under the strict threshold were calculated separately from those under the primary threshold (scores of 3 to 5).

When no hypothesis in the observed set matches the reference finding, the resampling percentile method gives a degenerate interval of 0.0 to 0.0. Such an interval reflects the composition of the observed set and gives no upper bound for the match rate in new tasks.

| Model | Generation method | Score of 4 or 5 from a majority of judges (95% CI) |
|---|---|---|
| GPT-6 Astra | Single request | 1.6% (4/250; 95% CI [0.4, 3.2]) |
| GPT-6 Astra | Multistep pipeline | 1.2% (3/250; 95% CI [0.0, 2.8]) |
| GPT-6 Astra | Omega Point | 1.2% (3/250; 95% CI [0.0, 3.2]) |
| Grok 4.5 | Single request | 2.8% (7/250; 95% CI [1.2, 4.8]) |
| Grok 4.5 | Multistep pipeline | 4.0% (10/250; 95% CI [1.2, 7.2]) |
| Grok 4.5 | Omega Point | 0.0% (0/250; 95% CI [0.0, 0.0]) |
| Claude Opus 5.5 | Single request | 2.0% (5/250; 95% CI [0.4, 4.0]) |
| Claude Opus 5.5 | Multistep pipeline | 1.2% (3/250; 95% CI [0.0, 3.2]) |
| Claude Opus 5.5 | Omega Point | 0.0% (0/250; 95% CI [0.0, 0.0]) |
| Gemini 4 Argon | Single request | 3.6% (9/250; 95% CI [1.2, 6.4]) |
| Gemini 4 Argon | Multistep pipeline | 3.2% (8/250; 95% CI [0.8, 6.0]) |
| Gemini 4 Argon | Omega Point | 2.0% (5/250; 95% CI [0.0, 4.4]) |

#### Appendix D. Full metrics for the prompt comparison

For each pair of approaches, four forecast metrics (positive predictive value, PPV; negative predictive value, NPV; sensitivity; specificity) were calculated on the same hypotheses; these shared sets are the same as in Table 4. All metrics are given with 95% CIs. A hypothesis that did not match the reference finding counted as a negative outcome.

| Predictor and prompt | PPV | NPV | Sensitivity | Specificity |
|---|---|---|---|---|
| Claude Sonnet 5.5; the detailed prompt | 20.6% (26/126; 95% CI [13.2, 28.9]) | 91.6% (109/119; 95% CI [86.0, 96.6]) | 72.2% (26/36; 95% CI [56.8, 87.5]) | 52.2% (109/209; 95% CI [46.3, 58.1]) |
| Claude Sonnet 5.5; the short prompt | 28.6% (14/49; 95% CI [16.7, 40.9]) | 88.8% (174/196; 95% CI [84.0, 93.3]) | 38.9% (14/36; 95% CI [27.0, 52.0]) | 83.3% (174/209; 95% CI [80.5, 86.1]) |
| GPT-6 Astra; the detailed prompt | 16.0% (32/200; 95% CI [10.1, 22.6]) | 92.0% (46/50; 95% CI [85.2, 98.0]) | 88.9% (32/36; 95% CI [75.8, 97.6]) | 21.5% (46/214; 95% CI [16.5, 26.9]) |
| GPT-6 Astra; the short prompt | 30.7% (23/75; 95% CI [20.0, 41.8]) | 92.6% (162/175; 95% CI [87.9, 96.6]) | 63.9% (23/36; 95% CI [48.8, 79.3]) | 75.7% (162/214; 95% CI [70.7, 80.5]) |
| Gemini 4 Argon; the detailed prompt | 21.1% (32/152; 95% CI [13.4, 29.6]) | 95.9% (94/98; 95% CI [91.8, 99.0]) | 88.9% (32/36; 95% CI [77.8, 97.5]) | 43.9% (94/214; 95% CI [38.6, 49.3]) |
| Gemini 4 Argon; the short prompt | 33.3% (21/63; 95% CI [21.8, 45.6]) | 92.0% (172/187; 95% CI [87.2, 96.2]) | 58.3% (21/36; 95% CI [43.2, 75.0]) | 80.4% (172/214; 95% CI [74.9, 85.0]) |
| Grok 4.5; the detailed prompt | 19.7% (27/137; 95% CI [12.2, 27.9]) | 95.5% (84/88; 95% CI [90.7, 98.9]) | 87.1% (27/31; 95% CI [73.3, 97.1]) | 43.3% (84/194; 95% CI [39.1, 47.5]) |
| Grok 4.5; the short prompt | 23.5% (20/85; 95% CI [14.1, 34.0]) | 92.1% (129/140; 95% CI [87.4, 96.3]) | 64.5% (20/31; 95% CI [48.1, 80.0]) | 66.5% (129/194; 95% CI [59.8, 73.0]) |

In an extended run, further Claude versions also received the detailed prompt. Each predictor's metrics are calculated over all its usable data, so the denominators may differ from those in the paired comparison above, and the intervals, calculated in a separate resampling, may differ in the last digit. The table lists the predictors that returned usable responses.

| Predictor | Usable hypotheses | PPV of recommendations (95% CI) | NPV of negative recommendations (95% CI) |
|---|---|---|---|
| Claude Sonnet 5.5 | 250 | 20.3% (26/128; 95% CI [12.8, 28.6]) | 91.8% (112/122; 95% CI [86.3, 96.7]) |
| Claude Opus 5.5 | 215 | 21.4% (22/103; 95% CI [12.0, 31.8]) | 93.8% (105/112; 95% CI [89.2, 97.5]) |
| Claude Haiku 5.5 | 250 | 18.3% (20/109; 95% CI [9.6, 28.1]) | 88.7% (125/141; 95% CI [83.2, 93.7]) |
| Claude Opus 4.8 | 240 | 20.9% (28/134; 95% CI [13.4, 29.0]) | 94.3% (100/106; 95% CI [89.1, 99.0]) |
| GPT-6 Astra | 250 | 16.0% (32/200; 95% CI [10.0, 22.7]) | 92.0% (46/50; 95% CI [85.2, 98.0]) |
| Gemini 4 Argon | 250 | 21.1% (32/152; 95% CI [13.5, 29.5]) | 95.9% (94/98; 95% CI [91.8, 99.0]) |
| Grok 4.5 | 225 | 19.7% (27/137; 95% CI [12.1, 27.9]) | 95.5% (84/88; 95% CI [90.7, 98.9]) |

#### Appendix E. Forecasting on the full corpus of 2997 hypotheses

The outcome was determined for each hypothesis regardless of how it was generated: it was positive if a majority of the judges scored the hypothesis 3 or higher. The self-forecast row gives the forecast of the model that generated the hypothesis. The majority and unanimity rules combined the self-forecast with the forecasts of all external predictors available for that hypothesis, whose number depended on the generator. For the manuscript, the intervals were recalculated by resampling the 50 tasks, keeping all hypotheses of each sampled task together. These descriptive results come from the internal development of the rule.

| Forecast rule on 2997 hypotheses | PPV (95% CI) | Sensitivity (95% CI) |
|---|---|---|
| The random control | 12.1% (179/1477; 95% CI [8.5, 16.1]) | 50.0% (179/358; 95% CI [44.1, 55.4]) |
| Generator forecasts its own hypotheses | 26.8% (164/611; 95% CI [18.5, 35.6]) | 45.8% (164/358; 95% CI [38.7, 52.4]) |
| Majority of predictors forecast "yes" | 37.8% (123/325; 95% CI [26.5, 49.2]) | 34.4% (123/358; 95% CI [28.5, 39.5]) |
| All predictors forecast "yes" | 50.0% (55/110; 95% CI [34.0, 64.2]) | 15.4% (55/358; 95% CI [10.5, 20.3]) |

#### Appendix F. Full intervals for four-model unanimity

The sets and outcome definitions are those of Table 6. Intervals come from 10,000 joint resamples of whole tasks (random seed 20261026). The predictor panel and the random control were analysed with identical formulas and denominators.

| Generator and rule | PPV | NPV | Sensitivity | Specificity |
|---|---|---|---|---|
| Grok 4.5; four-model unanimity | 73.7% (14/19; 95% CI [52.6, 93.3]) | 88.2% (195/221; 95% CI [83.4, 92.7]) | 35.0% (14/40; 95% CI [25.0, 46.2]) | 97.5% (195/200; 95% CI [95.3, 99.5]) |
| Grok 4.5; the random control | 17.9% (22/123; 95% CI [10.2, 26.4]) | 84.6% (99/117; 95% CI [78.0, 90.6]) | 55.0% (22/40; 95% CI [40.0, 69.2]) | 49.5% (99/200; 95% CI [43.1, 56.1]) |
| Gemini 4 Argon; four-model unanimity | 47.4% (9/19; 95% CI [25.0, 70.6]) | 89.6% (207/231; 95% CI [85.9, 93.1]) | 27.3% (9/33; 95% CI [13.6, 42.9]) | 95.4% (207/217; 95% CI [92.8, 97.7]) |
| Gemini 4 Argon; the random control | 12.1% (16/132; 95% CI [7.0, 17.7]) | 85.6% (101/118; 95% CI [78.7, 91.8]) | 48.5% (16/33; 95% CI [30.6, 66.7]) | 46.5% (101/217; 95% CI [39.4, 53.7]) |
| Claude Opus 5.5; four-model unanimity | 63.6% (14/22; 95% CI [42.9, 82.4]) | 86.4% (197/228; 95% CI [81.7, 90.8]) | 31.1% (14/45; 95% CI [18.4, 43.9]) | 96.1% (197/205; 95% CI [93.4, 98.5]) |
| Claude Opus 5.5; the random control | 16.8% (20/119; 95% CI [9.7, 24.6]) | 80.9% (106/131; 95% CI [74.5, 87.2]) | 44.4% (20/45; 95% CI [28.9, 58.8]) | 51.7% (106/205; 95% CI [44.9, 58.5]) |
| Claude Opus 5.5, repeat generation; four-model unanimity | 58.8% (10/17; 95% CI [35.0, 81.8]) | 85.5% (195/228; 95% CI [80.2, 90.3]) | 23.3% (10/43; 95% CI [12.5, 34.7]) | 96.5% (195/202; 95% CI [94.0, 98.6]) |
| Claude Opus 5.5, repeat generation; the random control | 18.8% (25/133; 95% CI [11.7, 26.8]) | 83.9% (94/112; 95% CI [77.5, 90.0]) | 58.1% (25/43; 95% CI [44.1, 71.1]) | 46.5% (94/202; 95% CI [40.4, 52.6]) |

#### Appendix G. Equal-length lists drawn from single-model forecasts

This calculation asks how many matching hypotheses would remain if a single model's recommendations were cut at random to the length of the predictor panel's list. For example, on the Grok hypotheses GPT-6 Astra made 53 positive forecasts, 25 of which match the reference finding. A random 19 of these 53 contain on average 8.96 matching hypotheses (47.2% of the list), with a central 95% range of 6 to 12. The predictor panel selected 19 hypotheses, 14 of them matching (73.7%; Appendix M.6).

For each generator and model, the table gives the pool of positive forecasts, the length of the random list, and its expected result. All hypotheses in a pool are equally likely to be drawn, and none is drawn twice. The "Full set" row repeats the baseline random control of Table M.6. Ranges come from the exact distribution, with hypotheses and judge scores held fixed. These additional calculations describe how the previously chosen rule performs on the same data.

| Generator | Pool from which hypotheses are drawn | Matching the reference finding / total in pool | Hypotheses in random list | Random selection, mean number matching the reference finding | Random selection, central 95% range of the number matching the reference finding |
|---|---|---|---|---|---|
| Grok 4.5 | Full set | 40/240 | 19 | 3.17 out of 19 (16.7%) | 0 to 6 |
| Grok 4.5 | "Yes" forecasts from Claude Sonnet 5.5 | 21/46 | 19 | 8.67 out of 19 (45.7%) | 5 to 12 |
| Grok 4.5 | "Yes" forecasts from GPT-6 Astra | 25/53 | 19 | 8.96 out of 19 (47.2%) | 6 to 12 |
| Grok 4.5 | "Yes" forecasts from Gemini 4 Argon | 23/52 | 19 | 8.40 out of 19 (44.2%) | 5 to 12 |
| Grok 4.5 | "Yes" forecasts from Grok 4.5 | 31/110 | 19 | 5.35 out of 19 (28.2%) | 2 to 9 |
| Gemini 4 Argon | Full set | 33/250 | 19 | 2.51 out of 19 (13.2%) | 0 to 6 |
| Gemini 4 Argon | "Yes" forecasts from Claude Sonnet 5.5 | 12/45 | 19 | 5.07 out of 19 (26.7%) | 2 to 8 |
| Gemini 4 Argon | "Yes" forecasts from GPT-6 Astra | 14/51 | 19 | 5.22 out of 19 (27.5%) | 2 to 8 |
| Gemini 4 Argon | "Yes" forecasts from Gemini 4 Argon | 18/53 | 19 | 6.45 out of 19 (34.0%) | 3 to 10 |
| Gemini 4 Argon | "Yes" forecasts from Grok 4.5 | 20/105 | 19 | 3.62 out of 19 (19.0%) | 1 to 7 |
| Claude Opus 5.5 | Full set | 45/250 | 22 | 3.96 out of 22 (18.0%) | 1 to 8 |
| Claude Opus 5.5 | "Yes" forecasts from Claude Sonnet 5.5 | 19/50 | 22 | 8.36 out of 22 (38.0%) | 5 to 12 |
| Claude Opus 5.5 | "Yes" forecasts from GPT-6 Astra | 21/49 | 22 | 9.43 out of 22 (42.9%) | 6 to 13 |
| Claude Opus 5.5 | "Yes" forecasts from Gemini 4 Argon | 22/56 | 22 | 8.64 out of 22 (39.3%) | 5 to 12 |
| Claude Opus 5.5 | "Yes" forecasts from Grok 4.5 | 26/110 | 22 | 5.20 out of 22 (23.6%) | 2 to 9 |

#### Appendix H. Proportion of correct forecasts in the 50/50 set

The original protocol specified balanced accuracy, the mean of sensitivity and specificity, as the primary metric. Because the set contains equal numbers of confirmed and refuted statements, balanced accuracy here equals the proportion of correct forecasts. The nominal 95% CIs are taken from the report and come from 10,000 resamples of articles, each resampled with both its statements. Under unanimity, a statement receives a final "yes" only if all five forecasts are positive; every other statement receives a final "no". A high proportion of confirmed statements among those selected unanimously therefore comes with many missed confirmed statements and a lower overall proportion of correct forecasts.

| Model or rule | Proportion of correct forecasts on the 50/50 set (95% CI) |
|---|---|
| Claude Sonnet 5.5 | 74.0% (95% CI [66.0, 82.0]) |
| GPT-6 Astra | 71.0% (95% CI [63.0, 79.0]) |
| Gemini 4 Argon | 78.0% (95% CI [69.0, 86.0]) |
| Grok 4.5 | 75.0% (95% CI [65.0, 84.0]) |
| Claude Opus 5.5 | 73.0% (95% CI [66.0, 80.0]) |
| Five-model unanimity | 70.0% (95% CI [63.0, 77.0]) |
| The random control | 52.0% (95% CI [43.0, 61.0]) |

#### Appendix I. Search query and publication selection log

The query is preserved in the original export of 13 October 2026. The edat field gives the date a record entered PubMed; each article's eligibility was also checked against the date of its first public version and against any preprints.

```
(aging[MeSH Terms] OR ageing[tiab] OR aging[tiab] OR longevity[tiab] OR lifespan[tiab] OR senescence[tiab] OR healthspan[tiab]) AND ("Nature"[ta] OR "Science"[ta] OR "Cell"[ta] OR "Nat Aging"[ta] OR "Cell Metab"[ta] OR "Nat Med"[ta] OR "Nat Cell Biol"[ta] OR "Cell Stem Cell"[ta] OR "Aging Cell"[ta] OR "Nat Commun"[ta] OR "Sci Adv"[ta] OR "Proc Natl Acad Sci U S A"[ta] OR "Mol Cell"[ta] OR "Immunity"[ta] OR "Neuron"[ta]) AND journal article[pt] NOT review[pt] NOT clinical trial[pt] AND ("2026/08/01"[edat] : "2026/10/12"[edat])
```

The recorded sequence comprises 284 retrieved records, 72 exclusions at the preliminary filter, and 212 provisionally eligible candidates; the final set has 50 tasks. The extraction log keeps only the last run for each article, because earlier runs were overwritten when articles were replaced; its count of records with the status "accepted" is therefore not a stage of the selection in Figure 2. The selection log gives the identifiers of the final tasks, the order of the 212 candidates, the reasons for the 72 preliminary exclusions, and the checksums of the source files. Selecting tasks for suitability and for a concealed reference finding limits how representative the set is.

#### Appendix J. Agreement among judges in binary votes on matching the reference finding

A judge's vote was positive if the judge's score reached the threshold: 3 for the primary threshold and 4 for the strict threshold. The first table compares the four judges pairwise across all 2997 generated hypotheses. These comparisons include the scores of the judge from the generator's developer, which were excluded when the outcome was determined. "First" and "second" judge in the column headers follow the order of the names in the first column. Hypothesis counts and the proportion of agreeing votes refer to the vote "score of 3 or higher". Cohen's kappa is 1 for complete agreement and 0 for the agreement expected from independent votes with each judge's observed proportion of positive votes. Quadratically weighted kappa compares scores across the full 0 to 5 scale. The 95% confidence intervals come from 10,000 resamples of the 50 tasks.

| Judge pair | Hypotheses with a score of 3 or higher from both judges | Hypotheses with a score of 3 or higher from the first judge only | Hypotheses with a score of 3 or higher from the second judge only | Hypotheses with a score below 3 from both judges | Proportion of hypotheses with agreeing votes from the two judges, % | Cohen's kappa for the vote "score of 3 or higher" (95% CI) | Cohen's kappa for the vote "score of 4 or 5" (95% CI) | Weighted kappa across the full 0 to 5 scale |
|---|---|---|---|---|---|---|---|---|
| GPT-5.6 Sol and Grok 4.7 | 325 | 155 | 61 | 2456 | 92.8 | 0.71 (95% CI [0.66, 0.76]) | 0.67 (95% CI [0.50, 0.81]) | 0.68 |
| GPT-5.6 Sol and Claude Opus 5.5 | 314 | 166 | 49 | 2468 | 92.8 | 0.70 (95% CI [0.65, 0.76]) | 0.54 (95% CI [0.39, 0.65]) | 0.45 |
| GPT-5.6 Sol and Gemini 4 Argon | 310 | 170 | 40 | 2477 | 93.0 | 0.71 (95% CI [0.64, 0.77]) | 0.61 (95% CI [0.45, 0.73]) | 0.62 |
| Grok 4.7 and Claude Opus 5.5 | 299 | 87 | 64 | 2547 | 95.0 | 0.77 (95% CI [0.73, 0.81]) | 0.47 (95% CI [0.28, 0.60]) | 0.38 |
| Grok 4.7 and Gemini 4 Argon | 281 | 105 | 69 | 2542 | 94.2 | 0.73 (95% CI [0.67, 0.78]) | 0.72 (95% CI [0.58, 0.80]) | 0.52 |
| Claude Opus 5.5 and Gemini 4 Argon | 275 | 88 | 75 | 2559 | 94.6 | 0.74 (95% CI [0.69, 0.78]) | 0.60 (95% CI [0.44, 0.71]) | 0.64 |

The second table includes, for each hypothesis, only the three judges from other developers, whose votes determined its outcome. The outcome is positive if two or three judges gave a score in the range of the first column. Fleiss' kappa generalises Cohen's kappa to three judges.

| Judge score | Hypotheses with this score from all three judges | Hypotheses with this score from two of the three judges | Hypotheses with this score from one of the three judges | Hypotheses with this score from none of the judges | Fleiss' kappa for three judges (95% CI) |
|---|---|---|---|---|---|
| 3 or higher | 263 | 95 | 188 | 2451 | 0.72 (95% CI [0.67, 0.77]) |
| 4 or 5 | 36 | 21 | 73 | 2867 | 0.57 (95% CI [0.41, 0.67]) |

#### Appendix K. Additional calculations for selection and forecasts

These calculations use the recorded run reports and make no new model calls. Intervals come from 10,000 resamples of whole tasks (random seed 20261030). The methods are described in Section 2.10 and Appendix L.6.

##### K.1. Ranking hypotheses by the numerical probability of confirmation

This calculation uses the 250 hypotheses that GPT-6 Astra generated in a single request and the probability each predictor assigned to them under the detailed prompt. The area under the ROC curve (AUC) is the probability that a hypothesis matching the reference finding receives a higher score than a non-matching one, with ties counted as 0.5. Top-k precision is the expected number of matches divided by the list length; when probabilities are tied at the list boundary, the expectation uses the match rate within the tied group. The baseline match rate is 14.4% (36/250).

| Predictor | AUC (95% CI) | Top-19 precision | Top-20 precision | Top-21 precision | Top-22 precision |
|---|---|---|---|---|---|
| Claude Sonnet 5.5 | 0.71 (95% CI [0.62, 0.79]) | 38.9% (7.4/19) | 38.0% (7.6/20) | 37.1% (7.8/21) | 36.4% (8.0/22) |
| Claude Opus 5.5 | 0.71 (95% CI [0.60, 0.79]) | 36.8% (7.0/19) | 36.7% (7.3/20) | 36.5% (7.7/21) | 36.4% (8.0/22) |
| Claude Haiku 5.5 | 0.62 (95% CI [0.50, 0.72]) | 27.6% (5.2/19) | 27.5% (5.5/20) | 27.4% (5.8/21) | 27.3% (6.0/22) |
| Claude Opus 4.8 | 0.67 (95% CI [0.56, 0.77]) | 33.3% (6.3/19) | 32.5% (6.5/20) | 31.7% (6.7/21) | 31.1% (6.8/22) |
| GPT-6 Astra | 0.68 (95% CI [0.57, 0.79]) | 42.1% (8.0/19) | 40.0% (8.0/20) | 39.3% (8.2/21) | 38.6% (8.5/22) |
| Gemini 4 Argon | 0.74 (95% CI [0.64, 0.82]) | 35.7% (6.8/19) | 35.0% (7.0/20) | 35.7% (7.5/21) | 36.4% (8.0/22) |
| Grok 4.5 | 0.71 (95% CI [0.60, 0.81]) | 36.8% (7.0/19) | 35.0% (7.0/20) | 33.3% (7.0/21) | 31.8% (7.0/22) |

Unanimous selection by the four predictors was applied to the same hypotheses, using the four predictors' recorded answers to the short prompt. All four predictors answered for 245 hypotheses in 49 tasks. The rule selected 22 of them, and 50.0% of the selected hypotheses matched the reference finding (11/22; 95% CI [30.0, 70.4]) against a baseline match rate of 14.7% (36/245), a ratio of 3.40 (95% CI [2.26, 4.95]). The table below compares this list with the 22 hypotheses ranked highest by probability among the same 245. Hypotheses without a numerical probability were excluded from the ranking. The difference is the match rate among unanimously selected hypotheses minus the match rate among the top-ranked hypotheses; the $p$-value comes from permuting the two lists within tasks, with Holm correction across the seven predictors.

| Predictor whose probability ranked the hypotheses | Hypotheses with a numerical probability | Match rate among the top 22 | Ratio to the baseline match rate (95% CI) | Unanimity minus ranking, percentage points (95% CI) | Holm-adjusted *p* |
|---|---|---|---|---|---|
| Claude Sonnet 5.5 | 245 | 36.4% (8.0/22) | 2.47 (95% CI [1.29, 4.00]) | 13.6 (95% CI [−9.1, 36.7]) | 0.6573 |
| Claude Opus 5.5 | 210 | 36.4% (8.0/22) | 2.47 (95% CI [1.25, 3.98]) | 13.6 (95% CI [−7.5, 34.2]) | 0.6573 |
| Claude Haiku 5.5 | 245 | 27.3% (6.0/22) | 1.86 (95% CI [0.70, 3.20]) | 22.7 (95% CI [2.2, 45.2]) | 0.3031 |
| Claude Opus 4.8 | 235 | 31.1% (6.8/22) | 2.11 (95% CI [0.82, 3.54]) | 18.9 (95% CI [−2.9, 42.5]) | 0.6573 |
| GPT-6 Astra | 240 | 38.6% (8.5/22) | 2.63 (95% CI [1.01, 4.28]) | 11.4 (95% CI [−11.1, 36.1]) | 0.6573 |
| Gemini 4 Argon | 245 | 36.4% (8.0/22) | 2.47 (95% CI [1.15, 4.08]) | 13.6 (95% CI [−2.6, 30.4]) | 0.6573 |
| Grok 4.5 | 220 | 31.8% (7.0/22) | 2.17 (95% CI [0.91, 3.88]) | 18.2 (95% CI [−4.0, 40.0]) | 0.6573 |

##### K.2. Predictor panel versus the best single model at equal list length

The best single model is the predictor with the highest match rate among its positive forecasts. Shortening its list at random to the panel's list length leaves the expected match rate unchanged, so the rates are compared directly. The *p*-value comes from permuting the two rules within each task, with Holm correction across the four sets. The rule was chosen on the Grok hypotheses, and both rules use the same forecasts.

| Hypothesis set | Match rate among hypotheses selected by the panel | Best single model | Match rate among its positive forecasts | Difference, percentage points (95% CI) | Holm-adjusted *p* |
|---|---|---|---|---|---|
| Grok 4.5 | 73.7% (14/19) | GPT-6 Astra | 47.2% (25/53) | 26.5 (95% CI [8.2, 45.9]) | $p < 0.05$ |
| Gemini 4 Argon | 47.4% (9/19) | Gemini 4 Argon | 34.0% (18/53) | 13.4 (95% CI [−4.3, 31.5]) | 0.0833 |
| Claude Opus 5.5 | 63.6% (14/22) | GPT-6 Astra | 42.9% (21/49) | 20.8 (95% CI [6.8, 35.6]) | $p < 0.05$ |
| Claude Opus 5.5, repeat generation | 58.8% (10/17) | Gemini 4 Argon | 34.4% (21/61) | 24.4 (95% CI [6.0, 44.7]) | $p < 0.01$ |

##### K.3. Selection without the predictor from the generator's developer

The predictor from the generator's developer was removed from the panel, and the three remaining predictors still had to agree unanimously. The last column repeats the full-panel result of Table 6.

| Hypothesis set | Excluded predictor | Match rate among selected hypotheses (95% CI) | Proportion of all matching hypotheses found | Match rate with the full panel |
|---|---|---|---|---|
| Grok 4.5 | Grok 4.5 | 68.2% (15/22; 95% CI [47.6, 86.7]) | 37.5% (15/40) | 73.7% (14/19) |
| Gemini 4 Argon | Gemini 4 Argon | 45.0% (9/20; 95% CI [23.1, 66.7]) | 27.3% (9/33) | 47.4% (9/19) |
| Claude Opus 5.5 | Claude Sonnet 5.5 | 53.6% (15/28; 95% CI [35.0, 70.8]) | 33.3% (15/45) | 63.6% (14/22) |
| Claude Opus 5.5, repeat generation | Claude Sonnet 5.5 | 46.4% (13/28; 95% CI [26.1, 65.6]) | 30.2% (13/43) | 58.8% (10/17) |

##### K.4. Decisions within pairs in the 50/50 set

Each article in the set contributed one confirmed and one refuted statement. The columns give the four possible decisions on a pair, and together they sum to the number of pairs.

| Model or rule | Article pairs | Confirmed accepted, reversed rejected | Both statements accepted | Both statements rejected | Reversed accepted, confirmed rejected |
|---|---|---|---|---|---|
| Claude Sonnet 5.5 | 50 | 26 (52.0%) | 13 | 9 | 2 |
| GPT-6 Astra | 50 | 23 (46.0%) | 18 | 7 | 2 |
| Gemini 4 Argon | 50 | 31 (62.0%) | 6 | 10 | 3 |
| Grok 4.5 | 50 | 30 (60.0%) | 6 | 9 | 5 |
| Claude Opus 5.5 | 50 | 24 (48.0%) | 18 | 7 | 1 |
| Random control | 50 | 12 (24.0%) | 15 | 13 | 10 |

The “effect in the favourable direction” rule labelled 61 of the 100 statements. The proportion of correct answers is reported separately for labelled and unlabelled statements.

| Model or rule | Proportion of correct answers among the 61 labelled statements | Proportion of correct answers among the 39 unlabelled statements |
|---|---|---|
| Claude Sonnet 5.5 | 68.9% (42/61) | 82.1% (32/39) |
| GPT-6 Astra | 77.0% (47/61) | 61.5% (24/39) |
| Gemini 4 Argon | 75.4% (46/61) | 82.1% (32/39) |
| Grok 4.5 | 75.4% (46/61) | 74.4% (29/39) |
| Claude Opus 5.5 | 70.5% (43/61) | 76.9% (30/39) |
| Random control | 52.5% (32/61) | 51.3% (20/39) |
| “Effect in the favourable direction” rule | 65.6% (40/61) | No label assigned |

The rule labelled both statements in 20 pairs. The table below shows the decisions of the rule, the models, and the random control on these pairs; the rule accepts a statement labelled “favourable” and rejects one labelled “unfavourable”.

| Model or rule | Article pairs | Confirmed accepted, reversed rejected | Both statements accepted | Both statements rejected | Reversed accepted, confirmed rejected |
|---|---|---|---|---|---|
| “Effect in the favourable direction” rule | 20 | 5 | 13 | 0 | 2 |
| Claude Sonnet 5.5 | 20 | 9 | 5 | 4 | 2 |
| GPT-6 Astra | 20 | 11 | 5 | 4 | 0 |
| Gemini 4 Argon | 20 | 11 | 3 | 5 | 1 |
| Grok 4.5 | 20 | 13 | 2 | 4 | 1 |
| Claude Opus 5.5 | 20 | 8 | 7 | 4 | 1 |
| Random control | 20 | 5 | 6 | 6 | 3 |

##### K.5. Enrichment under the strict threshold

The strict threshold requires a score of 4 or 5 from a majority of the judges. The ratio is the match rate among selected hypotheses divided by the baseline match rate at the same threshold; the selected lists are unchanged.

| Hypothesis set | Match rate among selected hypotheses | Baseline match rate | Ratio of these rates (95% CI) | Proportion of all matching hypotheses found |
|---|---|---|---|---|
| Grok 4.5 | 26.3% (5/19) | 2.9% (7/240) | 9.02 (95% CI [4.21, 16.00]) | 71.4% (5/7) |
| Gemini 4 Argon | 31.6% (6/19) | 3.6% (9/250) | 8.77 (95% CI [5.18, 14.71]) | 66.7% (6/9) |
| Claude Opus 5.5 | 9.1% (2/22) | 2.0% (5/250) | 4.55 (95% CI [0.00, 11.03]) | 40.0% (2/5) |
| Claude Opus 5.5, repeat generation | 11.8% (2/17) | 1.2% (3/245) | 9.61 (95% CI [0.00, 18.85]) | 66.7% (2/3) |

#### K.6. Originality, feasibility, and clarity of selected and excluded hypotheses

Each selected hypothesis was paired with an excluded hypothesis from the same task. A hypothesis's score is the mean of the three judges from other developers on the 1 to 10 scale. Differences are selected minus excluded; intervals come from 10,000 task resamples.

| Hypothesis set | Metric | Hypothesis pairs | Selected, mean score | Excluded, mean score | Difference, selected minus excluded (95% CI) | Pairs with a higher score for the selected hypothesis | Pairs with a lower score for the selected hypothesis | Pairs with equal scores |
|---|---|---|---|---|---|---|---|---|
| Grok 4.5 | Originality | 19 | 4.49 | 4.72 | −0.23 (95% CI [−0.75, 0.32]) | 7 | 11 | 1 |
| Grok 4.5 | Feasibility | 19 | 6.81 | 6.46 | 0.35 (95% CI [−0.35, 1.05]) | 8 | 6 | 5 |
| Grok 4.5 | Clarity | 19 | 8.14 | 7.95 | 0.19 (95% CI [−0.04, 0.46]) | 8 | 5 | 6 |
| Gemini 4 Argon | Originality | 19 | 4.82 | 5.00 | −0.18 (95% CI [−0.58, 0.26]) | 6 | 10 | 3 |
| Gemini 4 Argon | Feasibility | 19 | 6.70 | 6.02 | 0.68 (95% CI [0.39, 0.98]) | 13 | 1 | 5 |
| Gemini 4 Argon | Clarity | 19 | 8.04 | 7.26 | 0.77 (95% CI [0.51, 1.05]) | 16 | 0 | 3 |
| Claude Opus 5.5 | Originality | 21 | 5.24 | 5.48 | −0.24 (95% CI [−0.62, 0.13]) | 8 | 10 | 3 |
| Claude Opus 5.5 | Feasibility | 21 | 6.59 | 6.25 | 0.33 (95% CI [−0.02, 0.72]) | 11 | 6 | 4 |
| Claude Opus 5.5 | Clarity | 21 | 8.25 | 8.06 | 0.19 (95% CI [0.05, 0.33]) | 8 | 2 | 11 |
| All three sets combined | Originality | 59 | 4.86 | 5.08 | −0.21 (95% CI [−0.47, 0.04]) | 21 | 31 | 7 |
| All three sets combined | Feasibility | 59 | 6.69 | 6.24 | 0.45 (95% CI [0.17, 0.73]) | 32 | 13 | 14 |
| All three sets combined | Clarity | 59 | 8.15 | 7.77 | 0.38 (95% CI [0.24, 0.53]) | 32 | 7 | 20 |

Logistic regression on the 119 hypotheses in this comparison (60 selected and 59 excluded) estimated the association between selection and matching the reference finding after adjustment for clarity. The outcome is 1 if the hypothesis matches the reference finding at the primary threshold. Clarity and feasibility are the mean scores of the three judges, centred on their means across the 119 hypotheses. In total, 37 of the 60 selected and 15 of the 59 excluded hypotheses matched the reference finding. Wald intervals and $p$-values use standard errors clustered on the 58 generator-by-task combinations; a separate column gives the percentile interval from 10,000 resamples of these combinations.

| Variables in the regression | Variable | Odds ratio (95% CI) | Task-resample interval | *p* |
|---|---|---|---|---|
| Selection only | Selection (selected versus excluded) | 4.72 (95% CI [2.36, 9.42]) | [2.51, 10.30] | $p < 0.001$ |
| Selection and clarity | Selection (selected versus excluded) | 3.85 (95% CI [1.81, 8.20]) | [1.90, 9.04] | $p < 0.001$ |
| Selection and clarity | Clarity, per 1-point increase | 2.02 (95% CI [1.04, 3.95]) | [1.04, 4.42] | $p < 0.05$ |
| Selection, clarity, and feasibility | Selection (selected versus excluded) | 4.03 (95% CI [1.89, 8.60]) | [1.96, 9.90] | $p < 0.001$ |
| Selection, clarity, and feasibility | Clarity, per 1-point increase | 2.14 (95% CI [1.05, 4.34]) | [1.06, 4.99] | $p < 0.05$ |
| Selection, clarity, and feasibility | Feasibility, per 1-point increase | 0.88 (95% CI [0.57, 1.36]) | [0.56, 1.48] | 0.5647 |
| Clarity only | Clarity, per 1-point increase | 2.75 (95% CI [1.56, 4.88]) | [1.62, 5.41] | $p < 0.001$ |

#### K.7. Repeat run of the forecasts

The comparison covers hypotheses that received forecasts in both runs. Cohen's kappa (κ) was calculated on each model's "yes" and "no" decisions. The match rate among hypotheses selected in the initial run is reported for the same hypotheses, so its denominator can differ from Table 6.

| Hypothesis set | Hypotheses with forecasts in both runs | Decision agreement, Claude Sonnet 5.5 | Decision agreement, GPT-6 Astra | Decision agreement, Gemini 4 Argon | Decision agreement, Grok 4.5 | Selected in the initial run | Selected in the repeat run | Selected in both runs | Match rate among hypotheses selected in the repeat run (95% CI) | Match rate among hypotheses selected in the initial run |
|---|---|---|---|---|---|---|---|---|---|---|
| Grok 4.5 | 235 | 93.2% (219/235; κ = 0.78) | 94.0% (221/235; κ = 0.83) | 96.2% (226/235; κ = 0.89) | 91.9% (216/235; κ = 0.84) | 18 | 19 | 16 | 68.4% (13/19; 95% CI [46.7, 88.9]) | 72.2% (13/18) |
| Gemini 4 Argon | 245 | 93.5% (229/245; κ = 0.78) | 95.1% (233/245; κ = 0.85) | 95.5% (234/245; κ = 0.86) | 88.6% (217/245; κ = 0.76) | 19 | 18 | 17 | 50.0% (9/18; 95% CI [26.7, 75.0]) | 47.4% (9/19) |
| Claude Opus 5.5 | 240 | 90.8% (218/240; κ = 0.71) | 96.2% (231/240; κ = 0.88) | 93.8% (225/240; κ = 0.83) | 88.3% (212/240; κ = 0.76) | 21 | 21 | 15 | 66.7% (14/21; 95% CI [43.5, 86.4]) | 61.9% (13/21) |

#### K.8. Lexical check of key elements

An element of the reference finding counted as named if at least two-thirds of its content words appeared in the hypothesis text. The calculation covers the 1000 single-request hypotheses. The AUC measures how well the share of named elements separates hypotheses that the judges scored as matching the reference finding from those they did not; its interval comes from 10,000 task resamples. Among the 60 hypotheses selected unanimously by the four predictors, matching hypotheses named on average 17.3% of the elements and the remaining hypotheses 6.1%; at least two elements were named in 6 of the 37 matching hypotheses and in 2 of the 23 others. Among all 1000 hypotheses, at least one element was named in 46.2% (72/156) of matching and 16.4% (138/844) of non-matching hypotheses (odds ratio 4.39, 95% CI [2.87, 6.81]).

| Generator | Hypotheses | Matching the reference finding | AUC of the share of named elements |
|---|---|---|---|
| Claude Opus 5.5 | 250 | 45 | 0.62 |
| Gemini 4 Argon | 250 | 33 | 0.76 |
| GPT-6 Astra | 250 | 36 | 0.63 |
| Grok 4.5 | 250 | 42 | 0.63 |
| All four models combined | 1000 | 156 | 0.65 (95% CI [0.60, 0.70]) |

#### Appendix L. Additional methodological details

##### L.1. Model run settings

The token limit applies to one model call and covers both reasoning and the response. Response attempts give how many times a request was repeated when the response could not be parsed or the model refused to answer.

| Role | Models | System prompt | Token limit per call | Response attempts | Tools and internet access | Reasoning setting |
|---|---|---|---|---|---|---|
| Hypothesis generators | GPT-6 Astra, Grok 4.5, Claude Opus 5.5, Gemini 4 Argon | the task, without the reference finding or key elements | 32,000 | up to three per single request and up to three per pipeline step | disabled; not confirmed per request for the original generations (Section 2.9) | high reasoning effort |
| Judges | GPT-5.6 Sol, Grok 4.7, Claude Opus 5.5, Gemini 4 Argon | the reference finding, the key elements, and the matching scale | 32,000 | up to three | disabled; not confirmed per request | `reasoning = medium` |
| Predictors, detailed prompt | Claude Sonnet 5.5, GPT-6 Astra, Gemini 4 Argon, Grok 4.5; in the extended run, also Claude Opus 5.5, Claude Haiku 5.5, and Claude Opus 4.8 | scientific criteria and an instruction to return JSON (Appendix B.1) | 32,000 | two | disabled; confirmed from logs for the runs of 22 to 24 October | high reasoning effort |
| Predictors, short prompt | Claude Sonnet 5.5, GPT-6 Astra, Gemini 4 Argon, Grok 4.5, Claude Opus 5.5 | an instruction to return JSON (Appendices B.2 and B.3) | 32,000 | two | disabled; confirmed from logs | high reasoning effort |

##### L.2. Vote distribution and judge agreement across the full scale

All three judges scored 263 hypotheses 3 or higher; two of the three judges did so for 95 hypotheses, one judge for 188, and no judge for 2451. A single changed vote would flip the majority decision for 283 hypotheses: 95 were classified as matching by two votes of three, and 188 as non-matching with one vote in favour. Across individual judge pairs, the bounds of the 95% CI for Cohen's kappa on the "score of 3 or higher" vote ranged from 0.64 to 0.81.

Quadratically weighted kappa counts disagreements across the whole 0 to 5 scale, including those within the 0 to 2 and 3 to 5 ranges, which do not change the outcome at the primary threshold. For every judge pair it was lower than Cohen's kappa for the "score of 3 or higher" vote, ranging from 0.38 to 0.68; Spearman's rank correlation between scores ranged from 0.57 to 0.66. Appendix J gives the weighted kappa for each judge pair.

##### L.3. The random control and equal-length random lists

Random answers were generated with the Node.js `crypto.randomInt` function for the 2997-hypothesis set, the unanimous-selection sets, and the Claude Opus 5.5 repeat generation, and with `crypto.randomBytes` for the 250 GPT-6 Astra hypotheses and the 50/50 set. Forecasting on the full corpus of 2997 hypotheses used a separate recorded sequence with 1477 "yes" answers.

To control for list length, we considered drawing, with equal probability and without replacement, exactly as many distinct hypotheses as the predictor panel selected: 19 of the 240 Grok hypotheses (40 of which match the reference finding), 19 of the 250 Gemini hypotheses, and 22 of the 250 Opus hypotheses. The program computed the exact hypergeometric distribution of the number of matches in such a list, its mean, and its central 95% range; the mean was also checked against 100,000 random draws from this distribution. The calculation needs only the set size, the number of matching hypotheses in the set, and the list length; the simulations did not sample hypothesis identifiers.

For comparison with a single model, that model's positive forecasts were randomly subsampled to the panel's list length. For example, GPT-6 Astra answered "yes" for 53 Grok hypotheses, 25 of which match the reference finding; we calculated the result of drawing 19 of those 53 at random. This control was run for each of the four predictors and each generator. All hypotheses in the "yes" group had equal probability, because binary answers define only a "yes" group and a "no" group. Appendix G gives the results.

##### L.4. Enumeration of selection rules

The enumeration of 80 voting rules on the Grok hypotheses tested, for each combination of k models, all k thresholds on the number of "yes" answers: 5 rules for single models, 20 for pairs, 30 for triples, 20 for quadruples, and 5 for the full panel. The final four-model unanimity rule is one of these 80 rules.

In the 32 two-stage schemes, one "yes" answer was enough to pass the second stage. In 22 of them, the first-stage model was also on the second-stage panel, so its own "yes" already passed the second stage; these schemes duplicated selection by a single model, and the remaining 10 defined additional rules.

The sets and the rule for the Gemini 4 Argon and Claude Opus 5.5 hypotheses were fixed on 23 October 2026 at 21:43 and 22:09 UTC, respectively, before the first requests to the predictors.

##### L.5. Isolation of the initial Claude Opus 5.5 generation

The benchmark script requested that tools be disabled, but until a fix on 21 October the tool-disabling parameter was lost when requests were streamed. Request logs for 13 to 18 October were not preserved. On 27 October this configuration was reproduced with the same software version and settings, and the model was asked to open a web page, run a web search, and read the task file containing the reference findings. All five of the model's tool calls were rejected; no web search or page load took place. A search of the 685 saved raw responses of the initial generation and the 647 of the repeat generation found no URLs, DOIs, PubMed identifiers, mentions of the year 2026, preprint servers, or the DOI of the task article. The same search of the 2916 saved raw responses of the four generators also found none.

In the repeat generation of 27 to 28 October, each of the 792 successful requests was matched to the call log and to a saved record confirming that tools were disabled. The same three judges from other developers scored the hypotheses with the same prompt. If the Claude safety classifier cut off the response on every attempt and no hypotheses remained, the set was counted as a refusal, that is, as five non-matching hypotheses. The multistep pipeline set for task R42736449 was not run on 27 October: its number of generated tokens is matched to Omega Point on the same task, and the script did not recognise the Claude refusal in Omega Point. The set was generated on 28 October after the script was fixed.

##### L.6. Statistical procedures

**Random seeds.** Generation intervals used random seed 20261025; intervals for unanimous selection and for the random control on the same hypotheses (Table 6, Appendix F) used seed 20261026. These intervals replace those in the initial reports, which were based on 2000 resamples. The sensitivity analyses and the cueing analysis used seed 20261027, and the AUC intervals used 20261030. Generators were compared across the 48 shared tasks with 10,000 joint resamples.

**Family of 150 originality comparisons.** Five metrics were evaluated: originality, technical feasibility, clarity, distinctiveness of the two ideas within one response, and an overall score, the mean of those four ratings. For each metric, we compared three pairs of generation methods within each of the four models ($4 \times 3 \times 5 = 60$) and six pairs of models within each of the three methods ($6 \times 3 \times 5 = 90$). Each pairwise comparison used the judges common to both models, excluding the judges from both models' developers. Adjustments for individual judges and for all judges combined were calculated separately and are not part of the family of 150 comparisons.

**Sensitivity analyses.** The overall match rate in single-request generation and the match rate among selected hypotheses in the final selection were recalculated on the tasks with a cueing score of 0 or 1. The final selection was also re-evaluated leaving out one judge at a time; a positive outcome then required a score of 3 or higher from both remaining judges, and the selected lists were unchanged. Opus generation without the sets containing a Claude refusal was recalculated with the parameters of the primary analysis and Holm correction across the three within-model comparisons.

**Probability ranking.** For the 250 hypotheses that Astra generated in a single request, we calculated, for each predictor under the detailed prompt, the AUC of the confirmation probability and the match rate among the top k ranked hypotheses for k = 19 to 22, the lengths of the final unanimous-selection lists. When probabilities were tied at the list boundary, the expected number of matches was calculated from the match rate within the tied group. Unanimous selection by the four predictors was applied to their saved answers to the short prompt for 245 hypotheses in 49 tasks, and its list was compared with each model's top k ranked hypotheses, k being the length of the rule's list. The difference in match rates was tested by permuting the two lists within tasks, with Holm correction across the seven models.

**Comparison with the best single model.** The best model is the one with the highest match rate among its "yes" forecasts. Randomly subsampling its list to the panel's list length leaves the expected match rate among selected hypotheses unchanged, so the rates themselves were compared. Two-sided *p*-values come from permuting the two rules within each task, by exhaustive enumeration when few tasks differed and otherwise by 19,999 random permutations with one added to both numerator and denominator; Holm correction was applied across the four sets. The panel's list is a subset of its members' "yes" forecasts, so the two rules are dependent.

**Selection without the predictor from the generator's developer.** The predictor from the generator's developer was removed from the panel, and the three remaining predictors had to agree unanimously.

**Within-pair decisions on the 50/50 set.** For each article, we checked whether the model accepted the confirmed statement and rejected the refuted one. The "effect in the favourable direction" control is a deterministic rule applied to the statement text: the direction of change follows from the verb and from the sign that the measured indicator has for the organism. Statements about indicators with no agreed sign were left unlabelled, and the labelling did not use the experimental labels. Model accuracy was calculated separately for labelled and unlabelled statements.

**Enrichment at the strict threshold.** We divided the match rate among selected hypotheses by the baseline match rate at the same threshold; the interval comes from the same task resamples.

**Lexical check of key elements.** In the 1000 single-request hypotheses, an element of the reference finding counted as named if at least two-thirds of its content words appeared in the hypothesis text. The check detects shared words and does not count paraphrase. The association between the share of named elements and the judges' decision was measured by the AUC and by the odds ratio for naming at least one element; intervals come from 10,000 task resamples. This check does not replace the judges' evaluation.

**Repeat run of the forecasts.** On 30 October, the four predictors received requests byte-identical to those of 23 October: the cohorts, the random-control answers, and all 600 request items matched. Each request's isolation was verified against the call log and the records confirming that tools were disabled; one model ran on three sets concurrently, and the log was cross-checked with those concurrent runs taken into account. Repeatability was measured by the proportion of agreeing decisions, Cohen's kappa, and the number of hypotheses selected in both runs.

**Originality of selected and excluded hypotheses.** Each selected hypothesis was paired with one excluded hypothesis from the same task, chosen by hash. Three judges from other developers scored both with the prompt of the originality experiment: originality, feasibility, and clarity on a 1 to 10 scale, one hypothesis per request, with no information about selection. The three judges' scores were averaged; the "selected minus excluded" difference was evaluated with 10,000 task resamples and within-task sign permutation (19,999 permutations), with Holm correction across the three metrics. The association between selection and matching after adjustment for clarity was estimated by logistic regression of a match with the reference finding at the primary threshold on 119 hypotheses; the independent variables were membership in the selected set and the mean clarity score, and standard errors were clustered on the generator-by-task combinations.

**Degree of cueing.** For each task, the match rates of the four generators in single-request generation were averaged; the task was the unit of analysis (50 tasks, 1000 hypotheses). Two-sided $p$-values were based on the absolute correlation over 99,999 permutations of question scores between whole tasks, with one added to both numerator and denominator. Confidence intervals come from 10,000 resamples of whole tasks, with ranks recalculated in each. Exploratory checks included a joint rating of the research question and the background, other generation methods, alternative ratings of borderline questions, and exclusion of previously discussed examples. The annotations, data, and parameters are given in the report on cueing in the tasks; this analysis did not use model forecasts.

##### L.7. Order of candidate review

Candidates were ordered by the SHA-256 hash of the string `omega-creativity-2026-10-11` followed by the PubMed identifier. This ordering reproduces the saved order of the 212 candidates.

##### L.8. Definitions of measures

- overall match rate: the proportion of all hypotheses in a set that match the reference finding;
- positive predictive value (PPV): the proportion of matching hypotheses among those selected or forecast “yes” (in the 50/50 set, the proportion of confirmed statements);
- negative predictive value (NPV): the proportion of non-matching hypotheses among those forecast “no” (in the 50/50 set, the proportion of refuted statements);
- sensitivity: the proportion of matching hypotheses (confirmed statements) that were selected or forecast “yes”;
- specificity: the proportion of non-matching hypotheses (refuted statements) forecast “no”;
- balanced accuracy: the mean of sensitivity and specificity;
- area under the ROC curve (AUC): the probability that a randomly chosen matching hypothesis receives a higher score than a randomly chosen non-matching one.

#### Appendix M. Additional results

##### M.1. Claude Opus 5.5 repeat generation and Claude refusals

With single-request generation, the match rate in the repeat generation was higher than in the initial generation in 11 tasks, lower in 13, and the same in 26. With the Omega Point method, the match outcome differed in eight of the fifty tasks; exhaustive enumeration of within-task permutations gives $p = 0.0625$ before correction and 0.188 after, while the smallest value attainable with this task composition is 0.0234. The confidence interval estimates the difference under task resampling, whereas the permutation test checks whether the two generations are exchangeable within each task.

The repeat-generation match rates include two sets that Opus refused to generate: the multistep pipeline for task R42664348 and the Omega Point method for task R42736449. In both, the classifier cut off the response at the hypothesis-generation stage in all six attempts over two runs (three per run), after which the model refused to try again. Without these sets, the overall match rate was 14.7% (36/245) with the multistep pipeline and 1.2% (3/245) with the Omega Point method. Retained sets also contained refusals at individual stages: 10 multistep-pipeline sets and 10 Omega Point sets in the initial generation, and 7 and 2 sets in the repeat generation; in these, the next stage received a model refusal or a truncated response instead of a result. In two single-request responses in the repeat generation, the model rewrote the complete set of hypotheses after a cutoff. In the repeat generation, the multistep pipeline differed from single-request generation by −3.6 percentage points (95% CI [−8.4, 1.2]; $p = 0.194$ after Holm correction across three within-model comparisons).

A separate analysis excluded every set in which the saved response of at least one stage contained a Claude refusal or a fragment truncated by the classifier: 10 multistep-pipeline sets and 10 Omega Point sets in the initial generation, and 8 multistep-pipeline sets, 3 Omega Point sets, and 2 single-request sets in the repeat generation, including both sets that failed on every attempt. Before the fix of 14 October, the truncated fragment was saved together with the retry response; a cutoff followed by a complete response leaves no trace in the saved text. The repeat-generation log records 103 message cutoffs by the classifier, including those in the excluded sets. Without the excluded sets, the match rate among the remaining hypotheses with the Omega Point method was 3.0% (6/200; 95% CI [0.6, 5.9]) in the initial generation and 1.3% (3/235; 95% CI [0.0, 2.9]) in the repeat generation; with the multistep pipeline it was 11.5% (23/200; 95% CI [4.6, 20.0]) and 15.2% (32/210; 95% CI [9.4, 21.4]). In tasks where both sets were retained, the match rate with the Omega Point method minus that with single-request generation was −16.5 percentage points (95% CI [−23.0, −10.5]; 40 tasks) in the initial generation and −17.8 percentage points (95% CI [−23.6, −12.4]; 45 tasks) in the repeat generation.

Leaving aside the initial Opus generations, the overall match rate of the other three models ranged from 13.2% to 16.8% with a single request, from 16.0% to 18.8% with the multistep pipeline, and from 2.8% to 7.6% with the Omega Point method. The difference between the multistep pipeline and single-request generation had $p > 0.05$ for all three models after Holm correction across nine comparisons.

#### M.2. Degree of cueing and generation

In the 38 tasks with a cueing score of 0 or 1, the overall match rate in single-request generation was 10.5% (20/190) for Astra, 13.2% (25/190) for Grok, 14.7% (28/190) for Opus, and 11.1% (21/190) for Gemini, against 14.4%, 16.8%, 18.0%, and 13.2% across all 50 tasks. In the Opus repeat generation, it was 13.7% (26/190) against 18.0%.

Under the prespecified alternative coding of borderline research questions, the correlation between the cueing score and the match rate was $\rho = -0.06$ (95% CI [−0.34, 0.23]; uncorrected $p = 0.703$). For the joint cueing score of the research question and the background, the correlation with the match rate at the primary threshold was $\rho = 0.29$ (95% CI [−0.06, 0.59]; uncorrected permutation $p = 0.040$). This exploratory finding, the between-group difference, and the recalculation on the 38 tasks are consistent with cueing in the prompt contributing to matches with the reference finding.

#### M.3. Comparison of generators on shared tasks

The comparison uses the 48 tasks common to the three sets, 240 hypotheses per generator. On these tasks, the final selected list contains 14 matching hypotheses of 19 for Grok, 9 of 19 for Gemini, and 13 of 20 for Opus. Two tasks in the full Gemini and Opus sets are excluded here, so the counts can differ from Table 6.

| Generators compared, first vs second | Match rate among selected hypotheses, % (n/N) | Difference in match rates, pp [95% CI] | Proportion of matching hypotheses found, % (n/N) | Difference in sensitivity, pp [95% CI] |
|---|---|---|---|---|
| Grok 4.5 vs Gemini 4 Argon | 73.7% (14/19) vs 47.4% (9/19) | 26.3 [−0.2, 52.6] | 35.0% (14/40) vs 28.1% (9/32) | 6.9 [−9.7, 24.0] |
| Grok 4.5 vs Claude Opus 5.5 | 73.7% (14/19) vs 65.0% (13/20) | 8.7 [−21.2, 37.3] | 35.0% (14/40) vs 29.5% (13/44) | 5.5 [−11.2, 21.2] |
| Gemini 4 Argon vs Claude Opus 5.5 | 47.4% (9/19) vs 65.0% (13/20) | −17.6 [−46.4, 11.0] | 28.1% (9/32) vs 29.5% (13/44) | −1.4 [−21.8, 18.4] |
| Claude Opus 5.5, initial vs repeat generation | 61.9% (13/21) vs 58.8% (10/17) | 3.1 [−24.8, 31.3] | 29.5% (13/44) vs 23.3% (10/43) | 6.3 [−7.6, 19.6] |

Each difference is the first value minus the second. CIs come from 10,000 paired resamples of the 48 tasks (for the last row, of the 49 tasks complete in both Opus sets). Every difference interval includes zero, so the comparison identifies no generator as better under this rule.

The random answers for the Opus repeat generation were recorded on 28 October at 22:44 UTC, before the hypothesis set was frozen at 22:45 and before the first request to the predictors at 22:45:57. All 213 requests to the four models passed the check that tools were disabled. For one task, Grok twice failed to return a forecast, and the task's five hypotheses were excluded as a technical failure. At the strict threshold, 2 of the 17 selected hypotheses matched the reference finding in the repeat generation, as did 2 of the 22 selected in the initial generation.

#### M.4. Number of forecasts

The unit is one forecast by one model for one hypothesis. Each of the four models forecast every hypothesis in the set, receiving the five hypotheses of one task per request.

| Generator | Hypotheses in the set | Forecasts by the four models |
|---|---|---|
| Grok 4.5 | 240 | 960 |
| Gemini 4 Argon | 250 | 1000 |
| Claude Opus 5.5 | 250 | 1000 |

#### M.5. Final selection under the two match thresholds

The selected lists are unchanged. The denominator of the match rate among selected hypotheses is the number of selected hypotheses; the denominator of sensitivity is the number of matching hypotheses in the full set at the same threshold. CIs come from 10,000 resamples of whole tasks.

| Generator | Score from a majority of judges | Match rate among selected hypotheses, % (n/N) [95% CI] | Selection sensitivity, % (n/N) [95% CI] |
|---|---|---|---|
| Grok 4.5 | at least 3 | 73.7% (14/19) [52.6, 93.3] | 35.0% (14/40) [25.0, 46.2] |
| Grok 4.5 | at least 4 | 26.3% (5/19) [7.1, 47.4] | 71.4% (5/7) [33.3, 100.0] |
| Gemini 4 Argon | at least 3 | 47.4% (9/19) [25.0, 70.6] | 27.3% (9/33) [13.6, 42.9] |
| Gemini 4 Argon | at least 4 | 31.6% (6/19) [11.1, 53.8] | 66.7% (6/9) [37.5, 100.0] |
| Claude Opus 5.5 | at least 3 | 63.6% (14/22) [42.9, 82.4] | 31.1% (14/45) [18.4, 43.9] |
| Claude Opus 5.5 | at least 4 | 9.1% (2/22) [0.0, 21.1] | 40.0% (2/5) [0.0, 100.0] |
| Claude Opus 5.5, repeat generation | at least 3 | 58.8% (10/17) [35.0, 81.8] | 23.3% (10/43) [12.5, 34.7] |
| Claude Opus 5.5, repeat generation | at least 4 | 11.8% (2/17) [0.0, 28.6] | 66.7% (2/3) [0.0, 100.0] |

#### M.6. Predictor panel recommendations and a random list of the same length

For the predictor panel, the table gives the observed result; for random selection, it gives the mean and the central 95% range of the number of matching hypotheses under the exact distribution (Appendix L.3). Because the distribution is discrete, the range can cover more than 95% of possible outcomes. Table 6 gives confidence intervals for the panel result.

| Generator | Predictor panel, matching / selected | Random selection, mean number matching | Random selection, central 95% range | Full set, matching / total |
|---|---|---|---|---|
| Grok 4.5 | 14/19 (73.7%) | 3.17 of 19 (16.7%) | 0 to 6 of 19 (0.0% to 31.6%) | 40/240 (16.7%) |
| Gemini 4 Argon | 9/19 (47.4%) | 2.51 of 19 (13.2%) | 0 to 6 of 19 (0.0% to 31.6%) | 33/250 (13.2%) |
| Claude Opus 5.5 | 14/22 (63.6%) | 3.96 of 22 (18.0%) | 1 to 8 of 22 (4.5% to 36.4%) | 45/250 (18.0%) |

When a single model's positive forecasts on the Grok hypotheses are randomly subsampled to 19, the upper bounds of the central ranges are 12 for Sonnet, Astra, and Gemini and 9 for Grok, whereas the predictor panel selected 14 matching hypotheses. On the Gemini hypotheses, the panel's 9 of 19 lies within the range of 3 to 10 obtained by subsampling Gemini's own 53 "yes" forecasts to 19 (18 of the 53 match). On the Opus hypotheses, the panel's 14 of 22 exceeds by one the upper bound of the range of 6 to 13 obtained by subsampling Astra's 49 "yes" forecasts to 22 (21 of the 49 match).

##### M.7. Robustness checks for the final selection

Of the 38 tasks with a cueing score of 0 or 1, the match rate among selected hypotheses was calculated on 36 tasks for Grok, 38 for Gemini and Opus, and 37 for the Opus repeat generation; the 95% CIs were [33.3, 90.9], [11.1, 62.5], [38.5, 84.2], and [22.2, 85.7]. In the full sets for the same tasks, 12.8% (23/180), 11.1% (21/190), 14.7% (28/190), and 13.0% (24/185) of hypotheses matched the reference finding.

With judges left out one at a time, the largest drop was for the Opus hypotheses without the xAI judge: from 63.6% to 54.5% (12/22). Leaving out the judge from Google, the developer of the Gemini predictor, did not change the result for the Grok hypotheses and removed one match among the selected Opus hypotheses in both generations. The ratio of the match rate among selected hypotheses to the baseline match rate ranged from 3.5 to 4.8. With two judges left, the match rule is stricter, so part of the drop reflects the change of rule.

Without the predictor from the generator's own developer, the 95% CIs for the match rate among selected hypotheses were [47.6, 86.7] for Grok, [23.1, 66.7] for Gemini, [35.0, 70.8] for Opus, and [26.1, 65.6] for the Opus repeat generation. The proportion of matching hypotheses found rose by 0.0 to 6.9 percentage points. Astra stayed in the panel in every set, because its developer built none of the three generators. In the repeat forecast run, the CIs for the match rate among selected hypotheses were [46.7, 88.9] for Grok, [26.7, 75.0] for Gemini, and [43.5, 86.4] for Opus; 235, 245, and 240 hypotheses received forecasts in both runs. The odds ratio per one-point increase in clarity score was 2.02 (95% CI [1.04, 3.95]; $p < 0.05$), and adding feasibility to the regression left the estimates almost unchanged. The mean originality score was 4.86 for selected and 5.08 for excluded hypotheses; one Opus task had no excluded hypothesis left, so the comparison covers 59 pairs. By the lexical criterion, matching hypotheses named on average 13.0% of key elements and the remaining hypotheses 4.1%.

##### M.8. The "effect in the favourable direction" rule on the 50/50 set

Among the 61 labelled statements, 35 of the 38 confirmed and 18 of the 23 refuted statements had a favourable direction. Model forecasts agreed with the rule on 50.8% to 62.3% of labelled statements. On the 39 unlabelled statements, the models were correct in 61.5% to 82.1% of cases.

#### Appendix N. Contents of the published data

Reports of the forecasting runs of 22 to 24 October are published at `https://eternalsearch.net/omega/<directory>/report.json`, with the logs and saved requests in the same directories: `experimental-support-20261022`, `direct-confirmation-20261022`, `direct-confirmation-grok-20261022`, `balanced-confirmation-20261022`, `balanced-opus-20261023`, `grok-generated-confirmation-20261023`, `gemini-generated-confirmation-20261023`, `opus-generated-confirmation-20261023`, and `selection-replication-20261023`. The final selected lists are in the `selected.csv` files in the `two-stage-unanimous-20261023`, `two-stage-unanimous-gemini-20261023`, and `two-stage-unanimous-opus-20261023` directories.

Forecasts and selection results for the hypotheses of the Opus repeat generation are in the `opus-isolated-confirmation-20261028`, `two-stage-unanimous-opus-isolated-20261028`, and `opus-isolation-selection-20261028` directories. The repeat forecast run, with verification of each request's isolation, is in `forecast-repeat-20261030-v2/comparison.json`, and the originality scores of selected and excluded hypotheses are in the `selection-originality-20261030` directory. The Opus repeat-generation logs verifying each request's isolation, the records of Claude refusals with full model responses, the reproduction of the earlier configuration, and the recalculations on the 38 tasks and without each developer's judge are available on request.

The data description sets out the structure of the row-level data and the difference between the outcome `y` and the decision `selected`. In `analysis.json`, the `selected` field is computed from the four actual forecasts. The unused `yes` field, which wrongly contained `false` in the historical reports, has been removed from the row-level data of this edition; the archived reports are kept in their original form. The data release keeps the checksums of the judge scores.

#### Appendix O. Control rules and robustness checks

These analyses use the published files and the saved model responses; three of them (O.7, O.9, and O.14) also used new model responses. All were added after the results were known. Intervals are 95% percentile intervals over resamples of tasks (of articles in O.11).

##### O.1. Matches by position in the generator's response

| Generator | Position 1 | Position 2 | Position 3 | Position 4 | Position 5 |
|---|---|---|---|---|---|
| GPT-6 Astra | 36.0% (18/50) | 16.0% (8/50) | 10.0% (5/50) | 8.0% (4/50) | 2.0% (1/50) |
| Grok 4.5 | 36.0% (18/50) | 20.0% (10/50) | 8.0% (4/50) | 12.0% (6/50) | 8.0% (4/50) |
| Claude Opus 5.5 | 38.0% (19/50) | 20.0% (10/50) | 12.0% (6/50) | 8.0% (4/50) | 12.0% (6/50) |
| Gemini 4 Argon | 40.0% (20/50) | 12.0% (6/50) | 4.0% (2/50) | 10.0% (5/50) | 0.0% (0/50) |
| Four generators | 37.5% (75/200) | 17.0% (34/200) | 8.5% (17/200) | 9.5% (19/200) | 5.5% (11/200) |
| Four generators, strict threshold | 10.0% (20/200) | 1.0% (2/200) | 0.5% (1/200) | 0.5% (1/200) | 0.5% (1/200) |

Single request, 50 tasks. In 75 of the 112 generator-task pairs with at least one match, the first-listed hypothesis matched; allocating the matches to positions at random within each pair gives an expected 31.2 (exact test, $p < 0.001$). Judges saw the five hypotheses of a set in a pseudo-random order fixed for the task and generation method; the proportion of votes of 3 or higher by displayed position was 16.8% (101/600), 18.3% (110/600), 13.7% (82/600), 14.0% (84/600), and 18.5% (111/600) for positions 1 to 5 (the three judges from other developers). The first-listed hypothesis was displayed first in 60 of 200 sets.

##### O.2. Convergence of hypotheses across generators

| Generators in the group | Hypotheses | Match rate [95% CI] |
|---|---|---|
| 1 | 330 | 7.6% (25/330) [4.8, 10.7] |
| 2 | 78 | 6.4% (5/78) [0.0, 13.8] |
| 3 | 52 | 11.5% (6/52) [2.2, 22.2] |
| 4 | 540 | 22.2% (120/540) [15.9, 29.0] |

Single request, 1000 hypotheses. Within each task, groups are connected components of hypotheses whose statement embeddings (embeddinggemma 300m, 768 dimensions) have a cosine similarity of at least 0.75, computed across the four generators of this study. The match rate of groups shared by all four generators exceeded that of hypotheses without a counterpart by 14.6 pp [8.2, 21.3]. The mean number of generators in a hypothesis's group was 3.53, 2.94, 2.74, 2.52, and 2.28 for positions 1 to 5. Per task, the number of generators with at least one matching hypothesis was none in 10 tasks, one in 7, two in 12, three in 3, and four in 18.

##### O.3. The first-listed hypothesis as a control rule

| Set | Unanimity, PPV | First-listed, PPV [95% CI] | PPV difference, pp [95% CI] | Sensitivity difference, pp [95% CI] |
|---|---|---|---|---|
| Grok 4.5 | 73.7% (14/19) | 37.5% (18/48) [25.0, 52.1] | 36.2 [19.2, 54.5] | −10.0 [−25.6, 2.9] |
| Gemini 4 Argon | 47.4% (9/19) | 40.0% (20/50) [26.0, 54.0] | 7.4 [−12.7, 28.0] | −33.3 [−52.6, −15.4] |
| Claude Opus 5.5 | 63.6% (14/22) | 38.0% (19/50) [24.0, 52.0] | 25.6 [9.7, 43.0] | −11.1 [−24.4, 1.8] |
| Three sets | 61.7% (37/60) | 38.5% (57/148) [27.2, 50.0] | 23.2 [13.0, 32.6] | −16.9 [−27.5, −7.8] |
| Claude Opus 5.5, repeat generation | 58.8% (10/17) | 30.6% (15/49) [18.4, 42.9] | 28.2 [9.2, 48.6] | −11.6 |

| Model | OR of selection [95% CI] | OR of first position [95% CI] |
|---|---|---|
| Selection | 9.01 [4.70, 21.21] | — |
| Selection and first position | 3.30 [1.54, 8.06] | 4.08 [2.09, 8.52] |

Differences are unanimity minus first-listed hypotheses on the same resamples. Selected hypotheses by position in the three sets: 50, 7, 1, 2, and 0 for positions 1 to 5; 6 of the 61 matching hypotheses in positions 2 to 5 were selected. The second table is a conditional logistic regression with each generator's tasks as strata (147 strata, 85 of them informative). The three generators answered the same tasks, so the resampling unit is the task with all of its strata; intervals come from 2000 resamples of tasks.

##### O.4. Selection-adjusted estimate for the procedure that picks a rule

| Rule (members, minimum number of "yes") | List length | Matches | PPV, % | Sensitivity, % |
|---|---|---|---|---|
| Sonnet, Astra, Gemini, Grok, Opus ≥ 5 | 16 | 12 | 75.0 | 30.0 |
| Sonnet, Astra, Gemini, Grok ≥ 4 | 19 | 14 | 73.7 | 35.0 |
| Sonnet, Astra, Grok, Opus ≥ 4 | 18 | 13 | 72.2 | 32.5 |
| Sonnet, Astra, Grok ≥ 3 | 23 | 16 | 69.6 | 40.0 |
| Sonnet, Astra, Gemini, Opus ≥ 4 | 19 | 13 | 68.4 | 32.5 |
| Sonnet, Astra, Gemini ≥ 3 | 22 | 15 | 68.2 | 37.5 |

| Criterion for choosing a variant | Apparent PPV of the best variant, % | Leave-one-task-out PPV | Optimism-corrected PPV, % |
|---|---|---|---|
| highest PPV, list of at least 10 | 75.0 | 66.7% (12/18) | 69.7 |
| highest PPV, sensitivity of at least 30% | 75.0 | 66.7% (12/18) | 69.8 |
| highest F1 | 56.1 | 47.7% (21/44) | 50.9 |

Grok hypotheses; 112 variants yield 87 distinct lists. The chosen rule (unanimity of Sonnet, Astra, Gemini, and Grok) ranks 2nd by PPV among variants with lists of at least 10, after unanimity of all five models. The three right-hand columns describe the procedure "take the variant that this criterion ranks first", not the chosen rule: in 46 of the 48 folds the criterion took unanimity of all five models, in 1 fold the chosen rule, and in 1 fold unanimity of Sonnet, Astra, Grok, and Opus. Leave one task out: for each of the 48 tasks, the variant is chosen by the criterion on the other 47 tasks and applied to the held-out task. Optimism correction: the mean difference between apparent and out-of-sample PPV over 2000 bootstrap resamples, with the selection repeated in each. For the chosen rule itself, the check is its application to the Gemini and Opus hypotheses, fixed on 23 October before those forecasts were obtained (Section 3.5 and Appendix L.4).

##### O.5. Random subsets of the best single model and ranking by votes

| Set | Unanimity list | Matches | Best single model | Its "yes" forecasts | Matches among them | Expected in a random subset | *p* of at least as many |
|---|---|---|---|---|---|---|---|
| Grok 4.5 | 19 | 14 | GPT-6 Astra | 53 | 25 | 8.96 | 0.0043 |
| Gemini 4 Argon | 19 | 9 | Gemini 4 Argon | 53 | 18 | 6.45 | 0.1084 |
| Claude Opus 5.5 | 22 | 14 | GPT-6 Astra | 49 | 21 | 9.43 | 0.0087 |
| Claude Opus 5.5, repeat generation | 17 | 10 | Gemini 4 Argon | 61 | 21 | 5.85 | 0.0152 |

| List length | By votes | By votes, then listing order | By listing order | Random list |
|---|---|---|---|---|
| 30 | 18.5 | 18.6 | 11.6 | 4.8 |
| 60 | 37.0 | 37.0 | 23.1 | 9.6 |
| 75 | 41.9 | 42.7 | 28.9 | 12.0 |
| 109 | 53.0 | 53.0 | 42.0 | 17.4 |
| 148 | 59.6 | 61.5 | 57.0 | 23.6 |
| 150 | 60.0 | 61.9 | 57.3 | 23.9 |
| 300 | 80.4 | 81.2 | 82.3 | 47.8 |

Three sets together, 740 hypotheses with 118 matches; cells give the expected number of matches in a list of the given length, with ties broken at random. The combined unanimity lists have length 60, and the first-listed hypotheses number 148.

| Set | All | At least 1 "yes" | At least 2 | At least 3 | All four |
|---|---|---|---|---|---|
| Grok 4.5 | 240 / 40 | 132 / 37 | 73 / 30 | 37 / 19 | 19 / 14 |
| Gemini 4 Argon | 250 / 33 | 130 / 22 | 72 / 18 | 33 / 15 | 19 / 9 |
| Claude Opus 5.5 | 250 / 45 | 134 / 32 | 70 / 23 | 39 / 19 | 22 / 14 |
| Three sets | 740 / 118 | 396 / 91 | 215 / 71 | 109 / 53 | 60 / 37 |

Each cell gives hypotheses in the list / matching hypotheses (Figure 3).

##### O.6. Judge panels, leniency, and self-preference

| Judge | Votes ≥ 3, % | Votes ≥ 4, % | Mean score | Generator from its developer | Difference in differences, points [95% CI] | Same for votes ≥ 3, pp [95% CI] |
|---|---|---|---|---|---|---|
| GPT-5.6 Sol | 16.0 | 2.2 | 1.99 | GPT-6 Astra | 0.13 [0.07, 0.18] | 0.6 [−1.9, 3.3] |
| Grok 4.7 | 12.9 | 1.5 | 2.04 | Grok 4.5 | 0.00 [−0.04, 0.05] | 2.2 [0.6, 4.0] |
| Claude Opus 5.5 | 12.1 | 4.1 | 1.32 | Claude Opus 5.5 | 0.03 [−0.03, 0.10] | 0.3 [−1.3, 2.1] |
| Gemini 4 Argon (medium) | 11.7 | 2.0 | 1.66 | Gemini 4 Argon | 0.15 [0.10, 0.20] | 0.9 [−1.4, 3.2] |

All 2997 hypotheses. Difference in differences: the judge's score minus the other judges' scores on hypotheses from its own developer's generator, compared with the same difference on the other generators' hypotheses.

| Judges deciding the outcome | GPT-6 Astra | Grok 4.5 | Claude Opus 5.5 | Gemini 4 Argon |
|---|---|---|---|---|
| Primary rule (judge from the generator's developer excluded) | 36 | 42 | 45 | 33 |
| Sol, Grok 4.7, Opus | 38 | 47 | 44 | 33 |
| Sol, Grok 4.7, Gemini | 36 | 42 | 45 | 35 |
| Sol, Opus, Gemini | 38 | 42 | 41 | 29 |
| Grok 4.7, Opus, Gemini | 36 | 45 | 42 | 31 |
| All four judges, at least 2 votes | 39 | 47 | 48 | 35 |
| All four judges, at least 3 votes | 35 | 41 | 38 | 29 |

Matching hypotheses out of 250 in single-request generation.

#### O.7. Composition of the predictor panel

| Panel | Grok 4.5 | Gemini 4 Argon | Claude Opus 5.5 | Claude Opus 5.5, repeat generation |
|---|---|---|---|---|
| Four models (primary rule) | 73.7% (14/19) | 47.4% (9/19) | 63.6% (14/22) | 58.8% (10/17) |
| Without Claude Sonnet 5.5 | 61.5% (16/26) | 50.0% (14/28) | 53.6% (15/28) | 46.4% (13/28) |
| Without GPT-6 Astra | 60.9% (14/23) | 43.5% (10/23) | 57.1% (16/28) | 50.0% (10/20) |
| Without Gemini 4 Argon | 69.6% (16/23) | 45.0% (9/20) | 60.0% (15/25) | 43.5% (10/23) |
| Without Grok 4.5 | 68.2% (15/22) | 47.4% (9/19) | 62.5% (15/24) | 55.0% (11/20) |
| Gemini 3.8 Flash in place of Gemini 4 Argon | 73.3% (11/15) | 50.0% (8/16) | 57.9% (11/19) | 46.7% (7/15) |

Cells give the PPV of the list selected unanimously by the panel. The predictor from the generator's developer is Grok 4.5 for the Grok hypotheses, Gemini 4 Argon for the Gemini hypotheses, and Claude Sonnet 5.5 for the Opus hypotheses. Gemini 3.8 Flash forecasts were obtained under a protocol fixed before the first request; this model was not a judge in this study.

#### O.8. Dependence of predictor errors

| Set | Unanimous "yes" for non-matching hypotheses | Expected under independence | Ratio [95% CI] | PPV under independence, % | Observed PPV |
|---|---|---|---|---|---|
| Grok 4.5 | 5 | 0.20 | 24.9 [6.3, 59.7] | 96.7 | 73.7% (14/19) |
| Gemini 4 Argon | 10 | 0.36 | 28.1 [15.4, 50.6] | 82.6 | 47.4% (9/19) |
| Claude Opus 5.5 | 8 | 0.29 | 27.8 [13.9, 52.5] | 89.7 | 63.6% (14/22) |
| Three sets | 23 | 0.84 | 27.5 [18.3, 40.7] | 91.7 | 61.7% (37/60) |

| Models required to say "yes" | Combinations | Mean list length | Mean PPV, % | Mean sensitivity, % |
|---|---|---|---|---|
| 1 | 4 | 195.0 | 34.7 | 53.4 |
| 2 | 6 | 102.2 | 47.7 | 40.7 |
| 3 | 4 | 72.2 | 56.9 | 34.7 |
| 4 | 1 | 60.0 | 61.7 | 31.4 |

Independence given the outcome: the expected number of unanimous "yes" forecasts is the product of the four models' "yes" rates among non-matching (or matching) hypotheses, multiplied by the number of such hypotheses. Second table: unanimity within every combination of the given number of the four models, three sets together.

#### O.9. Reference findings extracted by Gemini 4 Argon and re-scoring

| Set | Tasks | Matches, Sol / Gemini reference | Unanimity PPV, Sol / Gemini reference | Gain over first-listed, pp [95% CI], Sol / Gemini |
|---|---|---|---|---|
| Grok 4.5 | 45 | 37 / 32 | 76.5% (13/17) / 70.6% (12/17) | 38.7 [19.6, 57.8] / 37.3 [17.8, 57.8] |
| Gemini 4 Argon | 47 | 29 / 34 | 44.4% (8/18) / 44.4% (8/18) | 4.0 [−16.9, 25.3] / 6.1 [−13.4, 26.1] |
| Claude Opus 5.5 | 47 | 44 / 40 | 63.6% (14/22) / 63.6% (14/22) | 25.3 [9.6, 42.1] / 29.6 [9.9, 48.1] |
| Three sets | 139 | 110 / 106 | 61.4% (35/57) / 59.6% (34/57) | 22.6 [11.8, 33.6] / 24.4 [13.2, 36.0] |

Gemini 4 Argon at high reasoning effort received the original extraction request. It accepted 47 of the 50 tasks, judging R42642518, R42647635, and R42696595 unsuitable, and for all 47 Grok 4.7 judged the core finding, the mechanism or target, and the direction of the effect to be the same as in GPT-5.6 Sol's version. The three judges from other developers re-scored the single-request hypotheses of the three selection sets against Gemini's reference findings with the original request, order, and settings; the original request, rebuilt by the same script, matched the logged request byte for byte. Votes of 3 or higher against the new and the original reference findings agreed in 95.4% of cases (1989/2085): GPT-5.6 Sol 94.4% (656/695), Grok 4.7 96.6% (454/470), Claude Opus 5.5 94.6% (435/460), and Gemini 4 Argon (medium) 96.5% (444/460). Outcomes agreed in 95.7% (665/695). The Sol-reference values are recalculated on the same tasks.

#### O.10. Calibration of the probability of confirmation

| Predictor | Hypotheses | Mean probability | Match rate | Brier score | Skill score [95% CI] | Calibration slope | Match rate at probability 0–0.2 | 0.2–0.4 | 0.4–0.6 | 0.6–1.0 |
|---|---|---|---|---|---|---|---|---|---|---|
| Claude Sonnet 5.5 | 250 | 0.24 | 0.14 | 0.123 | 0.01 [−0.21, 0.13] | 1.14 | 6.3% (5/79) | 12.9% (17/132) | 35.3% (12/34) | 40.0% (2/5) |
| Claude Opus 5.5 | 215 | 0.41 | 0.14 | 0.187 | −0.60 [−1.54, −0.17] | 1.09 | 0.0% (0/10) | 6.5% (6/93) | 17.5% (14/80) | 28.1% (9/32) |
| Claude Haiku 5.5 | 250 | 0.26 | 0.14 | 0.135 | −0.10 [−0.33, 0.03] | 0.77 | 10.8% (7/65) | 13.5% (21/155) | 23.1% (6/26) | 50.0% (2/4) |
| Claude Opus 4.8 | 240 | 0.43 | 0.14 | 0.197 | −0.62 [−1.34, −0.24] | 1.46 | 0.0% (0/2) | 7.5% (6/80) | 15.6% (22/141) | 35.3% (6/17) |
| GPT-6 Astra | 245 | 0.47 | 0.14 | 0.228 | −0.86 [−1.72, −0.39] | 0.99 | 20.0% (1/5) | 5.5% (4/73) | 12.0% (14/117) | 32.0% (16/50) |
| Gemini 4 Argon | 250 | 0.51 | 0.14 | 0.250 | −1.03 [−2.03, −0.48] | 1.15 | 0.0% (0/7) | 6.8% (5/74) | 7.6% (6/79) | 27.8% (25/90) |
| Grok 4.5 | 225 | 0.40 | 0.14 | 0.180 | −0.52 [−1.19, −0.18] | 1.25 | 0.0% (0/5) | 6.2% (7/112) | 19.1% (17/89) | 36.8% (7/19) |

The 250 GPT-6 Astra single-request hypotheses with a numerical probability under the detailed prompt. Skill score: 1 minus the ratio of the Brier score to that of a constant forecast equal to the match rate; negative values mean a larger error than the constant forecast. Slope: logistic regression of a match on the logit of the probability. The probability refers to experimental confirmation, whereas the outcome counts a correct hypothesis about a mechanism other than the article's as a non-match.

#### O.11. The 50/50 set by publication date

| Model | Stated training cutoff | Balanced accuracy, 25 articles before 1 September, % | Balanced accuracy, 25 articles from 1 September, % | Difference, pp [95% CI] |
|---|---|---|---|---|
| Claude Sonnet 5.5 | 2026-06-30 | 72.0 | 76.0 | 4.0 [−12.0, 20.0] |
| GPT-6 Astra | 2026-04-30 | 76.0 | 66.0 | −10.0 [−26.0, 6.0] |
| Gemini 4 Argon | 2026-03-31 | 82.0 | 74.0 | −8.0 [−24.0, 8.0] |
| Grok 4.5 | 2026-01-31 | 80.0 | 70.0 | −10.0 [−28.0, 8.0] |
| Claude Opus 5.5 | 2026-06-30 | 78.0 | 68.0 | −10.0 [−24.0, 6.0] |
| Five-model unanimity | — | 74.0 | 66.0 | −8.0 [−22.0, 6.0] |

Mean change for Opus and Sonnet minus mean change for Astra, Gemini, and Grok: 6.3 pp [−5.0, 18.0]. If training-data leakage contributed to performance on the earlier articles, this value would be negative. Intervals come from 2000 resamples of articles within each period.

#### O.12. Second random sequence for the Astra hypotheses

| Threshold | Match rate | PPV | NPV | Sensitivity | Specificity | PPV range across fair coins |
|---|---|---|---|---|---|---|
| Primary threshold | 14.4% (36/250) | 13.0% (17/131) | 84.0% (100/119) | 47.2% (17/36) | 46.7% (100/214) | [10.0, 18.8] |
| Strict threshold | 1.6% (4/250) | 0.8% (1/131) | 97.5% (116/119) | 25.0% (1/4) | 47.2% (116/246) | [0.0, 3.2] |

Seed 20261104 was fixed in a protocol file committed before the calculation; the sequence has 131 "yes" answers among 250. The range covers the central 95% of 10,000 further fair-coin sequences on the same hypotheses.

#### O.13. Selection adjusted for lexical overlap, length, mechanisms, and entities

| Model | Unanimous selection | Named key elements, per 10 pp | Length, per 100 words | Named mechanisms, per one | Named entities, per one |
|---|---|---|---|---|---|
| Selection | 9.01 [4.70, 21.21] | — | — | — | — |
| Selection and key elements | 7.55 [3.98, 17.86] | 1.90 [1.53, 2.86] | — | — | — |
| All covariates | 5.47 [2.75, 13.14] | 1.84 [1.47, 2.86] | 13.51 [1.17, 253.12] | 0.85 [0.39, 1.63] | 1.01 [0.91, 1.13] |

Odds ratios [95% CI] from conditional logistic regression on the three sets, with each generator's tasks as strata and the task as the resampling unit. Named key elements follow the rule of Appendix K.8. Gemini 4 Argon counted mechanisms and entities in the 1000 single-request hypotheses. Selected versus other hypotheses: mean share of named key elements 0.130 and 0.051, mean length 140.5 and 133.1 words, mechanisms 1.20 and 1.35, and entities 12.60 and 12.77.

#### O.14. Blind re-coding of the question cue

| Codings | Same score | Cohen's kappa | Quadratic-weighted kappa |
|---|---|---|---|
| Original and Gemini 4 Argon | 39/50 | 0.59 | 0.71 |
| Original and Grok 4.7 | 34/50 | 0.47 | 0.65 |
| Gemini 4 Argon and Grok 4.7 | 43/50 | 0.75 | 0.81 |

| Coding | Questions with score 2 | Match rate, score 2 | Match rate, scores 0–1 | Difference, pp [95% CI] | Spearman's rho [95% CI] |
|---|---|---|---|---|---|
| Original | 12 | 25.8% (62/240) | 12.4% (94/760) | 13.5 [2.1, 25.2] | 0.17 [−0.14, 0.45] |
| Gemini 4 Argon | 5 | 35.0% (35/100) | 13.4% (121/900) | 21.6 [−0.9, 38.0] | 0.18 [−0.17, 0.47] |
| Grok 4.7 | 6 | 37.5% (45/120) | 12.6% (111/880) | 24.9 [5.5, 38.8] | 0.23 [−0.10, 0.51] |
| Score 2 from both re-coders | 5 | 35.0% (35/100) | 13.4% (121/900) | 21.6 [−0.9, 38.0] | 0.30 [−0.11, 0.57] |

Single request, four generators. The re-coders received the frozen rubric, the background, the question, the reference finding, and its key elements, but no outcome. Intervals come from 10,000 resamples of tasks (2000 for rho); the intervals of the original coding therefore differ slightly from those of the primary analysis in Section 3.1.

#### O.15. Difference in the gain from selection between generators

| Generators compared | Shared tasks | Gain of each, pp | Difference, pp [95% CI] | *p* | Holm-adjusted *p* |
|---|---|---|---|---|---|
| Grok 4.5 and Gemini 4 Argon | 48 | 36.2 and 7.8 | 28.4 [1.4, 56.1] | 0.053 | 0.16 |
| Claude Opus 5.5 and Gemini 4 Argon | 50 | 25.6 and 7.4 | 18.3 [−7.8, 44.6] | 0.17 | 0.34 |
| Grok 4.5 and Claude Opus 5.5 | 48 | 36.2 and 27.5 | 8.7 [−18.5, 34.6] | 0.54 | 0.54 |

The gain is the match rate among selected hypotheses minus the match rate among first-listed hypotheses of the same set, computed on the tasks the two generators share. Under the null hypothesis, the two generators' whole hypothesis sets exchange labels within each task, the procedure that compares generation methods in Section 2.10; 19,999 permutations, with Holm correction across the three pairs. Intervals come from 10,000 paired resamples of the shared tasks.

##### O.16. Selection without the tasks whose reference findings the audit scored below 3

| Tasks kept | Number of tasks | Unanimity, PPV | First-listed, PPV | Sensitivity | Gain, pp [95% CI] |
|---|---|---|---|---|---|
| All tasks | 50 | 61.7% (37/60) | 38.5% (57/148) | 31.4% (37/118) | 23.2 [13.0, 32.6] |
| Reference finding scored 3 or not audited | 38 | 63.0% (29/46) | 38.4% (43/112) | 33.7% (29/86) | 24.7 [13.0, 35.7] |
| Reference finding scored 1 or 2 | 12 | 57.1% (8/14) | 38.9% (14/36) | 25.0% (8/32) | 18.3 [0.0, 35.1] |

Three sets together. Of the 20 audited reference findings, 8 scored 3, 11 scored 2, and 1 scored 1 on the accuracy scale of Appendix P.3; the reference findings of the remaining 30 tasks were not in the frozen sample. The middle row therefore keeps the tasks whose reference finding the audit confirmed in full together with the tasks it did not cover.

##### O.17. Scores behind the matches of the selected lists

| Set | Selected | Matches | No score above 3 | Matches at the strict threshold | Majority of two votes |
|---|---|---|---|---|---|
| Grok 4.5 | 19 | 14 | 4 | 5 | 1 |
| Gemini 4 Argon | 19 | 9 | 1 | 6 | 0 |
| Claude Opus 5.5 | 22 | 14 | 11 | 2 | 2 |
| Three sets | 60 | 37 | 16 | 13 | 3 |

Scores of the three judges from developers other than the generator's. A match with no score above 3 agrees with the mechanism or target of the reference finding while at least one of its key elements is missing or replaced, the level the primary threshold accepts (Section 2.4). The last column counts the matches whose majority one changed vote would reverse.

#### Appendix P. Manual audit of hypotheses and reference findings

##### P.1. Frozen samples and scoring

On 5 November 2026, we completed the audit of both frozen samples; all scores were given by one of us. The hypothesis sample contains 100 of the 1000 single-request hypotheses from the four generators, drawn from 44 of the 50 tasks. The rater saw the background, question, reference finding, key elements and hypothesis, and scored agreement with the reference finding on the judges' 0 to 5 scale. Generator identity, original position, judge scores, selection status and sampling stratum were hidden. The sample was frozen on 2 November, before the first score (SHA-256 `2c2c1cf5fc1c820c4cdbf9b5c7d27ed1b2dbb53f8dd602899b1acfea44c8f18e`).

The table gives stratum sizes and results at the primary threshold. "Unselected" means not on the 60-item list of the three selection sets. The 40 split-vote records are unselected records sampled specifically for a split vote. Population estimates use inverse-inclusion weights: the population size of the stratum divided by its sample size.

| Stratum | Population | Sample | Matches, manual audit | Matches, judges | Same binary decision |
|---|---|---|---|---|---|
| Selected by unanimity | 60 | 30 | 19/30 | 20/30 | 25/30 |
| Unselected, two positive judge votes | 36 | 20 | 15/20 | 20/20 | 15/20 |
| Unselected, one positive judge vote | 56 | 20 | 4/20 | 0/20 | 16/20 |
| Unselected, three positive judge votes | 83 | 15 | 11/15 | 15/15 | 11/15 |
| Unselected, zero positive judge votes | 765 | 15 | 1/15 | 0/15 | 14/15 |
| Total | 1000 | 100 | 50/100 | 55/100 | 81/100 |

The reference sample comprises the first 20 of the 50 tasks ordered by SHA-256 of `benchmark-reference-check-v1:` followed by the task identifier. It was frozen on 4 November, before the first check (SHA-256 `05f6a81400aae45ad2c5bc8b9b5f25a669c3bb032701a630fa315ce553ca69eb`). We checked 16 reference findings against full texts; reference findings nos. 1 (R42601407), 4 (R42821663), 11 (R42759510) and 20 (R42822442) were checked against abstracts, and their score of 3 means agreement with those abstracts. The scale distinguishes a correct finding with all key elements (3), a correct central finding with an imperfect key element (2), a substantive discrepancy in the central finding (1), and an incorrect reference (0).

#### P.2. Agreement with the judges

The comparator is the majority decision of the three judges from other developers, which is equivalent to thresholding their median score. At the primary threshold, both we and the judges scored 43 hypotheses as matches and 38 as non-matches; the judges alone scored 12 as matches, and we alone scored 7. At the strict threshold, the corresponding counts were 8, 84, 2, and 6.

| Threshold | Agreement [95% CI], % | Cohen's kappa [95% CI] | Matches, manual audit / judges |
|---|---|---|---|
| Primary, score ≥ 3 | 81.0 [73.4, 88.2] | 0.62 [0.46, 0.76] | 50/100 / 55/100 |
| Strict, score ≥ 4 | 92.0 [86.9, 96.6] | 0.62 [0.24, 0.84] | 14/100 / 10/100 |

The manual score equalled the median judge score in 62/100 cases and differed from it by at most one point in 99/100; quadratic-weighted kappa was 0.73. Reweighting the five strata to their population sizes gave an estimated agreement at the primary threshold of 89.7% [77.1, 97.0] and a kappa of 0.64 [0.37, 0.87]. These are weighted estimates; the unweighted percentages describe the enriched validation sample.

Among the 30 sampled selected hypotheses, 19 matched by the manual score (63.3%; 95% CI [41.7, 81.8]) and 20 by the judges' decision (66.7%); the paired difference was −3.3 pp [−19.0, 10.7]. At the strict threshold, 11/30 matched by the manual score and 8/30 by the judges' decision. Intervals come from 10,000 bootstrap resamples of the 44 sampled tasks, keeping each task's hypotheses together and the original sampling weights fixed (seed 20261105). Agreement with the manual audit checks the judges' decisions on matching the reference finding; the outcome is still defined as correspondence with the published finding.

#### P.3. Accuracy of reference findings

Across the reviewed sources, eight reference findings scored 3 (40.0%, 8/20), eleven scored 2 (55.0%, 11/20), and one scored 1 (5.0%, 1/20); the central finding was rated correct in 15/16 full-text checks and 4/4 abstract checks. Four of the eight reference findings scoring 3 were checked against abstracts. The one scoring 1 was no. 9, R42844664, on time-restricted feeding, microbial acetate and cardiovascular ageing (doi:10.1111/acel.70728). The primary analysis keeps the original reference findings and outcomes; excluding that whole task is a sensitivity analysis.

| Source reviewed | Reference findings | Score 3 | Score 2 | Score 1 | Score 0 |
|---|---|---|---|---|---|
| Full text | 16 | 4 | 11 | 1 | 0 |
| Abstract | 4 | 4 | 0 | 0 | 0 |
| Total | 20 | 8 | 11 | 1 | 0 |

Section 3.8 of the article explains how the four abstract-only checks are interpreted; our explanation of the score of 1 for R42844664 follows.

A score of 1 was assigned to reference finding no. 9 (R42844664, Aging Cell), which presents the sequence "acetate — GPR43 — HDAC3 inhibition — H4K8 acetylation — suppression of NLRP3-dependent pyroptosis" as an established mechanism. For GPR43 and HDAC3, the study reports expression levels only (Figs. 6O and S14); the study's authors performed no loss-of-function experiments using gene knockout, receptor antagonists, or enzyme inhibitors, and the acetate-associated change in HDAC3 in the HUVEC experiments is reported in opposite directions in two places in the article (Fig. S11). The link between H4K8 acetylation and NLRP3 remains correlational, because chromatin immunoprecipitation at the NLRP3 promoter was not performed; the authors acknowledge in the discussion that this experiment is needed.

For the microbiota as a whole, the description "necessary and sufficient" is supported: antibiotic treatment abolishes protection, and faecal microbiota transfer reproduces it. For acetate, only sufficiency is shown (sodium acetate gavage, Figs. 6A–P), because neither acetate blockade nor an acetate-deficient donor was used. Butyric, valeric, and isobutyric acids also rose significantly together with acetate, so the data do not single out acetate. The between-group difference in Limosilactobacillus is significant in the metagenomic data but not in the 16S data (Fig. S13), and the bacterium was not given to the animals as a probiotic; "acetate-producing Limosilactobacillus" should therefore be read as the authors' interpretation.

The receptor and deacetylase steps of this sequence, and the step from histone acetylation to pyroptosis, therefore lack causal support, which determined the score.

An abstract-only check establishes agreement with the abstract from which the reference finding was compiled. The source of every reference finding and the full explanation accompany the item-level data.

Without task R42844664, the selected lists contained 14 matches among 18 Grok hypotheses (77.8%), 9 among 19 Gemini hypotheses (47.4%), and 13 among 21 Opus hypotheses (61.9%); pooled, 36/58 (62.1%), against 37/60 (61.7%) across all tasks. In the manual-audit sample, agreement at the primary threshold was 81/99, and both we and the judges scored 19 of the 29 remaining selected hypotheses as matches. Item-level scores, inputs and calculation results are published together with the code of this analysis.
